# SPIRED-Fitness: an end-to-end framework for the prediction of protein structure and fitness from single sequence

**DOI:** 10.1101/2024.01.31.578102

**Authors:** Yinghui Chen, Yunxin Xu, Di Liu, Yaoguang Xing, Haipeng Gong

**Affiliations:** MOE Key Laboratory of Bioinformatics, School of Life Sciences, Tsinghua University, Beijing, 100084, China; Beijing Frontier Research Center for Biological Structure, Tsinghua University, Beijing, 100084, China

**Keywords:** protein structure prediction, protein fitness prediction, mutation effect, end-to-end framework, pre-training

## Abstract

Significant research progress has been made in the field of protein structure and fitness prediction. Particularly, single-sequence-based structure prediction methods like ESMFold and OmegaFold achieve a balance between inference speed and prediction accuracy, showing promise for many downstream prediction tasks. Here, we propose SPIRED, a novel single-sequence-based structure prediction model that exhibits comparable performance to the state-of-the-art methods but with approximately 5-fold acceleration in inference and at least one order of magnitude reduction in training consumption. By integrating SPIRED with downstream neural networks, we compose an end-to-end framework named SPIRED-Fitness for the rapid prediction of both protein structure and fitness from single sequence. SPIRED-Fitness and its derivative SPIRED-Stab achieve state-of-the-art performance in predicting the mutational effects on protein fitness and stability metrics, respectively.

## Introduction

Proteins serve as the primary executors for biological functions typically by folding into unique three-dimensional structures, which highlights the importance of protein structure determination. The structural information of a protein is theoretically prescribed by its amino acid sequence^1^, but could be excavated from the evolution of protein sequences more easily in practice^2–6^. In the past years, deep learning models developed based on this principle have made remarkable progress in the field of protein structure prediction. Representative models, such as AlphaFold2^7^ and RoseTTAFold^8^, utilize evolutionary information extracted from multiple sequence alignments (MSA) to predict protein structures with accuracy close to experimental results. Later on, single-sequence-based predictors, such as ESMFold^9^, OmegaFold^10^, HelixFold-Single^11^, trRosettaX-Single^12^ and RGN2^13^, have been designed to accelerate protein structure prediction by employing the pre-trained protein language model (PLM), which learns evolutionary information hidden in the dependencies between amino acids from hundreds of millions of available protein sequences. As an example, ESMFold and OmegaFold can accomplish the structure prediction for generic proteins within a few seconds to tens of seconds, surpassing the speed of AlphaFold2 by at least an order of magnitude. Despite the amazing progress, current structure prediction methods still have limitations, mainly in the prohibitive computational cost in both inference and model training. Firstly, in situations with a limited number of GPUs (typical for an individual research group), both ESMFold and OmegaFold are still speed-inadequate for high-throughput structure prediction. For instance, it takes around 8 seconds to predict the structure of a 500-residue protein using ESMFold and OmegaFold (without recycling). If applied for tens of thousands or millions of mutant sequences, the aforementioned methods will be time-consuming for the generation of structural features required by downstream protein functional analysis and prediction. Secondly and more importantly, whether it is AlphaFold2 or ESMFold, the training procedure requires significant computational resources that far exceed the affordability of an individual research group. This hinders the extensive explorations on the modification of model architecture, the update of model parameters by re-training, as well as the fine-tuning of structure prediction models for downstream functional analysis.

Considering the decisive impact of protein structures on their functions, one of the ultimate goals of protein structure determination or prediction should be to facilitate the engineering of protein functions through sequence modification^14,15^. In the rational design or engineering, the protein sequence could be changed to alter the protein structure or structural stability, eventually achieving protein functional remodeling. Historically, intensive labors and resources have been invested to screen a large number of mutants for proteins with various desired functions (*e*.*g*., the enzyme activity and protein stability), which are comprehensively named as the protein fitness^16^. Obviously, accurate prediction of the fitness changes caused by single and multiple mutations is of high importance in protein design and functional studies. Numerous previous researches suggest that proper utilization of protein structural information could effectively enhance the accuracy of protein fitness prediction^17–19^. With the rapid development in the field of protein structure prediction, vast structures predicted by AlphaFold2 and ESMFold allow the overcoming of previous challenges caused by the limited number of experimentally resolved protein structures^20–22^. In a recent work, we proposed a method called GeoFitness^23^, which significantly improves the prediction of mutational effects on protein fitness as well as two specific downstream metrics of protein stability, namely the ΔΔ*G* and Δ*T*_*m*_, by utilizing input features extracted from AlphaFold2-predicted structures. Despite the success, such methods are still limited by the lack of communication between the sequence-based structural prediction and the structure-based functional prediction, because results of the former are simply used as input features of the latter. Ideally, the structural prediction algorithm should be integrated as a part of the overall fitness prediction model, which not only enables sufficient communication between the structural prediction module and the fitness prediction module in the whole neural network through end-to-end model training, but also allows comprehensive utilization of existing data of protein sequences, structures and fitness scores to improve the prediction of protein mutational effects. Unfortunately, the heavy computational consumption of models like AlphaFold2 and ESMFold prohibits such an implementation. In this respect, a light-weighted, fast protein structure prediction model with comparable performance to the state-of-the-art methods is still in urgent demand.

To address the limitations mentioned above, in this work, we first developed a novel structure prediction model, SPIRED (<u>s</u>tructural <u>p</u>rediction based on <u>i</u>nter-residue <u>re</u>lative <u>d</u>isplacement), aiming to reduce the computational consumption. Through an innovative design in model architecture and a newly proposed loss function, SPIRED achieves approximately 5-fold acceleration in the inference speed and at least 10-fold reduction in the training cost. Furthermore, SPIRED reaches comparable accuracy to the state-of-the-art methods like OmegaFold on CAMEO^24,25^ and CASP15^26^ benchmarks. Subsequently, by combining SPIRED and downstream graph neural networks, we proposed an end-to-end framework called SPIRED-Fitness, which is capable of rapidly and accurately predicting the protein structure and the fitness changes caused by all single and double mutations from the amino acid sequence simultaneously. Moreover, through end-to-end model fine-tuning, we demonstrated the feasibility of co-training the protein structural prediction module and fitness prediction module within a universal neural network and validated the assistance of this design in the further improvement of the protein functional prediction. Particularly, when re-utilizing the pre-trained SPIRED-Fitness for the prediction of downstream metrics of ΔΔ*G* and Δ*T*_*m*_ caused by mutations, the corresponding program SPIRED-Stab remarkably outperforms all of the other state-of-the-art predictors.

## Methods

### Overview of SPIRED, SPIRED-Fitness and SPIRED-Stab

In this work, we propose a novel algorithm SPIRED to predict the structure from the amino acid sequence of a protein. SPIRED adopts an innovative model design of sequentially arranged Folding Units (Figure 1a,b) and engages a novel Relative Displacement Loss (RD Loss, Figure 1c) to significantly improve the computational efficiency. After sufficient training using the PDB^27^ database, SPIRED achieves comparable prediction accuracy to the other state-of-the-art methods, but with remarkably enhanced inference speed and significantly reduced training cost. The pretrained SPIRED is then utilized as the information extractor in the SPIRED-Fitness model to predict the single and double mutational effects on the protein fitness from the wild-type amino acid sequence (Figure 1d). SPIRED-Fitness is sufficiently optimized using a plethora of multi-labeled deep mutational scanning (DMS)^28^ data. Subsequently, majority of the pre-trained SPIRED-Fitness model is re-utilized in the SPIRED-Stab model to predict the arbitrary mutational effects on the protein stability, namely ΔΔ*G* and Δ*T*_*m*_, given the input of wild-type and mutant sequences (Figure 1e).

**Figure 1.**
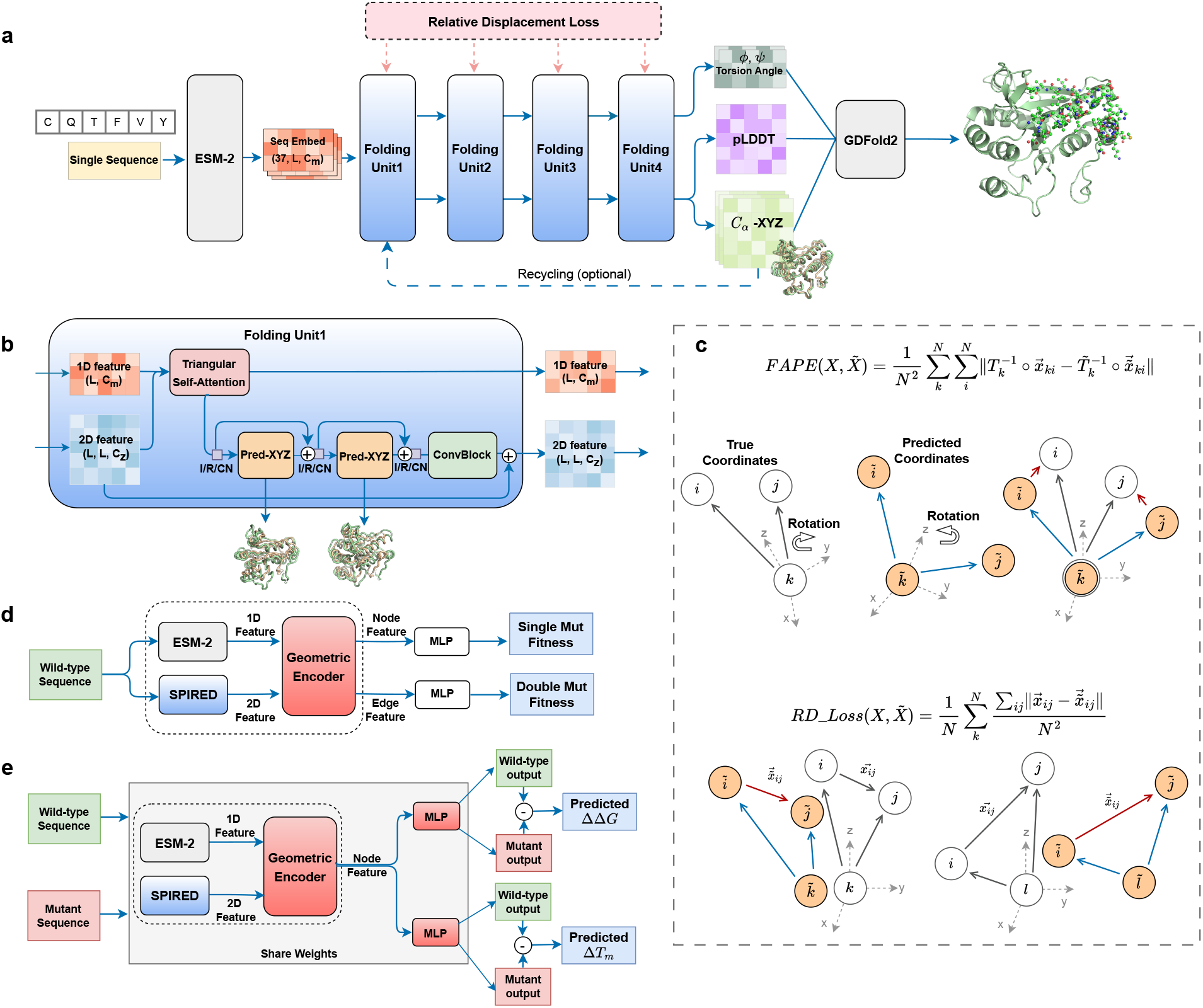
The model architecture of SPIRED, SPIRED-Fitness and SPIRED-Stab. **a)** The model architecture of SPIRED and the protein structure prediction procedure by SPIRED and GDFold2. **b)** Diagram for information flows in the Folding Unit. Each Folding Unit can predict multiple sets of C_*α*_ coordinates. I/R/CN means Instance/Row/Column Normalization. **c)** Comparison between the RD Loss and the FAPE Loss. *T*_*k*_ in the FAPE Loss refers to the rotation matrix for the local coordinate system of residue *k*. In the RD Loss, the relative displacement between a pair of residues *i* and *j* is evaluated over all local reference frames, exemplified by *k* and *l* in the figure. *N* is the number of amino acid residues in the protein. **d)** Prediction of protein fitness upon single and double mutations by SPIRED-Fitness using the wild-type sequence as input. **e)** SPIRED-Stab combines the wild-type sequence and mutant sequence to predict protein stability changes caused by arbitrary mutations, with anti-symmetry guaranteed. For both mutant and wild-type sequences, the model parameters within the gray box have shared weights. The SPIRED-Stab model parameters within the dashed box are initialized based on the final parameters of the SPIRED-Fitness model within the dashed box in subplot (**d**).

### Network architecture of SPIRED

The SPIRED model mainly consists of four Folding Units (Figure 1a, Algorithm 1). When predicting protein structure, SPIRED only requires the amino acid sequence of the target protein, which is encoded into high-dimensional embedding (1D information) by the ESM-2^9^ language model. The sequence embedding is then fed into the Folding Units, in each of which 1D and 2D information mutually updates and multiple sets of coordinates of C_*α*_ atoms are predicted. Unlike mainstream methods like AlphaFold2 and ESMFold that employ the 1D information to predict the atom coordinates in the global frame, in each Folding Unit of SPIRED, the 2D information is used to predict a total of *L* (*i*.*e*. the number of residues) sets of relative coordinates for the C_*α*_ atoms, each taking the local coordinate system of an individual residue as the reference frame. Since both outputs and labels (*i*.*e*. relative C_*α*_ coordinates in individual local frames) are ro-translationally invariant, our design avoids the equivariant operations that usually augment the computational complexity. The multiple C_*α*_ coordinates predicted by the fourth Folding Unit, along with pLDDT and main-chain torsional angles (also composed of 2D matrices), are passed to GDFold2, an in-house folding algorithm based on gradient descent optimization, for main-chain adjustment and side-chain packing, resulting in the full atomic coordinates of the protein.

The network structures of the first three Folding Units are essentially the same. Here, we take Folding Unit1 (Figure 1b, Algorithm 2) as an example to illustrate the network architecture of Folding Units. For Folding Unit1, the 1D feature is the sequence embedding provided by ESM-2 and the 2D feature is initialized as a zero-valued tensor. While for the other Folding Units, the input 1D and 2D features are generated by the preceding Folding Unit. Within a Folding Unit, the 1D and 2D features are first updated by the Triangular Self-Attention module^7,9,29^. The new 1D feature is directly passed on to the next Folding Unit, whereas the updated 2D feature goes through Instance/Row/Column Normalization operations^30^ (I/R/CN in Figure 1b, Algorithm 4), followed by the coordinate prediction module Pred-XYZ (Algorithm 5). The first Pred-XYZ module predicts the absolute C_*α*_ coordinates and generates a new 2D feature that is passed on to the next Pred-XYZ module, while the second Pred-XYZ module predicts additional corrections to the C_*α*_ coordinates. The two Pred-XYZ modules have shared parameters. The pairwise distances between C_*α*_ atoms are then calculated from the predicted coordinates (Algorithm 6). The distance matrix, along with the 2D feature, is then passed to ConvBlock (Algorithm 7), resulting in a new 2D feature that enters the next Folding Unit.

The network architecture of the Folding Unit4 (Algorithm 3) is slightly more complex than the other Folding Units as it engages six Pred-XYZ modules for coordinate prediction and updates, where the first four Pred-XYZ modules and the last two Pred-XYZ modules have two seperate sets of shared weights. The coordinates updated by the last Pred-XYZ module in Folding Unit4 serve as the final C_*α*_ coordinates.

In addition, the 2D feature generated by Folding Unit4 is also utilized to predict the C_*β*_ distance distribution, dihedral and scalar angles quantifying inter-residue orientation (Algorithm 8) as well as main-chain torsion angles (Algorithm 9). Incidentally, each Folding Unit has the capability to predict pLDDT (Algorithm 10), and we consider the pLDDT values output from Folding Unit4 as the representative ones. Finally, since the sequential arrangement of multiple Folding Units yield similar benefits for structure refinement akin to the recurrent expansion by recycling (Figure S1, see **Supplementary Materials** for details), recycling is abandoned by default (*i*.*e*. Cycle = 1) to accelerate inference but could be optionally activated (*e*.*g*., Cycle = 4) in SPIRED.

### Relative Displacement Loss in SPIRED

During the training process of SPIRED, the RD Loss (Figure 1c, Algorithms 11 and 12) is utilized to constrain the C_*α*_ coordinates predicted by each Folding Unit. RD Loss is a novel loss function that is designed to achieve the constraining role of the FAPE Loss^7^ in a computationally less intensive manner. In comparison to the FAPE Loss, it circumvents the laborious coordinate alignment and the costly prediction of rotation matrices, but focuses on evaluating the average prediction accuracy of relative displacement vectors between each pair of C_*α*_ atoms in the multiple reference local coordinate systems.

Before calculating the RD Loss, a local coordinate system is established for each individual residue, where C_*α*_ is set as the origin and the basis vectors are determined from the positions of C_*α*_, C and N atoms, following the AlphaFold2^7^ definition. SPIRED predicts the C_*α*_ coordinates of all residues in each local reference frame. As shown in Figure 1c, in the local coordinate system of residue *k*, the relative displacement between a pair of residues *i* and *j* is evaluated for the predicted structure 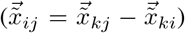 and the ground truth 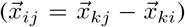, respectively. The RD Loss is then computed as the difference between the predicted and ground truth vectors averaged over all residue pairs and over all reference frames. In contrast, almost all mainstream structure prediction models (*e*.*g*., AlphaFold2, ESMFold and OmegaFold) use the FAPE Loss, which requires predicting quarternions to achieve rotation and laboriously aligning the predicted and true coordinates. Although the offsets between predicted and true coordinates are also evaluated in the FAPE Loss, the inter-residue relative displacement 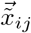 is not specifically considered. Therefore, the RD Loss brings two advantages for training the structure prediction model. Firstly, the RD Loss avoids predicting the rotation matrices, only requiring the prediction of relative positions between residues, thereby alleviating the difficulty of model training of SPIRED. Secondly, the RD Loss places more focuses on the relative displacement between residues, a metric that is more intensively correlated with the inter-residue vibrations rather than the global translation and rotation.

Besides the RD Loss, the inter-residue distance and angle distribution losses are computed based on the C_*β*_ distance distogram as well as the dihedral and scalar angles of all residue pairs following the trRosetta^31^ definition and are utilized as auxiliary losses for the training of SPIRED. In addition, the C_*α*_ distance loss (Algorithm 13), pLDDT loss (Algorithm 14) and C_*α*_ clash loss (Algorithm 15) are also computed as auxiliary losses. Details about the implementation and combination of these losses are described in the **Supplementary Materials**.

### Network architecture of SPIRED-Fitness and SPIRED-Stab

The SPIRED-Fitness model engages ESM-2 and SPIRED as the extractors for 1D and 2D information, respectively (Figure 1d). The downstream Fitness module is mainly composed of the Geometric Encoder that adopts the Graph Attention Network (GAT) architecture (Algorithms 16, 17 and 18) to iteratively update the node and edge features provided by ESM-2 and SPIRED. Specifically, the node feature is initialized by the sequence embedding of ESM-2 (650M), whereas the edge feature includes the multiple sets of C_*α*_ coordinates and the pLDDT values predicted by SPIRED. The updated node and edge features are then fed into MLP (*i*.*e*. multiple layer perceptron) layers for the prediction of fitness changes caused by single and double mutations, respectively. Notably, in the prediction of single mutational effects, the fitness landscape is generated from the 1D MLP output in combination with the ESM-1v^32^ logits, following the procedure of our prior work in GeoFitness v1^23^. As for the prediction of double mutational effects, the fitness scores of all possible mutations for each residue pair are predicted from each individual term of the 2D MLP output directly.

Since the SPIRED-Fitness model could be sufficiently optimized by the plenty of DMS data to learn the general mutational effects, re-utilization of SPIRED-Fitness modules in SPIRED-Stab would effectively overcome the challenge of limited amount of protein stability data. A similar idea has been validated in our prior work on GeoDDG/GeoDTm v1^23^. Specifically, majority of the SPIRED-Fitness model (ESM-2, SPIRED and the Geometric Encoder, as enclosed by a dashed box in Figure 1d) is directly implanted into SPIRED-Stab with the same network architecture and parameters, followed by an MLP layer for the prediction of stability score (Algorithms 19 and 20). Noticeably, SPIRED-Stab retains shared weights for the two channels of inputs, *i*.*e*. the wild-type and mutant sequences, and the difference of their prediction scores is then scaled to predict the absolute values of ΔΔ*G* and Δ*T*_*m*_, a similar design to our prior GeoDDG/GeoDTm v1 models that intrinsically guarantees the anti-symmetry of prediction results.

### Training set for SPIRED structure prediction

First, we collected protein structures available until March 2022 from the PDB^27^ database, but filtered out the structural files with > 5 polypeptide chains and with resolution > 5 Å. Then, we split the remaining structures into multiple single chains and retained chains with length between 40 and 1,200 residues. Next, we clustered these chains using MMseqs2^33^ easy-cluster with the sequence identity threshold of 100% and only kept the representative chains of clusters, which finally resulted in 113,609 chains. We also utilized domain structures from the CATH^34^ database (v4.2, S35) as supplementary training data, which contained 24,183 domains with length ranging from 63 to 600 residues.

### Training process for SPIRED structure prediction

As shown in Table 1, the training process of SPIRED could be mainly divided into four stages.

**Table 1.** SPIRED training protocol.

|  | First stage | Second stage | Third stage | Fourth stage |
| --- | --- | --- | --- | --- |
| **Sequence crop size** | 256 | 256 | 256 | 350 ~ 420 |
| **Batch size** | 64 | 64 | 64 | 64 |
| **Learning rate** | $10^{-3} \sim 5 \times 10^{-4}$ | $5 \times 10^{-4} \sim 10^{-4}$ | $10^{-4} \sim 5 \times 10^{-5}$ | $5 \times 10^{-5} \sim 10^{-5}$ |
| **Sampling strategy** | 30% cluster<br>PDB chains | Easy PDB chains<br>($L < 400, Res < 3\text{\AA}$) | PDB chains +<br>CATH domains | PDB chains +<br>CATH domains |
| **Sample number** | 22K | 63K | 126K | 136K |
| **Update steps** | 10K | 74K | 23K | 30K |

$L$ denotes the length of a protein, while $Res$ refers to the resolution of experimental structures.

First stage, we performed clustering on 101,915 polypeptide chains (before May 2020) with 30% sequence identity using MMseqs2, which resulted in 24,181 clusters. We trained SPIRED for ∼10,000 update steps with the clustered PDB chains, where one chain was iteratively chosen from every cluster in each epoch. During this process, the learning rate was linearly warmed up from 10^−6^ to 10^−3^ in the first 1,000 updates, retained at the peak value of 10^−3^ for the next 6,500 updates, and declined down to 5 *×* 10^−4^ for the final 2,500 updates.

Second stage, we selected an “easy subset” with length < 400 and resolution < 3 Å from the whole training set. We then trained SPIRED with the ∼63,000 “easy subset” chains for ∼74,000 updates. The learning rate was declined from 5 *×* 10^−4^ to 10^−4^ in this stage.

Third stage, we used the whole training set, containing 113,609 PDB chains (before March 2022) and 24,183 CATH domains, to train SPIRED for ∼23,000 updates, with the learning rate annealed from 10^−4^ to 5 *×* 10^−5^. The cropping size was kept at 256 in the first three stages.

Fourth stage, we trained SPIRED for 18,000 updates with the cropping size expanded to 350, and kept the cropping size at 420 for the next 12,000 updates. Learning rate was annealed from 5 *×* 10^−5^ to 10^−5^ during this stage.

The batch size was fixed to 64 and the Adam^35^ optimizer was used throughout the training process of SPIRED.

### Test sets for protein structure prediction

We used two test sets to evaluate the performance of structure prediction methods. The first test set was constructed from CAMEO^24^ targets (August 2022 ∼ August 2023), consisting of 680 protein chains with the length ranging from 50 to 1,126 residues. The second test set was composed of 45 target domains from the CASP15^26^ competition.

We used two kinds of structure classification databases, SCOPe^36^ database (v2.08, S95, September 2021) and CATH^34^ database (v4.2, S35, July 2017), to evaluate the structure prediction power on different types of protein backbone folds or topologies. We selected domains from SCOPe with length ranging from 50 to 800 residues, resulting in 1,231 folds and 34,021 domains in total. Similarly, 1,223 topologies and 24,183 domains were collected from CATH.

### Training and test sets for protein fitness prediction

We utilized DMS data to train and test fitness prediction models, which included data from three different databases.

**cDNA proteolysis dataset**^37^. Tsuboyama *et al*. constructed a library in which mutated proteins were covalently linked to cDNA. These proteins were subsequently subjected to proteolysis, and the cDNA fragments connected to those proteins that were not cleaved could be detected through sequencing, allowing the determination of the quantity of intact proteins at different protease concentrations. Due to the fact that mutated proteins with lower folding stability are more susceptible to proteolytic cleavage in the experiment, protein ΔG values can be inferred using protein cleavage rate data and Bayesian inference. This experimental method facilitates the large-scale analysis of the impact of mutations on protein stability, enabling the examination of folding stability across 900,000 protein domains in a week. From the data provided in the article, we selected 412 proteins with length ranging from 32 to 72 residues to compose a dataset for protein fitness prediction. Of these proteins, 153 proteins have data for both single and double mutations, while the rest only have data for single mutations.

**MaveDB**^38,39^ is a database that contains fitness data of mutated proteins obtained from DMS experiments and massively paralleled reporter assays, including enzymatic activity, binding affinity, *etc*. We selected 51 proteins from MaveDB for the training and testing of our models.

**DeepSequence Dataset**^40^ collects fitness data of mutated proteins from DMS experiments. We selected 22 proteins from this dataset for the subsequent fitness training and testing purposes.

The data from the three aforementioned databases collectively constituted a dataset of 485 proteins, consisting of ∼693,000 single mutations and ∼265,000 double mutations. For each protein, all fitness data were randomly assigned for training, validation and testing with a ratio of 7:1:2.

### Training process for SPIRED-Fitness

The training of SPIRED-Fitness could be mainly divided into two stages.

In the first stage, the SPIRED parameters were frozen and only parameters of the Fitness module were updated for ∼400 epochs. The initial learning rate was set to 10^−3^, and a learning rate scheduler was applied to adjust the learning rate (ReduceLROnPlateau, factor = 0.5, patience = 10). The Fitness module corresponding to the best performance of the fitness prediction on the validation set was used for continued training in the next stage (Fitness module hyperparameters: node_dim = 32, pair_dim = 32, N_head = 8, N_block = 2, see Algorithms 17 and 18). When calculating the loss of this training stage, single mutations and double mutations are combined as a comprehensive mutation set. The Soft Spearman Loss^41^ (see **Supplementary Materials** for details) between the predicted fitness logits and the ground truth values is computed within this mutation set (Equation 1).

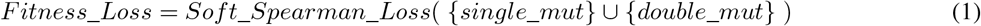

In the second stage, both the SPIRED module and the Fitness module were allowed to update parameters, using training data from two souces: the structural data and fitness data. The structural data were initially taken from the training set of the fourth training stage of SPIRED (see Table 1) with lDDT > 0.5 (∼133,000 protein chains), and were then randomly shuffled and divided into 133 subsets, each of 1,000 samples. The fitness data were the DMS data used in the first stage. For each epoch of the training process, the samples included 1,000 SPIRED training samples and nearly all fitness training samples (from 482 proteins after excluding 3 large proteins with length > 800 residues), in total of 1,482 proteins. After training for 133 epochs, SPIRED-Fitness was fine-tuned on CPU for the three large proteins excluded previously. The learning rate for the SPIRED module was fixed at 10^−5^, while that for the Fitness module was initialized to 10^−4^ and then manually adjusted to 10^−5^. The loss for this stage is represented by the Union Loss defined in Equation 2: the Structural Loss alone was applied for the structural samples, while the joint loss of structure and fitness was applied for the fitness samples. The Structure Loss took the same form as that used in the SPIRED model training (see **Supplementary Materials** for details), but was scaled by a weight of 0.05.

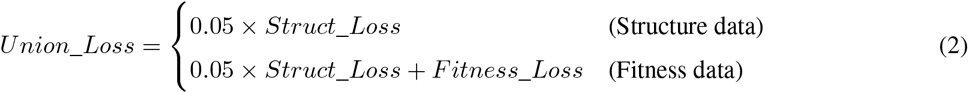

### Training and test sets for protein stability prediction

The datasets utilized for training and testing in SPIRED-Stab are described in details here.

**Dual Task Dataset** is a dataset constructed in this work for training SPIRED-Stab. We collected single, double and triple or higher-order mutation data of both ΔΔ*G* and Δ*T*_*m*_ from two protein stability database, ProThermDB^42^ and ThermoMutDB^43^, and cautiously cleaned each piece of data to generate the dataset for the ΔΔ*G*/Δ*T*_*m*_ dual task training of SPIRED-Stab. The final dataset contains 8,458 pieces of single mutation data, 966 pieces of double mutation data and 619 pieces of triple or higher-order mutation data (*i*.*e*. mutation points ≥ 3), where 5,331 pieces of data only have the ΔΔ*G* label, 2,560 pieces of data only have the Δ*T*_*m*_ label and 2,152 pieces of data have both ΔΔ*G* and Δ*T*_*m*_ labels.

**S669**^44^ is a widely used test set to assess the accuracy of ΔΔ*G* prediction. This dataset consists of 669 single-point mutations derived from 94 proteins selected from ThermoMutDB (v1.3). These proteins have sequence similarity of < 25% with the proteins in the S2648 and VariBench databases that have been extensively used as training data in many previous researches.

**S461**^45^, a subset of S669 dataset with the errors manually corrected, contains 461 single-point mutations. The S461 dataset is used as an auxiliary benchmark test dataset to evaluate ΔΔ*G* prediction.

**S557** is a subset of S571 dataset constructed in our previous work^23^ to specifically address the Δ*T*_*m*_ evaluation problem. We no longer consider pH values, and thus have removed redundancy from the original dataset. This dataset now contains 557 pieces of single mutation data, and is used as an objective benchmark test dataset to evaluate the Δ*T*_*m*_ prediction.

### Training process for SPIRED-Stab

The training of SPIRED-Stab could be divided into three stages. In all training stages, we adopted the Adam^35^ optimizer and the learning rate would decline by half if the validation loss didn’t decrease for five consecutive epochs.

In the first stage, the model parameters of SPIRED-Fitness (except for the MLP module) were used as the starting point of SPIRED-Stab. The training dataset was the cDNA proteolysis dataset described above, and the Soft Spearman Loss was used to evaluate the Spearman correlation coefficient between the predicted ΔΔ*G* and the experimental values. The initial learning rate of this stage was 10^−3^ and all parameters except the final ΔΔ*G*_*coef* and Δ*T*_*m*__*coef* parameters (Algorithm 19) were optimized.

Since the ΔΔ*G* values in the cDNA proteolysis dataset are derived from Bayesian inference, it is necessary to train the model on the ΔΔ*G*/Δ*T*_*m*_ dataset with experimentally measured values. In the second stage, SPIRED-Stab was further trained on our collected and curated ΔΔ*G*/Δ*T*_*m*_ dataset, namely the Dual Task Dataset, with the Soft Spearman Loss employed for the optimization of ranking correlation. In this stage, the MLP layer for predicting ΔΔ*G* was was optimized with the initial learning rate of 5*×*10^−4^ and the learning rate for the MLP layer for predicting Δ*T*_*m*_ was 5 *×* 10^−3^ (Figure 1e).

In the third stage, the numerical difference between the predicted and experimentally determined ΔΔ*G*/Δ*T*_*m*_ values was computed, following the Mean Squared Error (MSE) Loss. During the training in this stage, the majority of the parameters of SPIRED-Stab were frozen, and only the final ΔΔ*G*_*coef* and Δ*T*_*m*__*coef* parameters were updated with the initial learning rate at 10^−2^, aiming for matching the predicted values towards the actual ΔΔ*G*/Δ*T*_*m*_ distribution without perturbing the learned ranking of mutational effects.

### Evaluation metrics

In this study, we utilize TM-score and lDDT to assess the similarity between predicted and true protein structures. Besides, we mainly employ the Spearman correlation coefficient to assess the prediction power of the tested models on fitness, ΔΔ*G* and Δ*T*_*m*_ values. Specifically, we examine the correlation between the predicted logits values and the experimental fitness/ΔΔ*G*/Δ*T*_*m*_ for different mutations.

**TM-score**^46^ (Template Modeling score) is a metric used to assess the topological similarity between protein structures. The protein structure of interest (*i*.*e*. target) is aligned to a reference structure (*i*.*e*. template), and the root-mean-square-deviation (RMSD) of the aligned residue pairs is calculated. According to Equation 3, TM-score ranges from 0 to 1, with the value of 1 indicating a perfect match between structures. TM-score is more sensitive to the global topology than to local structural difference, with the value below 0.17 as an indicator of lack of relationship between the protein structures and a value greater than 0.5 as an indicator of belonging to the same topology.

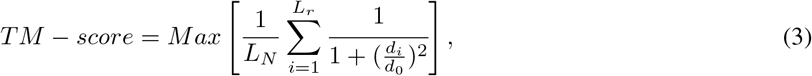

where *d*_*i*_ represents the distance between the *i*^*th*^ aligned residue pairs, *d*_0_ is a normalization scale, *L*_*N*_ denotes the original length of the protein, and *L*_*r*_ represents the number of aligned residues.

**lDDT**^47^ (local Distance Difference Test) is a superposition-free score that indicates the differece in local interresidue distances between the predicted structure and reference structure. First, the distance (*L*_*true*_) between each pair of atoms in the reference structure is calculated, excluding distances beyond the threshold *R*_0_ and atoms within the same residue. The distances (*L*_*pred*_) between corresponding atom pairs are then computed for the predicted structure. Next, the absolute difference in distances between the two structures for each atom pair is calculated (Diff = |*L*_*true*_ − *L*_*pred*_|). The counts of atom pairs with Diff values below four thresholds (0.5, 1, 2, and 4 Å) are calculated, and the average of these counts divided by the total number of atom pairs produces the lDDT score. In this study, we only calculate the lDDT score for C_*α*_ atoms (lDDT-C_*α*_) with *R*_0_ = 15 Å.

**Pearson correlation coefficient** (*r*) is a measure used to quantify the strength of the linear relationship between two variables, *X* and *Y*. As shown in the Equation 4, it is computed by calculating the ratio of the covariance between the two variables to the product of their standard deviations. The coefficient ranges from -1 to 1, where -1 indicates a perfect negative correlation, 0 indicates no correlation, and 1 indicates a perfect positive correlation.

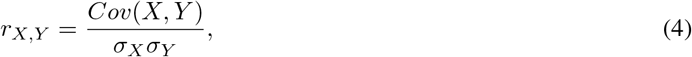

where *Cov* denotes the covariance and *σ* stands for the standard deviation.

**Spearman correlation coefficient** (*ρ*) is commonly used to describe the strength of a monotonic relationship between two variables. As shown in the Equation 5, Spearman correlation coefficient is calculated by utilizing the ranked values of both variables (*X, Y*). This characteristic makes the Spearman correlation coefficient more robust to outliers in the data.

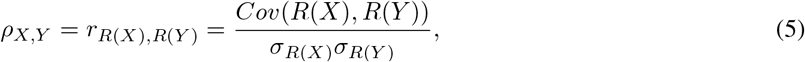

where *R* denotes the ranking operation for the variables.

**Kendall correlation coefficient** (*τ*) is another non-parametric metric to measure the correlation between ranks of the variables and can be interpreted as the probabilities of observing the agreeable (concordant) and non-agreeable (discordant) pairs (Equation 6). The Kendall correlation is more robust than the Spearman correlation while usually being smaller in magnitude.

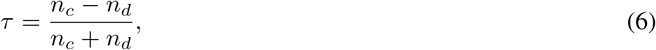

where *n*_*c*_ denotes the number of concordant pairs while *n*_*d*_ denotes the number of discordant pairs.

**Top K precision** is a metric measuring the fraction of the truely top K mutations among the predicted top K mutations (Equation 7). This metric serves as a reference for the success rate in the real world protein engineering process.

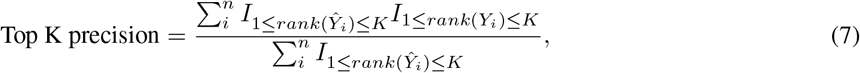

where *rank*(*Ŷ*_*i*_) and *rank*(*Y*_*i*_) denote the rank (in descending order) of the predicted value and that of the label, respectively, and *I* is the indicator function.

## Results

### SPIRED performs well for CAMEO and CASP15 targets without recycling

To validate the performance of our protein structure prediction method, we evaluate SPIRED against two state-of-the-art models, ESMfold and OmegaFold, on CAMEO and CASP15 targets at two options: Cycle = 1 (*i*.*e*. without recycling) and Cycle = 4 (*i*.*e*. four recycling cycles). GDFold2 is used to perform side-chain packing and main-chain adjustment for the structures predicted by SPIRED (SPIRED+GDFold2 in Figure 2). The CAMEO set has 680 single-chain proteins (released from August 2022 to August 2023). The CASP15 set contains 45 publicly released protein domains. Proteins in these test sets are all released after the date cutoff (March 2022) of the training set of SPIRED.

**Figure 2.**
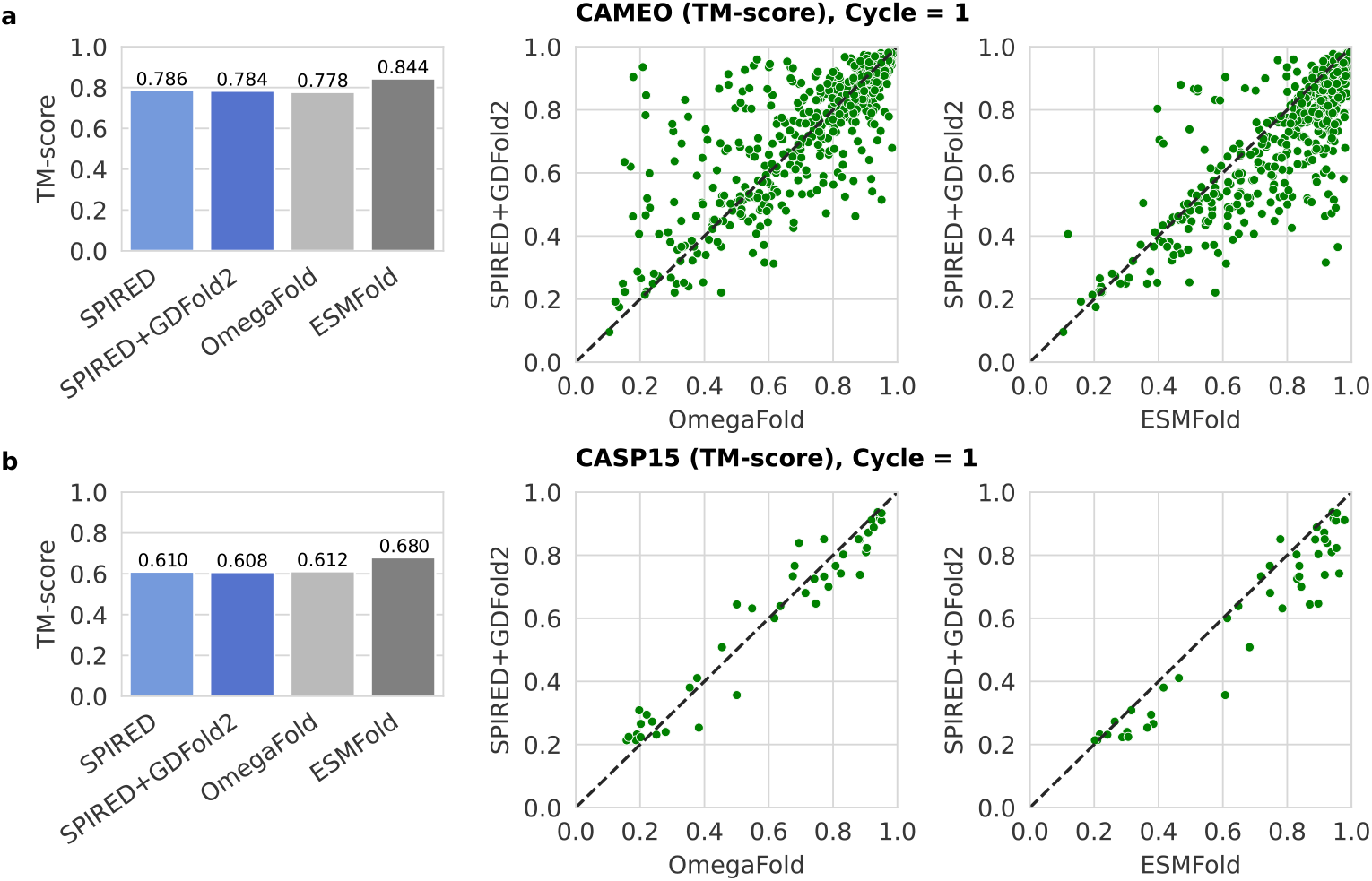
Comparison of model performance on CAMEO and CASP15 targets. **a)** The barplot displays the TM-scores of predicted structures by SPIRED, SPIRED+GDFold2, OmegaFold, and ESMFold on the CAMEO set (680 proteins) with Cycle = 1. In the scatter plot, each point represents a protein, with the vertical axis representing the SPIRED+GDFold2 prediction and the horizontal axis representing the results of OmegaFold and ESMFold, respectively. **b)** Similar evaluation of models on the CASP15 set (45 protein domains) with Cycle = 1. All models make predictions from the domain sequences released by the CASP15 official website. The TM-score is calculated by comparing the predicted structures with the released CASP15 domain structures.

Without recycling (Cycle = 1), SPIRED performs well on the CAMEO set (average TM-score = 0.786), slightly surpassing OmegaFold (average TM-score = 0.784), as shown in Figure 2a. When four recycling cycles are employed (Cycle = 4), SPIRED exhibits a slightly lower prediction accuracy compared to OmegaFold: TM-score of 0.787 vs. 0.805 (Figure S2a). As for the CASP15 targets, SPIRED exhibits a similar prediction accuracy to OmegaFold at both options (Figure 2b for Cycle = 1 and Figure S2b for Cycle = 4). Clearly, ESMFold shows better performances than SPIRED and OmegaFold on both CAMEO and CASP15 sets. This is, however, not unexpected considering that the model parameters of ESMFold outnumber those of SPIRED and OmegaFold by approximately five times, and that ESMFold engages a large amount of AlphaFold2-predicted proteins for model training (see Table 2).

**Table 2.**
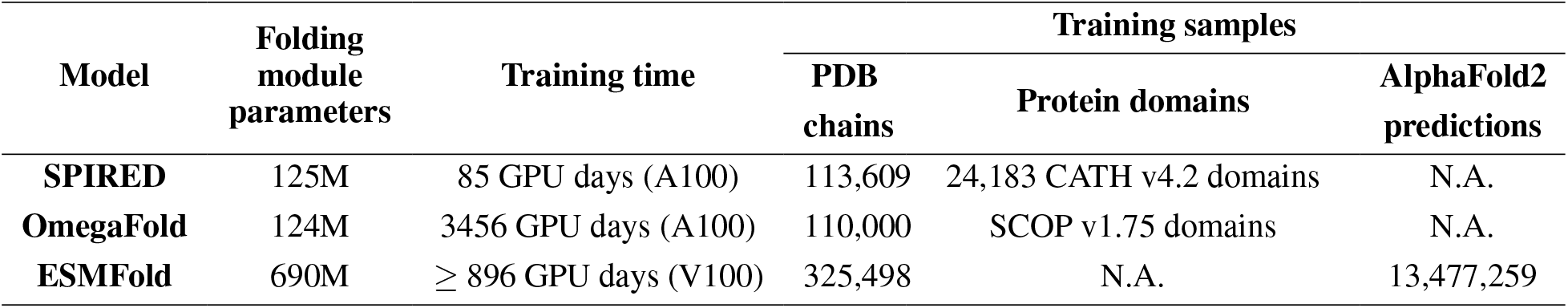
Model parameters, training time and training samples.

Nevertheless, SPIRED exhibits considerable prediction accuracy in the absence of recycling, with a generally comparable performance to the nowadays state-of-the-art single-sequence-based protein structure prediction models, which supports its candidate role for end-to-end training with downstream protein function prediction networks. Other Models like OmegaFold rely on recycling to improve prediction accuracy, and the truncation of gradients during recycling obviously hinders the joint training of such models with downstream networks in the recycling state. ESMFold, despite its high prediction accuracy without recycling, has vast model parameters and consumes large amounts of training memory, thus exceeding the affordability of ordinary research groups for the similiar joint training.

### Evaluation of structure prediction at the level of protein folds

In previous studies, the performance evaluation of structure prediction models often focused on dozens or hundreds of proteins from double-blinded competitions, which covered a limited range of topological structure types and therefore could not comprehensively evaluate the model performance on the protein folds of common interests. Therefore, in this study, we systematically analyze the model performance at the level of all known protein folds (or topologies). The SCOPe database categorizes experimentally determined protein structures into different folds, each containing one or more structural domains. We use SPIRED, ESMFold and OmegaFold to predict the structures for 34,021 SCOPe domains belonging to a total of 1,231 folds, and use the average TM-score of all domains within each fold as the indicator of the model performance for this specific type of topology.

In general, SPIRED exhibits significantly better performance than OmegaFold and ESMFold on numerous folds (see Figure 3a,b). Particularly, SPIRED shows an advantage of > 0.2 in TM-score over ESMFold in 91 folds, whereas ESMFold outperforms SPIRED by > 0.2 in TM-score only in 14 folds. The kernel density estimate (KDE) plot of TM-scores across different folds (Figure 3c) clearly suggests that SPIRED has the lowest density in the low prediction accuracy region (*i*.*e*. 0.2 < TM-score < 0.5) and the highest density in the high accuracy region (*i*.*e*. TM-score = ∼0.9) among the tested models. Furthermore, SPIRED has a higher average TM-score over all folds than both OmegaFold and ESMFold (Figure 3d). Taking the GFP protein folding type (SCOPe Fold ID: d.22) as an example, SPIRED has an average TM-score of 0.959, while OmegaFold and ESMFold can only achieve average TM-scores of 0.577 and 0.485, respectively. More case studies of successful SPIRED predictions could be found in Figures S3 and S4.

**Figure 3.**
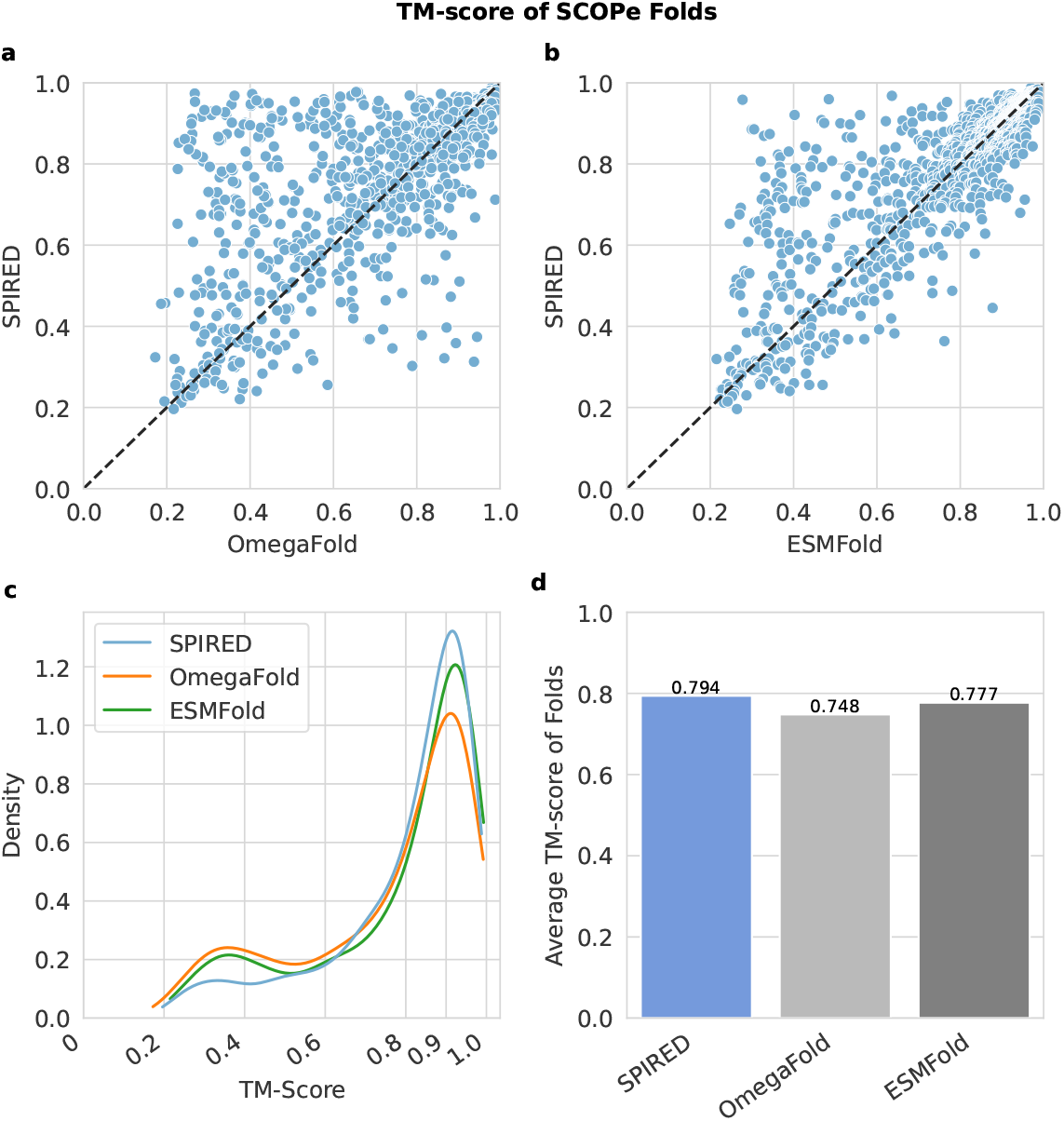
Comparison of the prediction accuracy on the SCOPe v2.08 S95 structural classification database. **a-b)** In the scatter plot, each point represents the average TM-score of all domains of one fold in SCOPe. The veritcal axis is the prediction performance (TM-score) of SPIRED on different folds, while the horizontal axis represents results for OmegaFold and ESMFold, respectively. **c)** The KDE plot is used to visualize the distribution of model performance (in TM-score) over different folds for the tested models. **d)** The bar plot compares the TM-score averaged over all folds for different models.

The same phenomenon is observed when evaluated at the level of CATH topologies (Figure S5, see **Supplementary Materials** for details). The advantage of SPIRED over the other state-of-the-art models at the protein fold/topology level supports its comprehensive competitiveness in facilitating the downstream functional analysis as well as the protein design and engineering.

### Remarkable advantage of SPIRED in training time and inference speed

SPIRED significantly prevails OmegaFold and ESMFold in terms of training consumption. As shown in Table 2, the training of SPIRED only costs 85 GPU days, in sharp contrast to the 3,456 GPU days of OmegaFold (derived from the Supplementary 2.8 and Table S3 of the OmegaFold paper) and the ∼896 GPU days of ESMFold (obtained by communication with the first author). Therefore, in comparison to the other state-of-the-art methods, SPIRED effectively reduces the training cost by at least one order of magnitude, mainly through the innovative design on the network architecture and loss function. On the other hand, the number of parameters in the structure prediction module of SPIRED (125M) is in a similar level to that of OmegaFold (124M), both of which is much less than that of ESMFold (690M). Regarding the training samples, both ESMFold and OmegaFold primarily focus on single-chain proteins from the PDB database, supplemented with protein domains from available structural classification databases. In contrast, besides the PDB data, ESMFold also incorporates a large amount of high-quality structures predicted by AlphaFold2 in its training set, which would further exacerbate the training cost.

Based on evaluation upon the model inference speed, we find that SPIRED is approximately 5 times faster than ESMFold and OmegaFold. The time consumption for inferring proteins with length ranging from 100 to 1,000 residues on the NVIDIA A100 GPU (80GB) is shown in the Figure 4. Without recycling, SPIRED takes < 1 second when inferring proteins smaller than 400 residues. For example, when inferring a protein of 300 residues, SPIRED takes approximately 0.5 seconds, while ESMFold and OmegaFold take 2.7 seconds and 2.1 seconds, respectively. For proteins of 600 residues, SPIRED takes ∼2.1 seconds, while ESMFold and OmegaFold take 13.5 seconds and 12.1 seconds, respectively. Even when recycling is activated at Cycle = 4, SPIRED still maintains a similar speed advantage in model inference.

**Figure 4.**
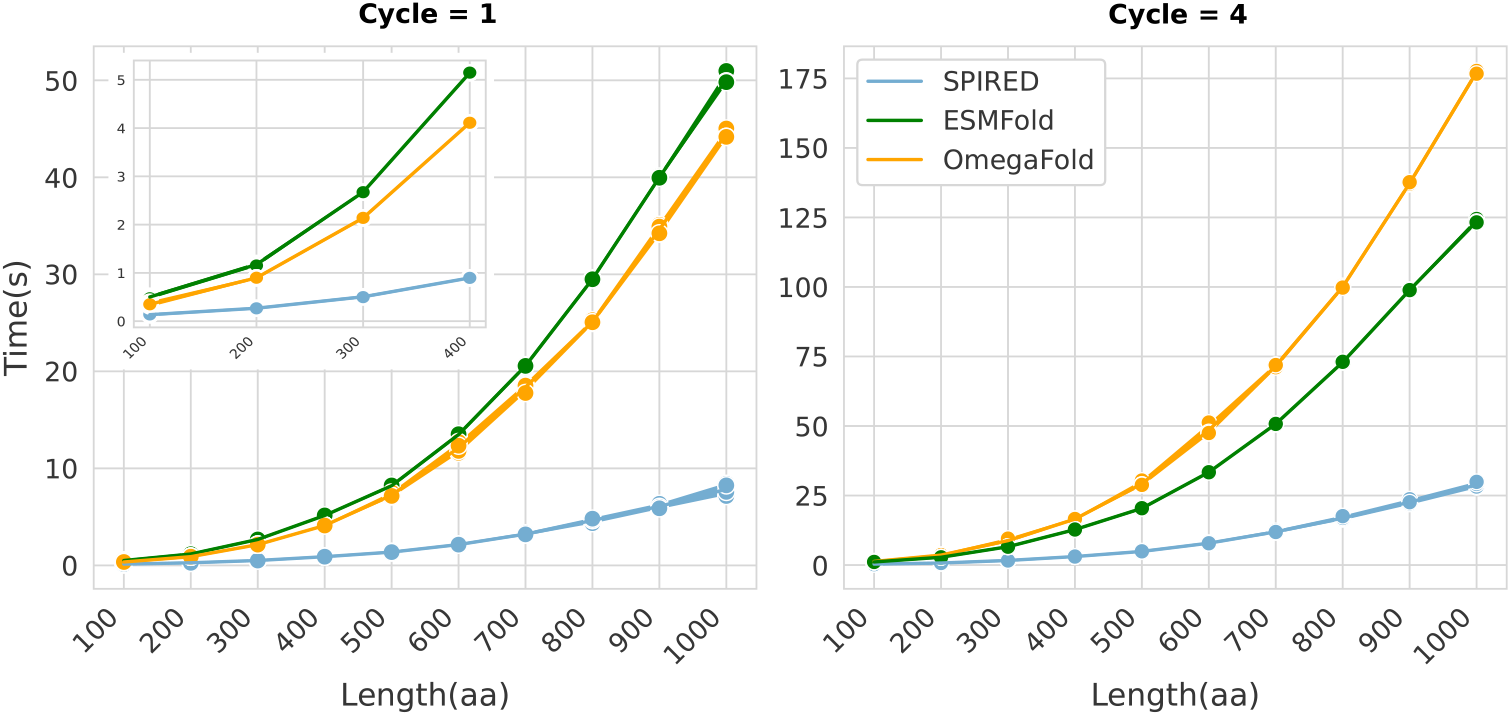
Comparison of inference time. The horizontal axis represents the length of the proteins, and the vertical axis represents the time taken for inference. In order to generate the plots, protein lengths are binnized into 10 intervals ranging from 100 to 1000 and 4 different proteins are selected in each interval to evaluate the time cost for structure prediction.

In summary, the significant advantage of SPIRED in both training cost and inference speed implies its potential in the high-throughput protein structure prediction as well as in the joint training with downstream protein funtional prediction models.

### Protein fitness prediction by SPIRED-Fitness

Based on the good performance, low training cost and high inference speed of SPIRED, we integrate this model into the protein fitness prediction network and compose an end-to-end framework named SPIRED-Fitness. As mentioned in **Methods**, we train the SPIRED-Fitness model primarily using a plethora of multi-labeled DMS data, assisted by PDB data to prevent biasing by the limited number of proteins in the DMS dataset. Notably, in the second stage of model training, parameters in the SPIRED module are released for optimization, aiming for further performance improvement by end-to-end training. The final SPIRED-Fitness model takes a single sequence as input and outputs the predicted structure as well as the predicted protein fitness changes caused by all possible single and double mutations in the time scale of seconds for generic proteins.

Here, we compare the fitness prediction performance of SPIRED-Fitness with two state-of-the-art protein fitness prediction models, including ECNet^48^ and our GeoFitness model (updated version, named GeoFitness v2), using all test data from 485 proteins. The GeoFitness v2 adopts the similar model architecture and training procedure with GeoFitness v1^23^, but is retrained with both single and double mutational data as described in the **Supplementary Materials**. As shown in Table 3, when tested on all single and double mutational data, SPIRED-Fitness slightly out-performs both ECNet and GeoFitness v2, in terms of the average Spearman correlation coefficient between predicted and experimental values: 0.85 vs. 0.84 and 0.83. Based on detailed protein-wise comparison (Figure S6), SPIRED-Fitness (Stage 2), the end-to-end training version, surpasses both GeoFitness v2 and SPIRED-Fitness (Stage 1), the version with frozen SPIRED parameters. Specifically, the end-to-end-training in Stage 2 introduces a gain of 2% in the fitness prediction, with only an acceptable minor loss in the structure prediction accuracy (Table S1, see **Supplementary Materials** for details). This observation further supports the postive impact of end-to-end training frameworks in the field of protein functional prediction. It is noteworthy that the apparently high performance of ECNet should be less concerned, since ECNet can only process single-labeled data and thus has to be retrained for each specific protein target with known data before inference, whereas both SPIRED-Fitness and GeoFitness v2 are universal models that only need to be trained once using the whole multi-labeled DMS data and are able to provide predictions for unseen proteins. Consequently, during the overall evaluation process, both SPIRED-Fitness and GeoFitness v2 cost significantly less time than ECNet (Table 3), by the avoidance of model retraining. Incidentally, SPIRED-Fitness is faster by 1900 times than GeoFitness v2, because of the highly efficient protein structure prediction by SPIRED in comparison to the AlphaFold2 employed in the GeoFitness pipeline for feature generation.

**Table 3.** Fitness prediction performance and time consumed for evaluation.

| Model | $\rho_{\text{single and double}}$ | Feature Generation<br>Time (sec) | Model Evaluation<br>Time (sec) | Total Time (day) |
| --- | --- | --- | --- | --- |
| ECNet | 0.84 | 325246 | 16496728 | 194.7 |
| GeoFitness v2 | 0.83 | 330223 | **32** | 3.8 |
| SPIRED-Fitness (Stage 1) | 0.83 | **115** | 62 | **0.002** |
| SPIRED-Fitness (Stage 2) | **0.85** | **115** | 62 | **0.002** |

The best performance in each metric is highlighted in bold. Here, $\rho_{\text{single and double}}$ denotes the Spearman correlation coefficient calculated for all single and double mutations of test data from 485 proteins described in **Methods**. The model evaluation time is calculated based on the 412 proteins from the cDNA proteolysis dataset.

Since most methods in the field of fitness prediction only allow inference on single mutational effects, for a broader performance comparison, we trained SPIRED-Fitness using the single mutational data in the training set and evaluated its performance against unsupervised models including RF_joint_^49,50^, MSA Transformer^6^, ESM-2^9^, ESM-1b^51^ and ESM-1v^32^ as well as supervised models including ECNet^48^ and SESNet^17^ also on the single mutational data in the test set. As shown in Figure S7, SPIRED-Fitness outperforms all unsupervised models on the vast majority of proteins and its overall performance is also better than the supervised models (average Spearman correlation coefficient of 0.87 in SPIRED-Fitness vs. 0.82 in ECNet and 0.83 in SESNet). Furthermore, we find that as the training sample size decreases, ECNet quickly loses its prediction power, in sharp contrast to the mild decline in the performance of SPIRED-Fitness (Figure 5a). Particularly, with only 10% of the training samples, the Spearman correlation coefficients of SPIRED-Fitness and GeoFitness v2 still stay above 0.7, while that of ECNet goes below 0.4, indicating that in the few-shot fitness prediction scenarios, SPIRED-Fitness and GeoFitness v2 are much more robust than ECNet, due to the effective utilization of multi-labeled data in model training. To further evaluate the generalizability of SPIRED-Fitness and GeoFitness v2, we conduct 10-fold cross validations, in each experiment of which 80% proteins are chosen for training and validation while 20% unseen protein are left for testing (Figure 5b). In such a test mimicking the zero-shot prediction scenarios, both SPIRED-Fitness and GeoFitness v2 achieve good performance (median of the average Spearman correlation efficient > 0.7).

**Figure 5.**
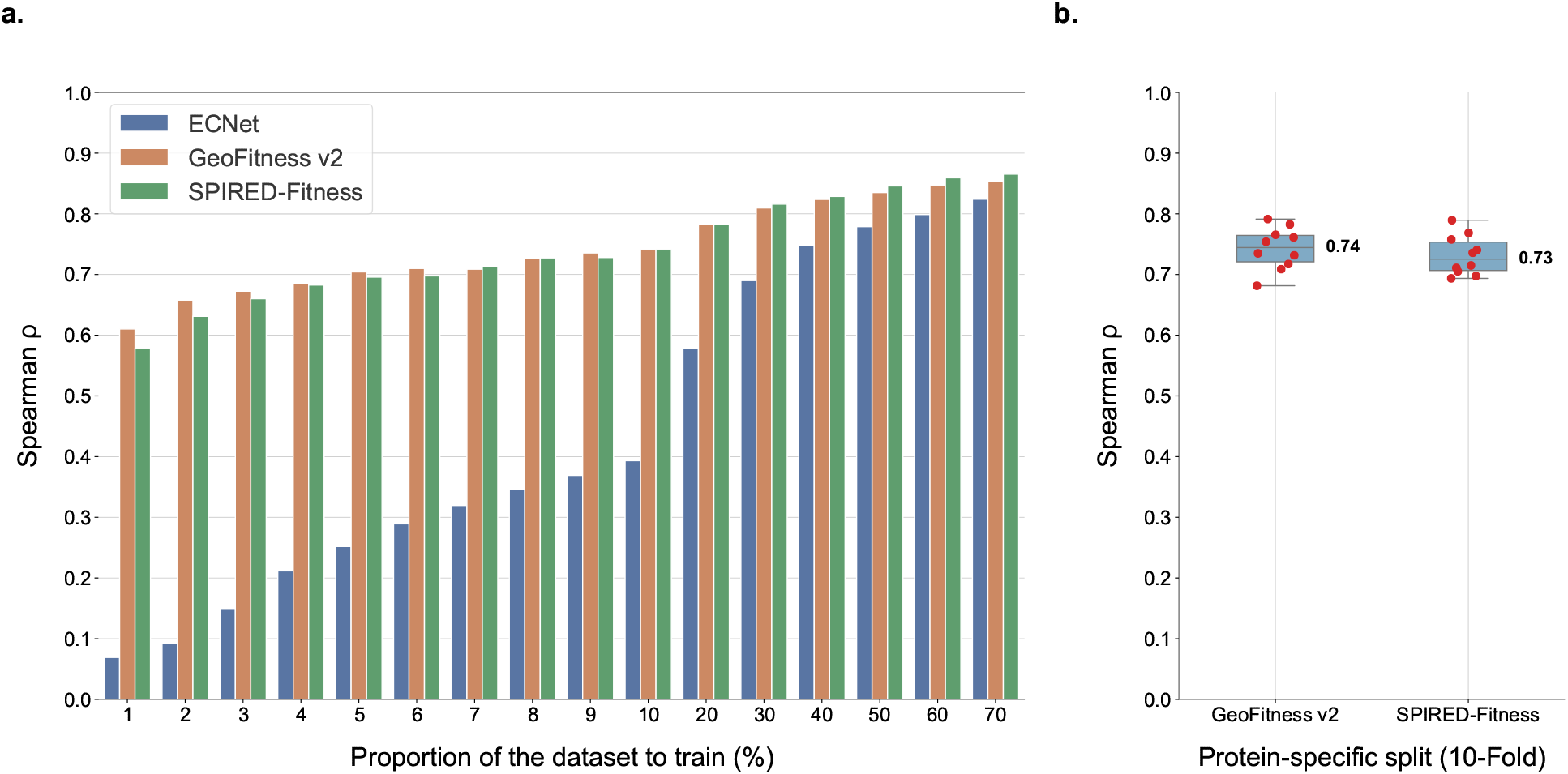
Detailed analysis of SPIRED-Fitness. **a)** Comparison of ECNet, GeoFitness v2 and SPIRED-Fitness when trained with various proportions of data on single mutations of 485 proteins. The full training set contains 70% of the single mutational data in the whole dataset, which corresponds to the maximum value on the horizontal axis in the bar chart. Each bar represents the Spearman correlation coefficient averaged over the 485 proteins. **b)** 10-fold cross validation of GeoFitness v2 and SPTRED-Fitness with protein-specific data splitting, which means that 80% proteins are used for training/validation and 20% unseen proteins are used for testing. Each red dot represents the result of one individual experiment in the 10-fold cross validation. The center line of each box plot shows the median of the validation results with the value marked aside. The box limits correspond to the upper and lower quartiles, whereas the whiskers extend to points that lie within 1.5 inter-quartile range of the lower and upper quartiles.

Hence, the newly developed end-to-end framework SPIRED-Fitness greatly enhances the power and convenience of protein fitness prediction, considering its high accuracy, robustness, broad generalizability, significant speed advantage and bypass of cumbersome feature engineering.

### Prediction of the mutational effects on protein stability by SPIRED-Stab

Considering that SPIRED-Fitness effectively learns the general mutational effects, we re-utilize of its major components in the SPIRED-Stab model to overcome the challenge of limited amount of labeled data for protein stability. As a proof of principle for this idea, we first test the zero-shot prediction behavior of SPIRED-Fitness on the S669/S461 and S557 datasets (see **Methods**) for ΔΔ*G* and Δ*T*_*m*_ predictions, respectively. Subsequently, we train the SPIRED-Stab model using the Dual Task Dataset and then evaluate its prediction behaviors for ΔΔ*G* and Δ*T*_*m*_ prediction tasks on the S669, S461 and S557 datasets. Notably, during conventional evaluation upon the ΔΔ*G*/Δ*T*_*m*_ predictors, the Spearman/Pearson correlation coefficients are estimated over all variants across proteins in the test set. Such a cross-protein evaluation approach will bring artifacts to the evaluation results, since relative ranking/magnitude of ΔΔ*G*/Δ*T*_*m*_ between variants from different proteins is less informative to the practical protein engineering that typically focuses on mutations within one individual protein. Consequently, in this work, we evaluate metrics like the Spearman correlation coefficient within each individual protein and then average the values over all proteins as indicators for the overall performance. Exemplar results of the conventional evaluation approach are shown for the S461 dataset in the **Supplementary Materials**.

As shown in Table 4, Table S2 and Table 5, the zero-shot prediction by SPIRED-Fitness has already surpassed most of the mainstream ΔΔ*G*/Δ*T*_*m*_ predictors, achieving comparable or even better performance to our old version of GeoDDG and GeoDTm that have been sufficiently trained using experimental ΔΔ*G*/Δ*T*_*m*_ labels as well as a recently proposed method Mutate Everything^52^, in the ΔΔ*G* and Δ*T*_*m*_ prediction tasks. Hence, SPIRED-Fitness has indeed learned the mutational effects from the DMS data fairly well and is capable of predicting the protein stability change with considerable power even in the absence of protein stability data. Furthermore, after sufficient training using the experimental ΔΔ*G*/Δ*T*_*m*_ data, SPIRED-Stab shows an additional performance enhancement as expected. Specifically, SPIRED-Stab and GeoStab v2 (updated from GeoFitness v2 as descripted in the **Supplementary Materials**) lead the other methods significantly in nearly all metrics. Noticeably, in addition to the slight advantage in performance, SPIRED-Stab is faster than GeoStab v2 by orders of magnitude in practice, due to the avoidance of MSA feature generation in AlphaFold2.

**Table 4.** Comparison of SPIRED-Stab with other ΔΔ*G* predictors on the S669 dataset.

| Method | $\rho$ | $\tau$ | $r$ | Top-5 | Top-10 |
| --- | --- | --- | --- | --- | --- |
| *Structure-based* |  |  |  |  |  |
| SDM<sup>53</sup> | 0.31 | 0.22 | 0.33 | 0.33 | 0.57 |
| ThermoNet<sup>54</sup> | 0.31 | 0.22 | 0.38 | 0.31 | 0.54 |
| mCSM<sup>55</sup> | 0.33 | 0.23 | 0.35 | 0.39 | 0.53 |
| Dynamut<sup>56,57</sup> | 0.34 | 0.25 | 0.34 | 0.32 | 0.52 |
| I-Mutant3.0<sup>58</sup> | 0.34 | 0.26 | 0.39 | 0.39 | 0.54 |
| MAESTRO<sup>59</sup> | 0.35 | 0.26 | 0.40 | 0.37 | 0.59 |
| DUET<sup>60</sup> | 0.38 | 0.27 | 0.41 | 0.40 | 0.56 |
| ACDC-NN<sup>61</sup> | 0.40 | 0.30 | 0.42 | 0.44 | 0.57 |
| FoldX<sup>62</sup> | 0.41 | 0.31 | 0.36 | 0.36 | 0.59 |
| PopMusic<sup>63</sup> | 0.42 | 0.30 | 0.45 | 0.37 | 0.63 |
| DDGun3D<sup>64</sup> | 0.43 | 0.32 | 0.48 | 0.41 | 0.62 |
| PremPS<sup>18</sup> | 0.45 | 0.33 | 0.48 | 0.39 | 0.58 |
| DDMut<sup>19</sup> | 0.46 | 0.35 | 0.52 | 0.44 | 0.59 |
| INPS3D<sup>65</sup> | 0.46 | 0.35 | 0.48 | 0.44 | 0.59 |
| GeoDDG-3D (old)<sup>23</sup> | 0.50 | 0.38 | 0.51 | 0.53 | **0.69** |
| *Sequence-based* |  |  |  |  |  |
| MuPro<sup>66</sup> | 0.18 | 0.13 | 0.21 | 0.31 | 0.53 |
| I-Mutant3.0-Seq<sup>58</sup> | 0.22 | 0.16 | 0.28 | 0.33 | 0.51 |
| SAAFEC-Seq<sup>67</sup> | 0.34 | 0.26 | 0.39 | 0.39 | 0.54 |
| ACDC-NN-Seq<sup>61</sup> | 0.39 | 0.29 | 0.44 | 0.43 | 0.58 |
| INPS-Seq<sup>65</sup> | 0.40 | 0.28 | 0.44 | 0.47 | 0.56 |
| DDGun<sup>64</sup> | 0.45 | 0.34 | 0.47 | 0.44 | 0.56 |
| GeoDDG-Seq (old)<sup>23</sup> | 0.51 | 0.39 | 0.52 | 0.49 | 0.63 |
| Mutate Everything (ESM-2)<sup>52</sup> | 0.50 | 0.38 | 0.52 | 0.41 | 0.62 |
| SPIRED-Fitness (zero-shot) | 0.54 | 0.41 | 0.52 | 0.51 | 0.64 |
| GeoStab v2 | 0.55 | 0.43 | 0.54 | 0.52 | **0.69** |
| SPIRED-Stab | **0.58** | **0.46** | **0.55** | **0.59** | 0.67 |

The winner in each metric is highlighted in bold. All metrics are first calculated within each protein and then averaged over all proteins. Here, $\rho$ , $\tau$ and $r$ denote Spearman, Kendall and Pearson correlation coefficients, respectively, while Top-5 and Top-10 refer to the Top 5 precision and Top 10 precision, respectively, as defined in **Methods**.

**Table 5.** Comparison of SPIRED-Stab with other Δ*T*_*m*_ predictors on the S557 dataset.

| Method | $\rho$ | $\tau$ | $r$ | Top-5 | Top-10 |
| --- | --- | --- | --- | --- | --- |
| *Structure-based* |  |  |  |  |  |
| AUTO-MUTE<sup>68,69</sup> | 0.23 | 0.19 | 0.27 | 0.33 | 0.61 |
| HoTMuSiC<sup>70</sup> | 0.32 | 0.24 | 0.30 | 0.36 | 0.64 |
| GeoDTm-3D (old)<sup>23</sup> | 0.40 | 0.29 | 0.39 | 0.40 | **0.69** |
| *Sequence-based* |  |  |  |  |  |
| GeoDTm-Seq (old)<sup>23</sup> | 0.45 | 0.33 | 0.41 | 0.46 | 0.66 |
| Mutate Everything (ESM-2)<sup>52</sup> | 0.40 | 0.31 | 0.38 | 0.39 | 0.60 |
| SPIRED-Fitness (zero-shot) | 0.46 | 0.35 | **0.47** | 0.46 | 0.62 |
| GeoStab v2 | 0.49 | 0.38 | 0.44 | 0.44 | **0.69** |
| SPIRED-Stab | **0.50** | **0.39** | 0.43 | **0.47** | **0.69** |

The winner in each metric is highlighted in bold. All metrics are first calculated within each protein and then averaged over all proteins. Here, $\rho$ , $\tau$ and $r$ denote Spearman, Kendall and Pearson correlation coefficients, respectively, while Top-5 and Top-10 refer to the Top 5 precision and Top 10 precision, respectively, as defined in **Methods**.

In conclusion, SPIRED-Stab developed from the end-to-end framework SPIRED-Fitness remarkably improves the accuracy and speed for the prediction of protein stability metrics, ΔΔ*G* and Δ*T*_*m*_, caused by arbitrary mutations.

## Discussion

Currently, mainstream single-sequence-based protein structure prediction models exemplified by ESMFold and OmegaFold tend to adopt structure folding modules similar to AlphaFold2 in order to achieve high prediction performance. Albeit successful, this approach also brings new issues. Firstly, models engaging AlphaFold2-type structural folding module require considerable time and vast computational resources to accomplish model training, which nearly precludes the chance of ordinary research groups to update model parameters by re-training, to freely modify the model architecture, and/or to fine-tune the model for downstream tasks. Secondly, the running time and memory costs of these models are still unsatisfactory, which not only hinders high-throughput inference required by downstream functional analysis, but also prohibits the integration of them with downstream models for end-to-end training. In this work, we introduce an initial endeavor to address this problem. By designing an innovative network architecture (*i*.*e*. the Folding Units) for structural modeling and proposing a novel loss function (*i*.*e*. the Relative Displacement Loss) for structural constraining, we successfully reduce the model training consumption in single-sequence-based protein struture prediction algorithms by at least one order of magnitude and improve the model inference speed by 4-5 times. Moreover, our SPIRED model shows a comparable performance to OmegaFold on CAMEO and CASP15 targets, and outperforms both ESMFold and OmegaFold when evaluated on all known protein folds or topologies, targets that are more relevant for the downstream functional analysis as well as the practical protein design and engineering. Our endeavor paves the way for the joint training of sequence-based structure prediction model and structure-based functional network in an end-to-end manner.

The deep learning data in biology exhibits a highly diverse nature. Protein fitness data, encompassing different types of labels such as protein stability, enzyme activity and binding affinity, differ from the single-type labels in language learning and image recognition. The integration of multiple small pieces of data with highly variable labels is crucial for improving the protein fitness prediction. In our prior work on GeoFitness^23^, we used the Soft Spearman Loss to leverage the multi-labeled data and successfully constructed a universal fitness prediction model with state-of-the-art performance. In this study, we construct a single-sequence version of the fitness prediction model, SPIRED-Fitness, by integrating the structure prediction module SPIRED and the fitness prediction module into an end-to-end framework. By this means, the model inference is accelerated by 1900 folds (in comparison with GeoFitness v2), through bypassing the time-consuming sequence alignment and structural modeling processes of AlphaFold2. More importantly, we demonstrate that the end-to-end training from sequence to structure to function can improve the prediction of single and double mutational effects by around 2% to 3%. Such an end-to-end scheme may be extended to other fields like the protein design, where the joint training of structure-based sequence generation modules and sequence-based structure prediction modules is expected to further improve the foldability of designed sequences.

The mutational effects on protein stability constitute an important problem within the scope of protein fitness. We achieve the state-of-the-art prediction for ΔΔ*G* and Δ*T*_*m*_ in SPIRED-Stab using a Russian-doll-style pre-training approach (Figure 6). Specifically, SPIRED-Stab is trained by a limited amount of protein stability data, but the SPIRED-Fitness module within this model has been pre-trained by a plethora of DMS data. Similarly, SPIRED-Fintess is trained by multi-labeled DMS data, but the SPIRED module within this model has been pre-trained by vast uniformly labeled PDB data. At the next level, SPIRED is trained by tens of thousands of pieces of protein structure data, but the ESM-2 module within this model has been pre-trained by billions of pieces of sequence data. Through such an hierarchical training scheme, our final SPIRED-Stab model benefits greatly from the comprehensive utilization of data from various sources, *e*.*g*., the sequence database, the structure database, the protein fitness data and the protein stability data. Such a pre-training strategy may be extended to the prediction of other biological properties, considering the “fragmented” and “multi-labeled” characteristics of most data in biological and medical sciences.

**Figure 6.**
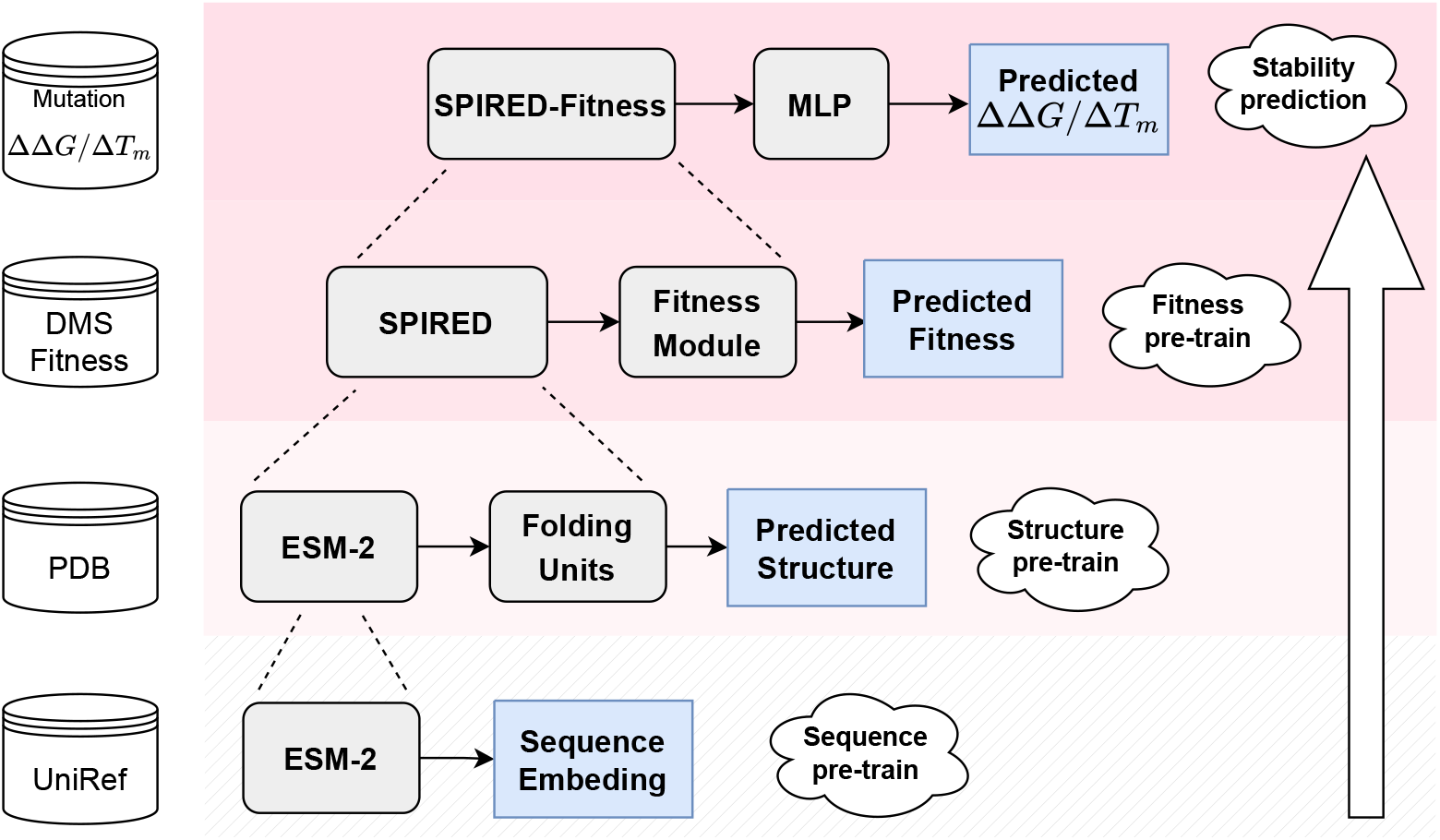
The Russian-doll-style pre-training strategy for SPIRED-Fitness and SPIRED-Stab. The step-by-step pre-training from sequence to structure, and then to fitness, and finally to stability. SPIRED can be regarded as the pre-trained structure model for SPIRED-Fitness, which itself is also the pre-trained model for SPIRED-Stab.

## Data availability

All the prediction results as well as the Dual Task Dataset are available at https://zenodo.org/records/10589086.

## Code availability

The models of SPIRED, SPIRED-Fitness and SPIRED-Stab are implemented in PyTorch^71^. All codes are freely downloadable at https://github.com/Gonglab-THU/SPIRED-Fitness. The model parameters are available at https://zenodo.org/records/10589086. The GDFold2 is our another work that is currently under preparation for publication. The codes of GDFold2 are available at https://github.com/Gonglab-THU/GDFold2.

The server of SPIRED-Fitness is available at http://structpred.life.tsinghua.edu.cn/server_spired_fitness.html. The server of SPIRED-Stab is available at http://structpred.life.tsinghua.edu.cn/server_spired_stab.html.

## Author contribution

H. Gong proposed the concept and theory. For the SPIRED model, Y. Chen, Y. Xing and H. Gong proposed the initial model design, Y. Chen and Y. Xing implemented coding and preliminary testing, and Y. Chen finalized the model details as well as the full model training. For the SPIRED-Fitness and SPIRED-Stab models, Y. Xu and H. Gong proposed the model architecture, Y. Xu and Y. Chen implemented model training and testing, and Y. Xu, Y. Chen and D. Liu analyzed the results. Y. Chen, Y. Xu, D. Liu and H. Gong wrote the manuscript. All authors agreed with the final manuscript.

## Supplementary Materials

### 1 Supplementary Results for Structure Prediction

#### 1.1 Sequentially arranged Folding Units incrementally enhance the accuracy of structure prediction

**Figure S1.**
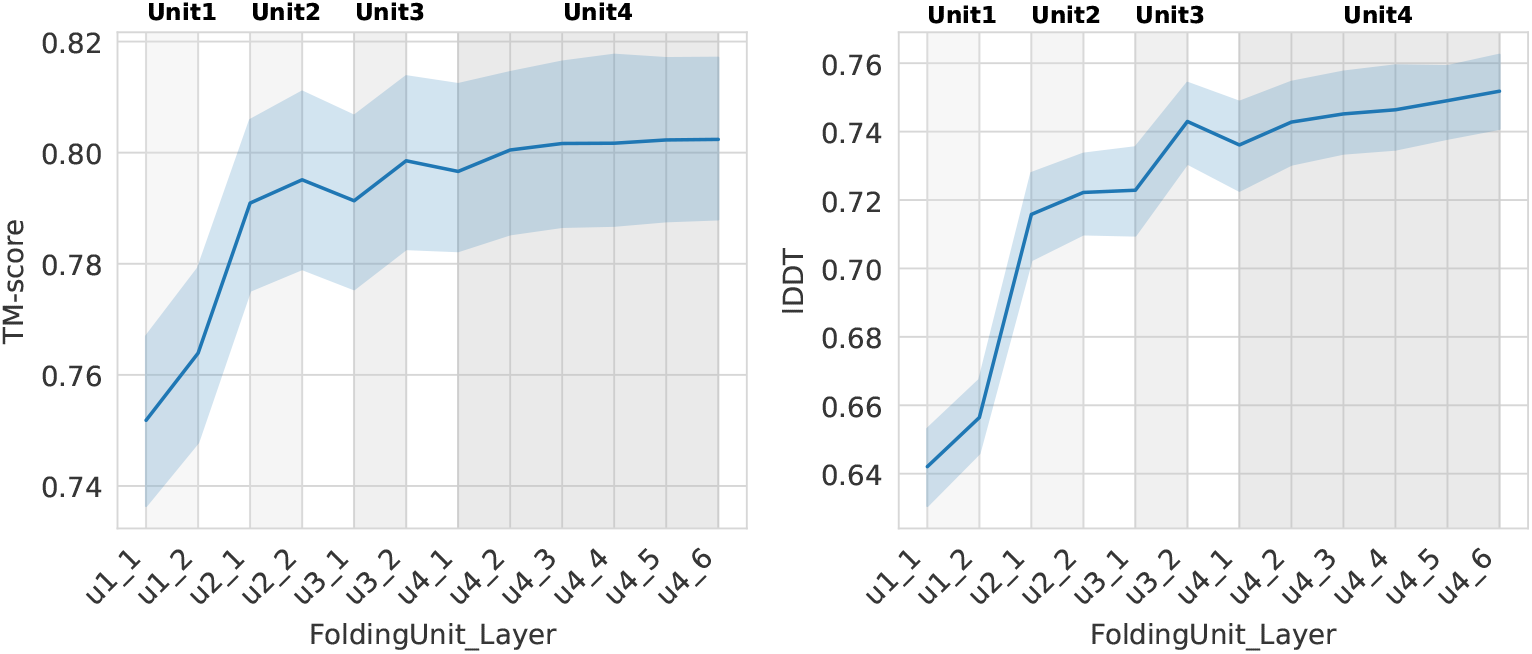
Evolution of TM-score and lDDT for coordinates generated by sequentially arranged Folding Units. The horizontal axis displays the structure prediction layers (*i*.*e*. Pred-XYZ modules) of sequentially arranged Folding Units. For example, u4_6 means the 6^*th*^ Pred-XYZ module of the 4^*th*^ Folding Unit. The vertical axis shows the TM-score or lDDT values of the generated C_*α*_ coordinates. The line connects the average TM-score/lDDT values for the outputs of sequential layers. The blue shadow shows the distribution of TM-score/lDDT values for among 568 CAMEO samples. The gray shadows mark different Folding Units.

We use SPIRED to predict structures for 568 proteins from the CAMEO set (Aug. 2022 ∼ Aug. 2023, 100 < *L* < 600). Analysis on the quality of the generated C_*α*_ coordinates by Pred-XYZ modules suggest a progressive optimization of predicted structures among the four Folding Units (Figure S1). Specifically, the average TM-score of generated C_*α*_ coordinates increases from 0.752 for the 1^*st*^ layer (*i*.*e*. Pred-XYZ module) of the 1^*st*^ Folding Unit to 0.802 for the 6^*th*^ layer of the 4^*th*^ Folding Unit. Similarly, the lDDT increases from 0.642 to 0.752. This trends indicates that the sequentially arranged Folding Units play a similar role to the recurrent expansion by recycling, both of which are capable of gradually optimizing the accuracy of the predicted structures.

#### 1.2 Performance of structure prediction models on CAMEO and CASP15 targets with 4 recycling cycles

**Figure S2.**
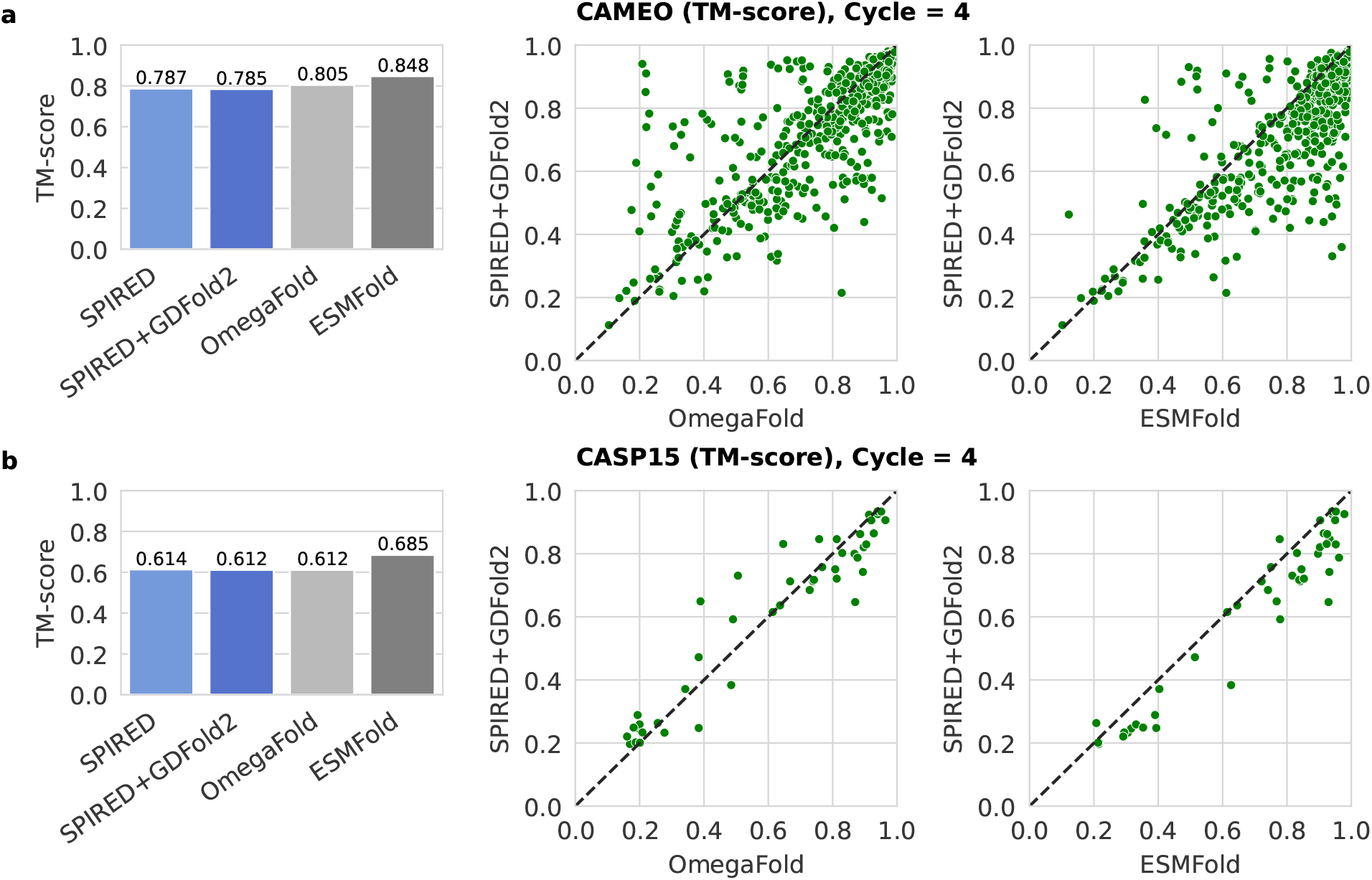
Comparison of model performance on CAMEO and CASP15. **a)** The barplot displays the TM-scores of predicted structures by SPIRED, SPIRED+GDFold2, OmegaFold, and ESMFold on the CAMEO set (680 proteins) with Cycle = 4. In the scatter plot, each point represents a protein, with the vertical axis representing the SPIRED+GDFold2 prediction and the horizontal axis representing the results of OmegaFold and ESMFold, respectively. **b)** Similar evaluation of models on the CASP15 set (45 protein domains) with Cycle = 4. All models make predictions from the domain sequences released by the CASP15 official website. The TM-score is calculated by comparing the predicted structures with the released CASP15 domain structures.

SPIRED+GDFold2, OmegaFold, and ESMFold2 are evaluated with recycling at Cycle = 4. As shown in the bar plots of Figure S2, ESMFold2 demonstrates the best performance on both CAMEO and CASP15 targets, while SPIRED+GDFold2 and OmegaFold2 are comparable to each other. The scatter plots show the significant differences between SPIRED and OmegaFold for different individual proteins, thus indicating the possible complementarity between different models.

#### 1.3 Case study for CAMEO, CASP15, SCOPe and CATH targets

**Figure S3.**
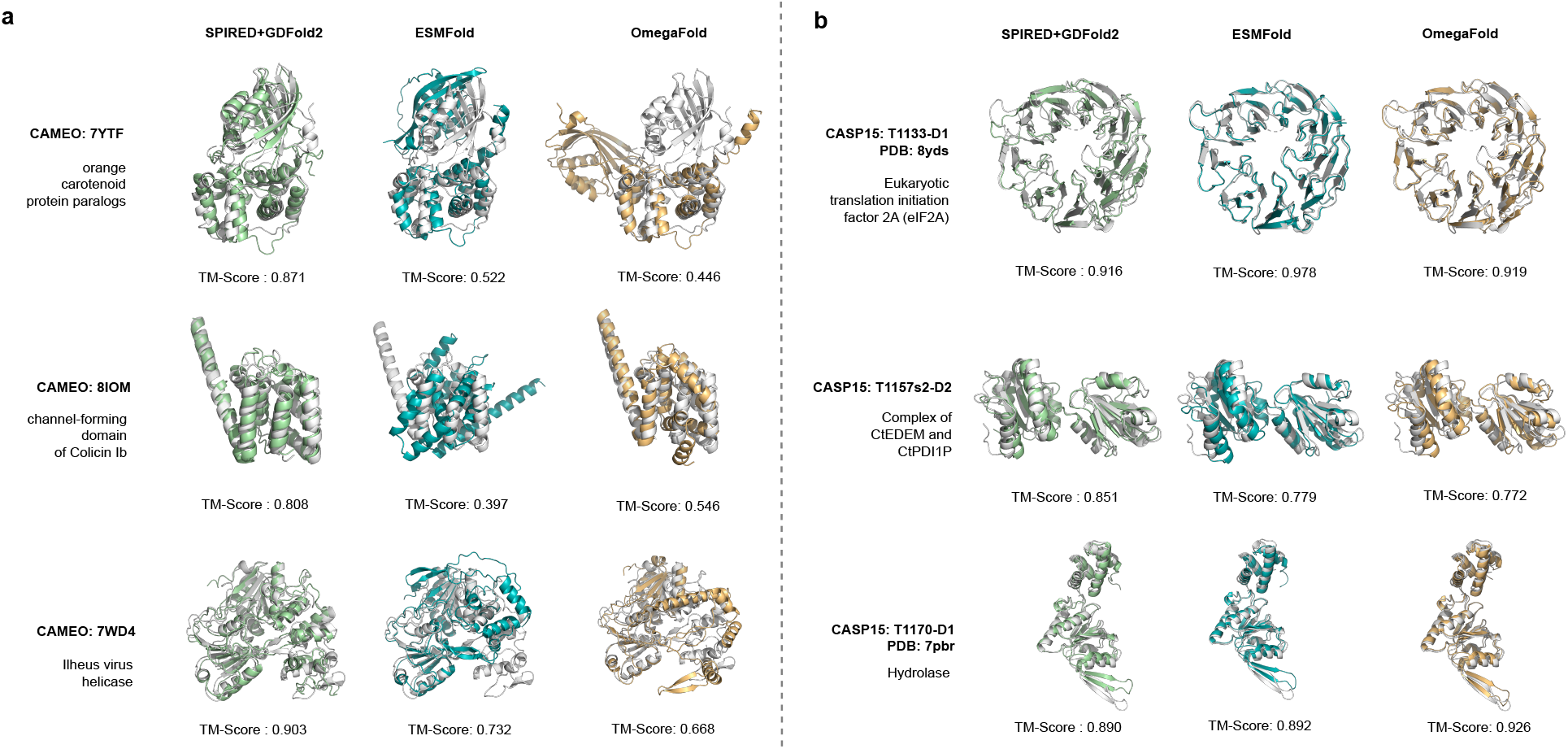
Case study for the CAMEO and CASP15 sets.

**Figure S4.**
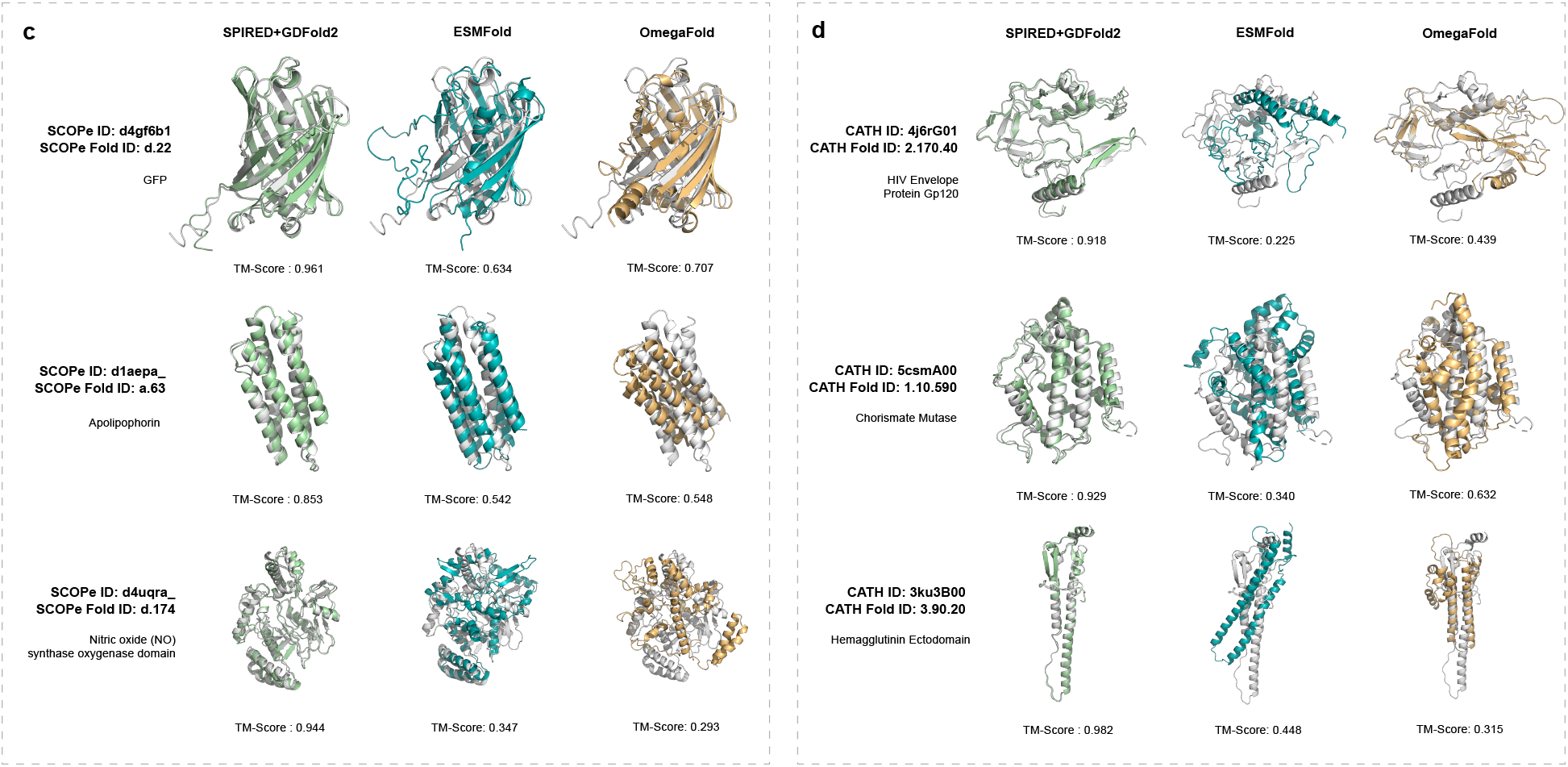
Case study for the SCOPe and CATH sets. The predicted structures of SPIRED+GDFold2 are represented in green, ESMFold’s predictions are represented in teal, OmegaFold’s predictions are colored in orange, and the ground-truth structures are colored in silver.

Structure prediction cases of SPIRED are shown in Figure S3 for the CAMEO and CASP15 sets, and are shown in Figure S4 for the SCOPe and CATH sets. SPIRED demonstrates predictive advantage on certain proteins of all alpha type, such as CAMEO: 8IOM, SCOPe: d1aepa_ (Apolipophorin), and CATH ID: 5csmA00 (Chorismate Mutase). Furthermore, SPIRED also shows advantage on GFP (SCOPe Fold ID: d.22), an extensively used target in protein design and engineering.

#### 1.4 Evaluation of structure prediction at the level of CATH topologies

**Figure S5.**
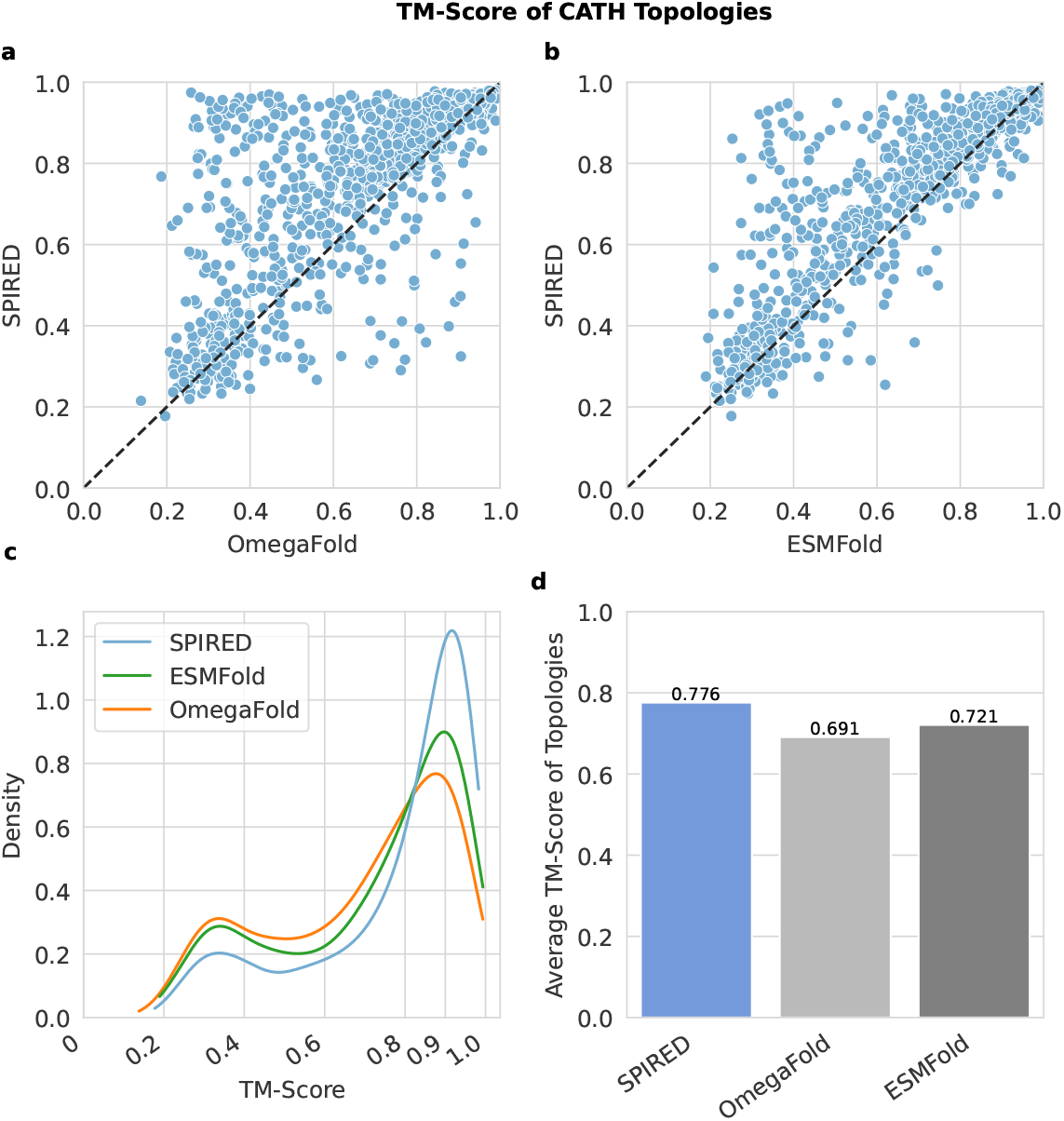
Evaluation of structure prediction on the CATH v4.2 S35 structural classification database. **a-b)** In the scatter plot, each point represents the average TM-score of all domains of one CATH topology. The veritcal axis is the prediction performance (TM-score) of SPIRED on different folds, while the horizontal axis represents results for OmegaFold and ESMFold, respectively. **c)** The KDE plot is used to visualize the distribution of model performance (in TM-score) over different folds for the tested models. **d)** The bar plot compares the TM-score averaged over all CATH topologies for different models.

CATH splits experimentally determined protein structures into domains, which are classified into 4 hierarchies: class (C), architecture (A), topology/fold (T) and homologous superfamily (H). We analyze the performance of structure prediction models on 1,223 topologies (consisting of 24,183 domains) from the CATH database (v4.2, S35, July 2017) in Figure S5. Among the topologies in which SPIRED performs better, there are 96 CATH topologies for which SPIRED achieve an advantage > 0.2 in TM-score over ESMFold, whereas ESMFold outperforms SPIRED by > 0.2 in TM-score for only 5 CATH topologies (Figure S5a,b). According to the KDE plot, SPIRED has the highest density in the high accuracy region (TM-score =∼0.9) (Figure S5c). SPIRED also reaches the highest mean TM-score when averaged over all CATH topologies (Figure S5d). Therefore, SPIRED exhibits similar advantageous performance on the CATH database as it does on SCOPe. Although OmegaFold and ESMFold do not explicitly use the CATH database as a training set, the majority of structures in CATH database are included in the PDB database they utilize. Nevertheless, the above results in combination with analysis over the SCOPe database (Figure 3) indicate that SPIRED is capable of learning the sequence-structure relationship from a wider range of topology/fold types than ESMFold and OmegaFold.

### 2 Supplementary Results for Fitness and Stability Prediction

#### 2.1 End-to-end training improves fitness prediction for SPIRED-Fitness

**Table S1.** Performance of the fitness and structure prediction on the test set.

| Model | Fitness Test Set |  |  | CAMEO |  |
| --- | --- | --- | --- | --- | --- |
| | $\rho_{\text{single and double}}$ | $\rho_{\text{single}}$ | $\rho_{\text{double}}$ | TM-score | IDDT |
| **SPIRED** | N.A. | N.A. | N.A. | **0.786** | 0.753 |
| **SPIRED+GDFold2** | N.A. | N.A. | N.A. | 0.784 | **0.756** |
| **SPIRED-Fitness (Stage 1)** | 0.830 | 0.816 | 0.766 | **0.786** | 0.753 |
| **SPIRED-Fitness (Stage 2)** | **0.850** | **0.837** | **0.795** | 0.779 | 0.740 |
| **SPIRED-Fitness + GDFold2** | **0.850** | **0.837** | **0.795** | 0.777 | 0.743 |

The best performance in each metric is highlighted in bold. Here, $\rho_{\text{single and double}}$ refers to the average Spearman correlation coefficients evaluated on all single and double mutations of proteins in the cDNA proteolysis dataset, whereas $\rho_{\text{single}}$ and $\rho_{\text{double}}$ stand for the corresponding value for single and double mutations, respectively.

We use the fitness testing dataset (485 proteins) described in **Methods** to evaluate the performance of fitness prediction. The CAMEO dataset (680 proteins) is utilized to assess the model performance on structure prediction without recycling (*i*.*e*. Cycle = 1). As shown in Table S1, after the end-to-end training, quality of the structural models provided by SPIRED-Fitness (Stage 2) drops marginally in comparison to the original SPIRED, which indicates that the end-to-end training process is not biased by the limited number of proteins in the fitness training set.

As described in **Methods**, in the first training stage of SPIRED-Fitness, parameters of SPIRED are frozen, and only the Fitness module is updated. At the end of the first training stage, the Spearman correlation coefficient on the union of single and double mutations in the fitness testing dataset reaches 0.830. After the completion of the second training stage of SPIRED-Fitness, where both SPIRED and the Fitness module are allowed to update their parameters, the Spearman correlation coefficient on the union of single and double mutations increases to 0.850. Regarding single mutations, the rise in Spearman correlation coefficient is approximately 2%, while for double mutations, it is about 3%. The TM-score of structure prediction for SPIRED-Fitness only decreases by < 0.01 compared to SPIRED, and the lDDT decreases by 0.013. Therefore, the end-to-end training has minimal impact on the structure prediction performance of SPIRED-Fitness. Considering the improvement in fitness prediction, this is an acceptable trade-off.

#### 2.2 Performance comparison of SPIRED-Fitness and other models

**Figure S6.**
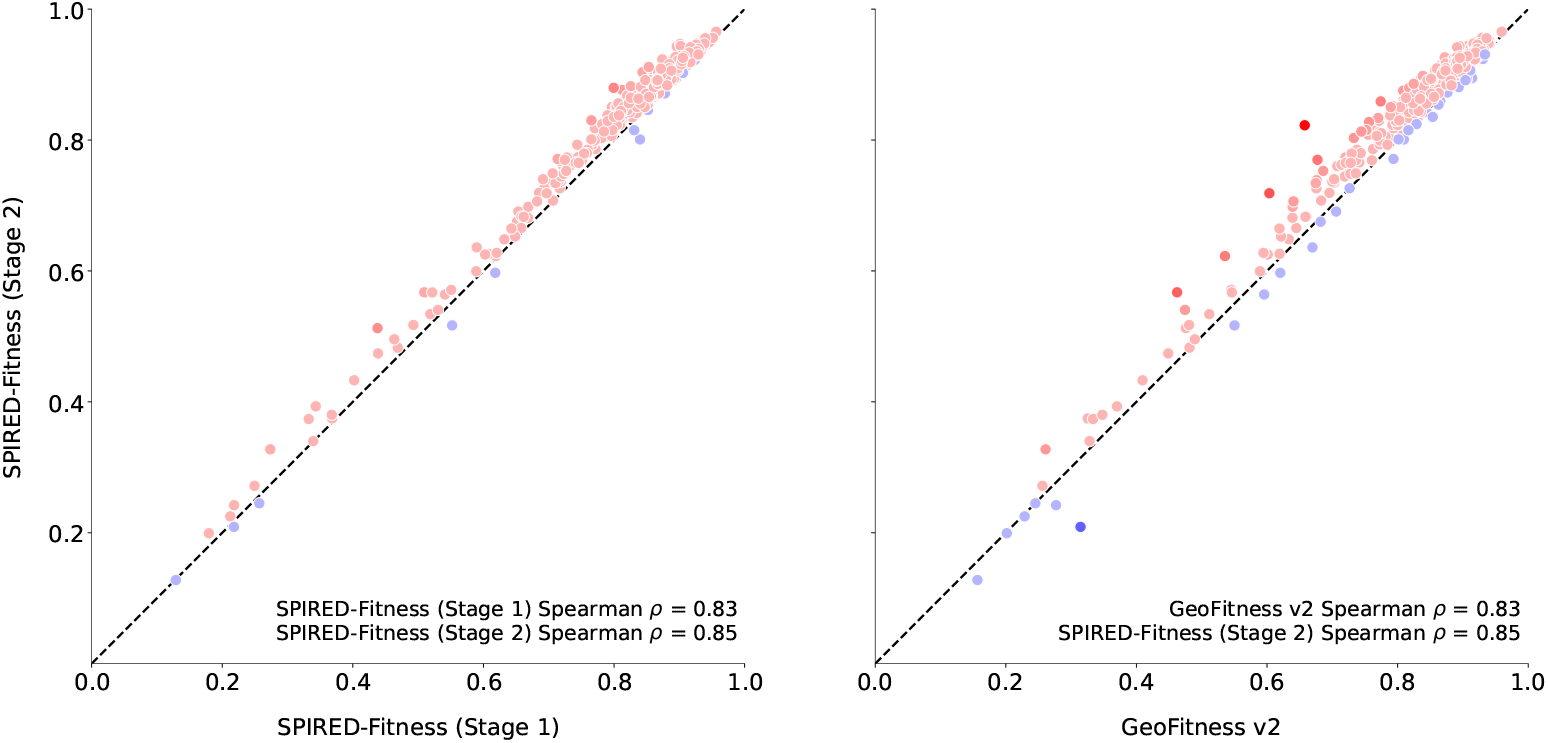
Protein-wise comparison between SPIRED-Fitness and GeoFitness on single and double mutations. The vertical axis represents SPIRED-Fitness (Stage 2), while the horizontal axis stands for SPIRED-Fitness (Stage 1) and GeoFitness v2. Each data point represents the Spearman correlation coefficient of the corresponding protein in the dataset. Data points are colored red if the Spearman correlation coefficient of SPIRED-Fitness (Stage 2) is higher than that of the other models; otherwise, points are colored blue. The test set comprises a total of 485 proteins.

The network architecture of the Fitness module of SPIRED-Fitness is almost identical to that of GeoFitness v2. Therefore, the principal difference between SPIRE-Fitness (Stage 1) and GeoFitness v2 lies in the module or model used for generating the structural features: SPIRED in SPIRED-Fitness (Stage 1) vs. AlphaFold2 in GeoFitness v2. Considering these facts, the advantage of SPIRED-Fitness (Stage 2) over both SPIRED-Fitness (Stage 1) and GeoFitness v2 demonstrates the positive effect of the end-to-end training on the performance of fitness prediction models.

**Figure S7.**
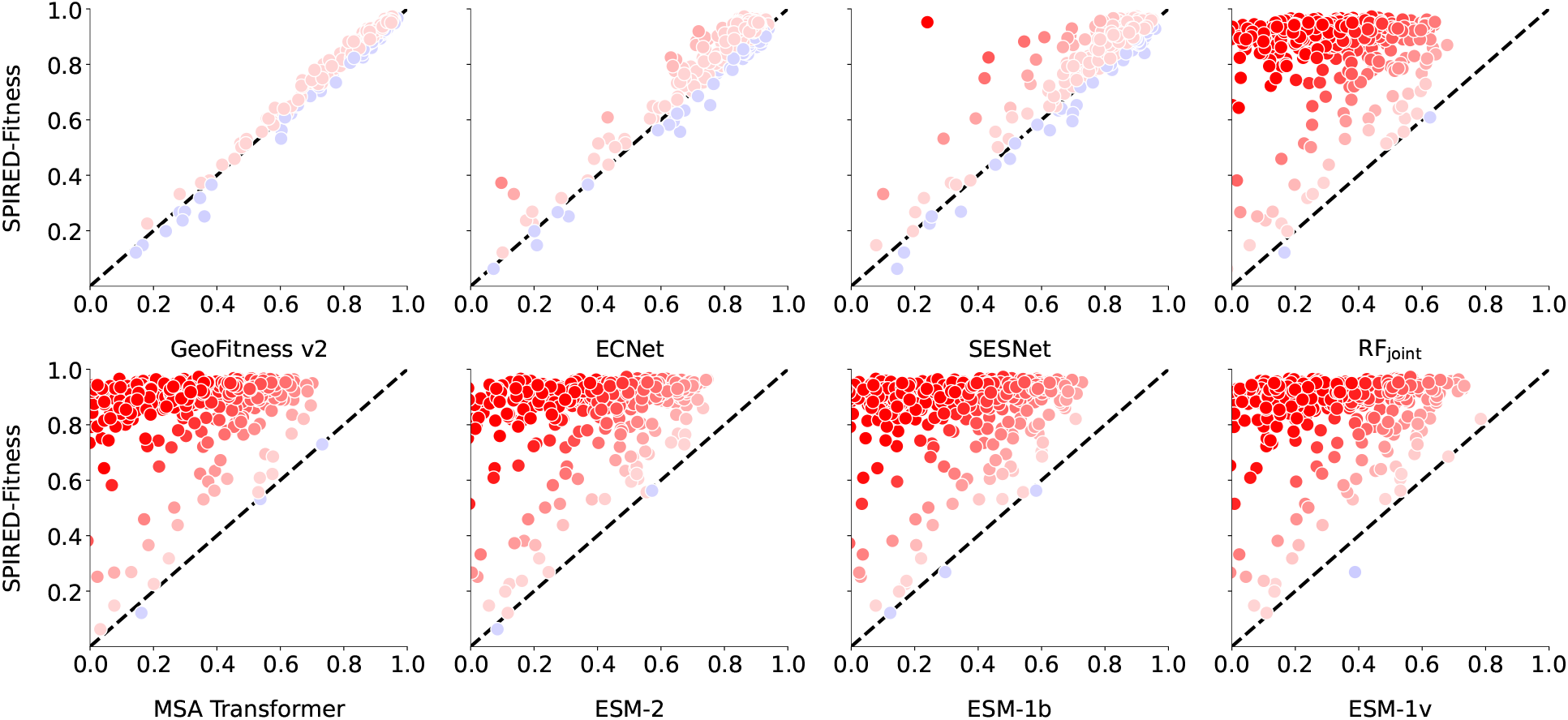
Protein-wise comparison of SPIRED-Fitness with other models on single mutations. The vertical axis represents SPIRED-Fitness, while the horizontal axis stands for different models. Each data point represents the Spearman correlation coefficient of the corresponding protein in the dataset. Data points are colored red if the Spearman correlation coefficient of SPIRED-Fitness is higher than that of the other models, but are colored blue otherwise. The test set comprises a total of 485 proteins.

#### 2.3 Supplementary results for the SPIRED-Stab evaluation

**Table S2.** Comparison of SPIRED-Stab with other ΔΔ*G* predictors on the S461 dataset.

| Method | $\rho$ | $\tau$ | $r$ | Top-5 | Top-10 |
| --- | --- | --- | --- | --- | --- |
| *Structure-based* |  |  |  |  |  |
| SDM | 0.32 | 0.24 | 0.36 | 0.40 | 0.58 |
| ThermoNet | 0.40 | 0.29 | 0.50 | 0.35 | 0.55 |
| mCSM | 0.39 | 0.29 | 0.44 | 0.40 | 0.56 |
| Dynamut | 0.39 | 0.28 | 0.36 | 0.32 | 0.53 |
| I-Mutant3.0 | 0.40 | 0.29 | 0.44 | 0.38 | 0.56 |
| MAESTRO | 0.49 | 0.36 | 0.52 | 0.42 | 0.60 |
| DUET | 0.45 | 0.33 | 0.50 | 0.43 | 0.59 |
| ACDC-NN | 0.45 | 0.34 | 0.47 | 0.42 | 0.58 |
| FoldX | 0.42 | 0.31 | 0.41 | 0.38 | 0.57 |
| PopMusic | 0.51 | 0.38 | 0.54 | 0.45 | 0.65 |
| DDGun3D | 0.51 | 0.39 | 0.55 | 0.48 | 0.66 |
| PremPS | 0.51 | 0.39 | 0.56 | 0.43 | 0.63 |
| DDMut | 0.48 | 0.37 | 0.52 | 0.42 | 0.60 |
| INPS3D | 0.54 | 0.40 | 0.56 | 0.50 | 0.63 |
| GeoDDG-3D (old) | 0.59 | 0.44 | 0.59 | **0.62** | 0.71 |
| *Sequence-based* |  |  |  |  |  |
| MuPro | 0.23 | 0.17 | 0.24 | 0.30 | 0.52 |
| I-Mutant3.0-Seq | 0.26 | 0.19 | 0.30 | 0.32 | 0.53 |
| SAAFEC-Seq | 0.40 | 0.29 | 0.44 | 0.38 | 0.56 |
| ACDC-NN-Seq | 0.43 | 0.32 | 0.49 | 0.45 | 0.59 |
| INPS-Seq | 0.47 | 0.34 | 0.51 | 0.48 | 0.61 |
| DDGun | 0.52 | 0.39 | 0.52 | 0.43 | 0.58 |
| GeoDDG-Seq (old) | 0.60 | 0.46 | 0.63 | 0.55 | 0.70 |
| Mutate Everything (ESM-2) | 0.63 | 0.46 | 0.62 | 0.42 | 0.68 |
| SPIRED-Fitness (zero-shot) | 0.58 | 0.45 | 0.59 | 0.57 | 0.68 |
| GeoStab v2 | 0.68 | 0.54 | 0.67 | 0.60 | **0.76** |
| SPIRED-Stab | **0.71** | **0.57** | **0.69** | 0.60 | 0.73 |

The winner in each metric is highlighted in bold. All metrics are first calculated within each protein and then averaged over all proteins. Here, $\rho$ , $\tau$ and $r$ denote Spearman, Kendall and Pearson correlation coefficients, respectively, while Top-5 and Top-10 refer to the Top 5 precision and Top 10 precision, respectively, as defined in **Methods**.

##### 2.3.1 Evaluation of SPIRED-Stab using conventional metrics and in the cross-protein manner

**Table S3.** Comparison of SPIRED-Stab with existing models on the S461 dataset in the cross-protein manner.

| Method | Direct |  | Inverse |  | Antisymmetry |  |
| --- | --- | --- | --- | --- | --- | --- |
| | $\rho/r$ | RMSE/MAE | $\rho/r$ | RMSE/MAE | $r_{d-i}$ | $\langle \delta \rangle$ |
| *Structure-based* |  |  |  |  |  |  |
| **SDM** | 0.51/0.54 | 1.36/1.03 | 0.13/0.13 | 1.98/1.50 | -0.35 | 0.52 |
| **ThermoNet** | 0.47/0.53 | 1.26/0.95 | 0.42/0.50 | 1.33/1.03 | -0.86 | 0.16 |
| **mCSM** | 0.51/0.52 | 1.11/0.84 | 0.15/0.19 | 2.17/1.80 | -0.09 | 0.85 |
| **Dynamut** | 0.43/0.49 | 1.30/0.98 | 0.43/0.44 | 1.37/1.02 | -0.61 | 0.29 |
| **I-Mutant3.0** | 0.45/0.47 | 1.17/0.88 | 0.14/0.15 | 2.19/1.82 | -0.04 | 0.81 |
| **MAESTRO** | 0.58/0.61 | 1.09/0.81 | 0.16/0.19 | 1.93/1.54 | -0.22 | 0.68 |
| **DUET** | 0.56/0.58 | 1.10/0.81 | 0.18/0.20 | 1.97/1.59 | -0.15 | 0.72 |
| **ACDC-NN** | 0.56/0.60 | 1.09/0.79 | 0.55/0.59 | 1.10/0.80 | -0.98 | 0.04 |
| **FoldX** | 0.33/0.21 | 2.26/1.41 | 0.44/0.39 | 1.77/1.16 | -0.22 | 0.71 |
| **PopMusic** | 0.58/0.59 | 1.07/0.79 | 0.25/0.28 | 1.90/1.52 | -0.30 | 0.69 |
| **DDGun3D** | 0.57/0.62 | 1.14/0.82 | 0.54/0.57 | 1.21/0.87 | -0.96 | 0.10 |
| **PremPS** | 0.57/0.60 | 1.08/0.83 | 0.56/0.60 | 1.04/0.77 | -0.86 | 0.20 |
| **DDMut** | 0.60/0.61 | 1.07/0.80 | 0.56/0.57 | 1.11/0.84 | -0.94 | 0.13 |
| **INPS3D** | 0.58/0.60 | 1.06/0.79 | 0.39/0.36 | 1.48/1.12 | -0.49 | 0.48 |
| **GeoDDG-3D (old)** | 0.65/0.65 | 1.05/0.75 | 0.65/0.65 | 1.05/0.75 | **-1.00** | **0.00** |
| *Sequence-based* |  |  |  |  |  |  |
| **MuPro** | 0.35/0.38 | 1.21/0.95 | 0.28/0.30 | 2.24/1.92 | -0.31 | 0.97 |
| **I-Mutant3.0-Seq** | 0.43/0.43 | 1.20/0.92 | 0.26/0.25 | 2.09/1.74 | -0.43 | 0.79 |
| **SAAFEC-Seq** | 0.45/0.47 | 1.17/0.88 | 0.02/0.03 | 2.29/1.91 | -0.05 | 0.85 |
| **ACDC-NN-Seq** | 0.53/0.56 | 1.13/0.82 | 0.53/0.56 | 1.13/0.82 | **-1.00** | **0.00** |
| **INPS-Seq** | 0.54/0.55 | 1.13/0.85 | 0.53/0.54 | 1.15/0.86 | **-1.00** | 0.03 |
| **DDGun** | 0.57/0.57 | 1.30/0.98 | 0.54/0.53 | 1.36/0.99 | -0.94 | 0.07 |
| **GeoDDG-Seq (old)** | 0.67/**0.67** | **1.03**/0.76 | 0.67/**0.67** | **1.03**/0.76 | **-1.00** | **0.00** |
| **Mutate Everything (ESM-2)** | 0.60/0.59 | 1.09/0.79 | -/- | -/- | - | - |
| **SPIRED-Fitness (zero-shot)** | 0.62/0.61 | 1.66/1.42 | -/- | -/- | - | - |
| **GeoStab v2** | 0.67/0.65 | 1.05/0.75 | 0.67/0.65 | 1.05/0.75 | **-1.00** | **0.00** |
| **SPIRED-Stab** | **0.68/0.67** | **1.03/0.72** | **0.68/0.67** | **1.03/0.72** | **-1.00** | **0.00** |

The winner in each metric is highlighted in bold. "Direct" refers to the mutations from the wild type to mutant, while "Inverse" means the corresponding mutations in the reverse direction. All metrics are calculated upon the data aggregated across the proteins and those denoted as "-" mean the model cannot provide results needed for the corresponding metric. Here, $\rho$ and $r$ denote Spearman and Pearson correlation coefficients, respectively, while RMSE and MAE refer to the root mean square error and mean absolute error, respectively. Definitions for the anti-symmetry metrics could be found in our prior work<sup>23</sup>. The raw results of the other models are taken from a few prior works<sup>23,44,52</sup>.

#### 2.4 Miscellany

##### 2.4.1 Impact of train-test-splitting strategy on the fitness prediction by SPIRED-Fitness

**Figure S8.**
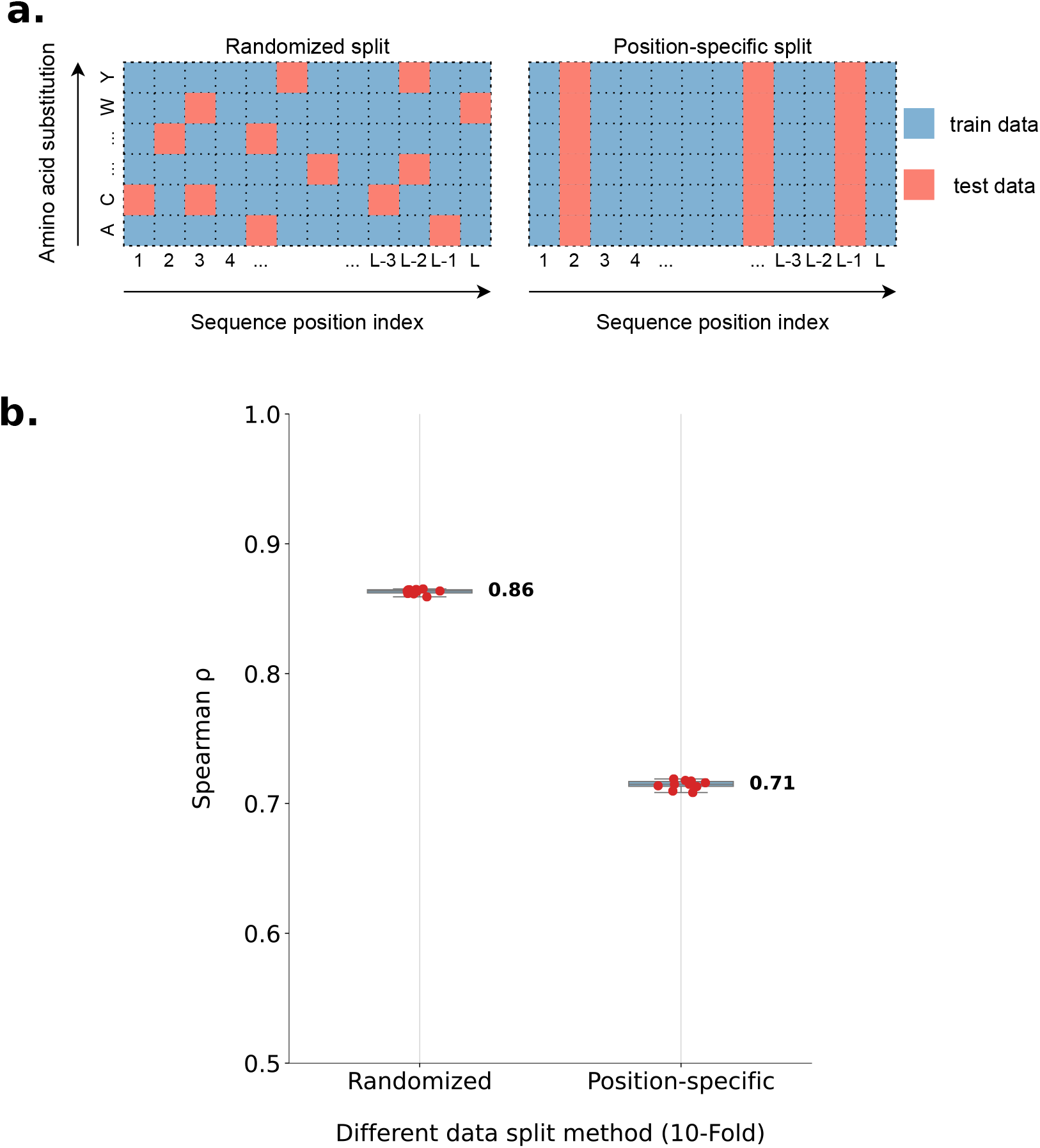
Performance evaluation of SPIRED-Fitness upon two data splitting strategies. **a)** Schematic representation of two data splitting strategies. **b)** Results of 10-fold cross validation on the two data splitting strategies. The horizontal axis represents different splitting strategies for DMS dataset. Each red dot in the graph represents the result of one 10-fold cross-validation, *i*.*e*. the mean Spearman correlation coefficient of all proteins. The median value of the boxplot is marked side.

We tested the fitness prediction performance of the SPIRED-Fitness model on three different fitness dataset splitting strategies (Figure S8 and Figure 5b). The first strategy randomly shuffles and splits all data with a ratio of training:validation:test = 7:1:2, while the second strategy splits data based on residue positions with the same ratio. The third strategy splits data based on proteins, with 70% proteins for training, 10% proteins for validation and 20% proteins for test. Given the above results, we chose the randomized splitting strategy for the final training of SPIRED-Fitness model. Results of the third strategy could be regarded as independent experiments for validating the model generalizability in the scenario of zero-shot prediction.

##### 2.4.2 Correlation between pLDDT and lDDT in the structure prediction by SPIRED/SPIRED-Fitness

**Figure S9.**
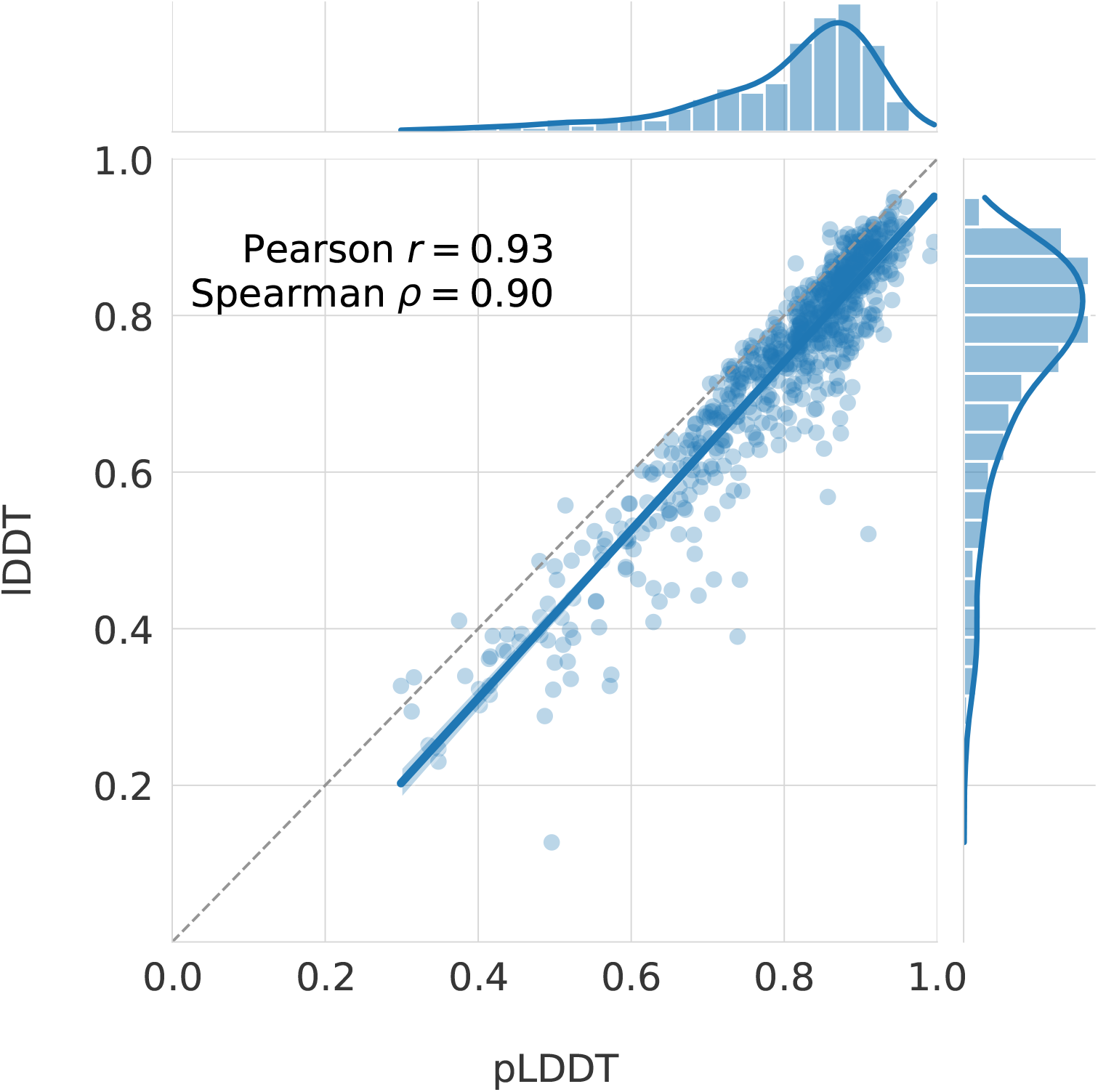
Correlation between pLDDT and lDDT for SPIRED-Fitness in the CAMEO dataset. The dataset is CAMEO test set (680 proteins, August 2022 to August 2023). Each data point represents a protein, with the average pLDDT (horizontal axis) and average lDDT (vertical axis) calculated based on the selection of the top 4 sets of C_*α*_ coordinates with the highest pLDDT values from the SPIRED-Fitness prediction.

For each protein of length *L*, SPIRED/SPIRED-Fitness can predict *L* sets of C_*α*_ coordinates, each with corresponding pLDDT values (a tensor of length *L*). For each protein target from the CAMEO and CASP15 set, among the *L* sets of protein coordinates predicted by SPIRED/SPIRED-Fitness, we select the top 4 sets of coordinates of the highest pLDDT values and compute the average TM-score as well as average lDDT of them for comparison with other models. As for the evaluation of structural prediction upon SCOPe and CATH targets, we compare the one set of coordinates predicted of the highest pLDDT value for comparison with other models.

We explore the correlation between lDDT and pLDDT values of the predicted structures from SPIRED-Fitness using the CAMEO dataset, by calculating the average pLDDT and lDDT for top 4 sets of the predicted coordinates of the highest pLDDT values. Figure S9 indicates the presence of a high Pearson correlation coefficient (0.93) and a high Spearman correlation coefficient (0.90) between the pLDDT and lDDT values for each protein, demonstrating that the pLDDT values output by SPIRED-Fitness are capable of effectively assessing the quality of predicted structures.

##### 2.4.3 Correlation between ΔΔ*G* and Δ*T*_*m*_ labels in the Dual Task Dataset of SPIRED-Stab

**Figure S10.**
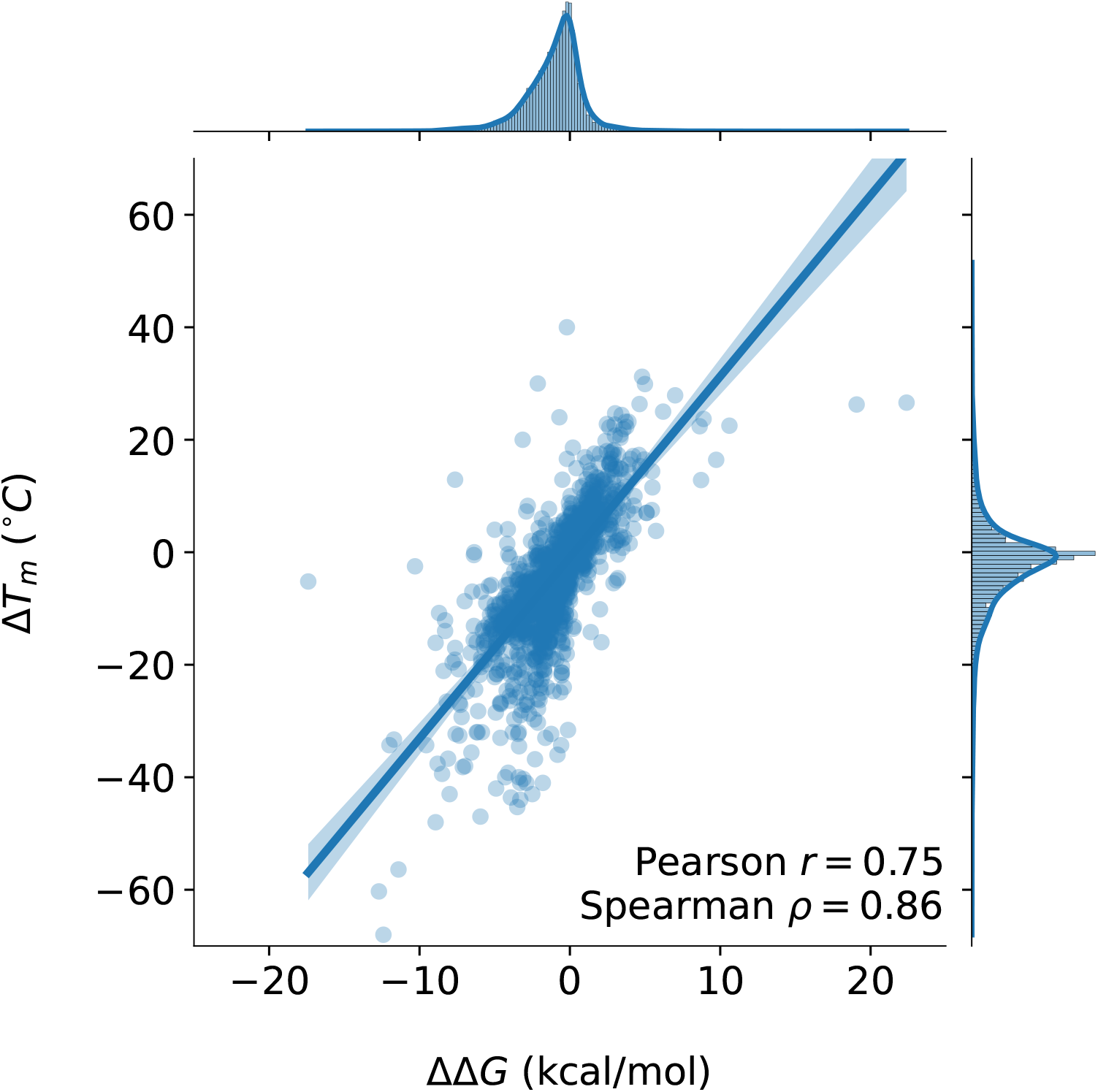
Correlation between ΔΔ*G* and Δ*T*_*m*_ in the Dual Task Dataset. Each point represents one entry in the Dual Task Dataset with both ΔΔ*G* and Δ*T*_*m*_ labels.

We evaluate the correlation between ΔΔ*G* and Δ*T*_*m*_ in the Dual Task Dataset constructed in this work. As shown in Figure S10, ΔΔ*G* and Δ*T*_*m*_ data present a strong correlation, with the Pearson correlation coefficient equal to 0.75, a phenomenon that is fully expected, since both metrics are descriptors of the protein stability, although ΔΔ*G* describes the resistance to denaturation at room temperature while Δ*T*_*m*_ reflects the resistance to thermal denaturation. Nevertheless, this observation provides supports for our dual task training strategy for SPIRED-Stab and Geo-Stab v2 in this work, *i*.*e*. optimization of the network parameters through the dual tasks of ΔΔ*G* and Δ*T*_*m*_ prediction in each epoch.

##### 2.4.4 Training details of GeoFitness v2 and GeoStab v2

We first trained the GeoFitness v2 using a similar training procedure to our prior work^23^ but with the additional training data of the cDNA proteolysis dataset. Notably, in this updated version, we slightly modified the output layer of the GeoFitness model to allow the prediction of double mutational effects, in a similar way to SPIRED-Fitness (see Figure 1d). Accordingly, during the model training, the Soft Spearman Loss was applied to both single and double mutations, as shown in Equation 1.

We then re-utilized the geometric encoder of GeoFitness v2 in the GeoStab v2 model and trained this model with a generally similar but slightly different procedure to our prior work^23^. Specifically, in the first stage, we completely fixed the parameters in the GeoFitness module and used the whole cDNA proteolysis dataset described in **Methods** as the training dataset to optimize the other parameters in the GeoStab v2 model except the ΔΔ*G*_*coef* and Δ*T*_*m*__*coef* parameters in the last output layer with the intial learning rate at 10^−3^. In the second stage, we further fine-tuned our model using the Dual Task Dataset newly collected in this work with the Soft Spearman Loss. The learning rate was assigned just as described for SPIRED-Stab in **Methods**. Finally, when the model had learned to capture the ranking information, we adopted the MSE loss to tune ΔΔ*G*_*coef* and Δ*T*_*m*__*coef* parameters in the last output layer with the initial learning rate of 10^−2^ in order to improve the model performance in predicting the absolute value of ΔΔ*G*/Δ*T*_*m*_.

### 3 Details for Structure Loss and Fitness Loss

#### 3.1 Loss for training SPIRED

The loss calculation method is similar for the first three Folding Units. Taking the Unit1_Loss of the first Folding Unit as an example, the Unit1_Loss is composed of RD_Loss, CaDist_Loss, pLDDT_Loss and Clash_Loss. Each Folding Unit predicts two groups of structures (by two Pred-XYZ modules), and we calculate the loss for each group of predicted structures separately, resulting in RD_Loss_1, RD_Loss_2, CaDist_Loss_1, and CaDist_Loss_2. The calculation method is based on Algorithm 11 and Algorithm 13. When the label of C_*α*_ distance is ≤ 24 Å, the RD_Loss and CaDist Loss are calculated and then normalized by the maximal distance of 24 Å. However, pLDDT_Loss and Clash_Loss are calculated only for the last predicted coordinates, following Algorithm 14 and Algorithm 15.

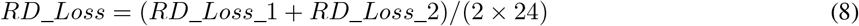

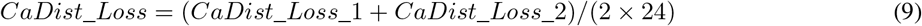

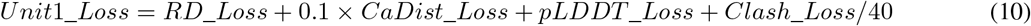

For Folding Unit4, the calculation of RD_Loss for the first 4 groups of predicted structures is based on Algorithm 11. When the label of C_*α*_ distance is ≤ 32 Å, the RD_Loss and CaDist Loss are calculated and then normalized by the maximal distance of 32 Å. When calculating the RD_Loss for the last 2 groups of predicted structures, Algorithm 12 is referenced, with a random selection of 10% of RD_Loss values that are greater than 40 into consideration of loss evaluation.

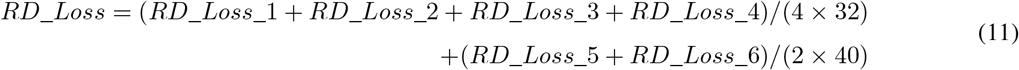

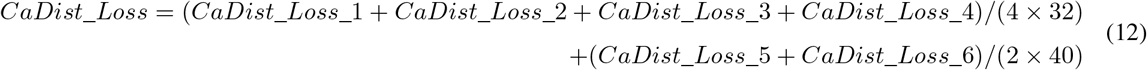

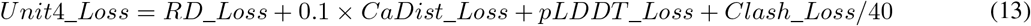

The total Loss is composed of four Folding Unit Losses, the Cross Entropy Loss of C_*β*_ distance distribution and that of dihedral/scalar angles (CE_Loss) as well as the L1 Loss of torsional angles (Torsion_Angle_Loss). *L* is the length of the protein. The hyper-paremeter *β* is 0.1 in the first two training stages shown in Table 1, and it is changed to 0.01 in the last two training stages.

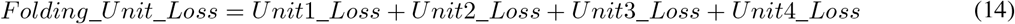

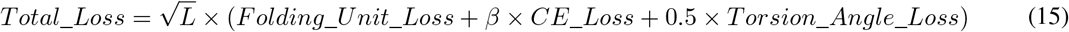

#### 3.2 Soft ranking loss for the differentiable Spearman correlation coefficient

The ordinary ranking operation is non-differentiable. To address this issue, we introduce the method proposed by Blondel *et al*.^41^ to achieve a differentiable Spearman correlation coefficient. First, the linear programming formulation for ranking can be derived after casting rankings onto the permutahedron. Second, the strong convex regularization (entropic regularization or quadratic regularization) is introduced into the linear programming formulation of ranking to smoothen discontinuous points of the ranking operators. Third, a parameter *ε* is introduced to adjust the regularization strength. Fourth, differentiation of ranking can be achieved by computing the Jacobian matrix of isotonic optimization of the projection onto the permutahedron. The authors found that excellent accuracy can be achieved in the ranking experiment without the need to adjust *ε*. In this study, we adopt entropic regularization and set *ε* to 0.01. The detailed mathematical derivation is described as follows.

In discrete ranking operations, ***θ*** := (*θ*_1_, …, *θ*_*n*_) is a vector of score, and ***ρ*** is reversing permutation for ***θ***. The *σ* refers to the permutation of [*n*], and the *π* is the reverse permutation of [*n*]. The Σ is the n! permutations for [*n*]. For all ***θ*** ∈ ℝ^*n*^, *π* ∈ Σ and = ***ρ*** := (*n, n* − 1, …, 1),

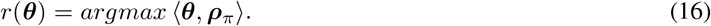

The permutahedron (***ω*** ∈ ℝ^*n*^) is introduced to achieve continuous and differentiable ranking operation. The convex hull of permutations of ***ω*** can be represented as:

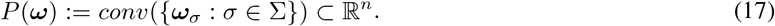

When ***ω*** = ***ρ***, the linear programming formulations of ranking can be described as below. For all ***θ*** ∈ ℝ^*n*^, ***ρ*** := (*n, n* − 1, …, 1) and *y* ∈ *P*(***ρ***),

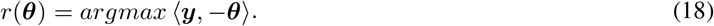

Setting (***z, ω***) = (-***θ, ρ***). Considering entropic regularization *E*(***µ***) := ⟨***µ***, *log****µ*** − 1⟩, Kullback-Leibler (KL) divergence is be introduced as

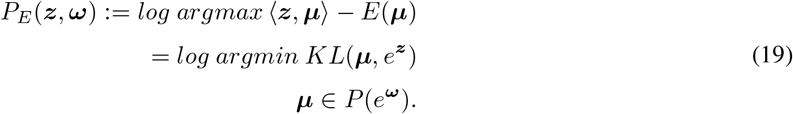

The convex regularization is defined as Ψ. The parameter *ε* > 0 is introduced to control the regularization strength. The Ψ-regularized soft rank can be defined as:

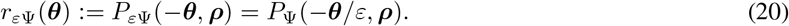

After reducing the projections to isotonic optimization, the differentiation of projections can be calculated by solving the Jacobian of the projections. For all *z* ∈ ℝ^*n*^ and sorted *ω* ∈ ℝ^*n*^:

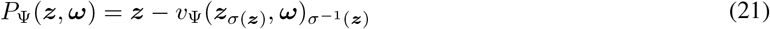

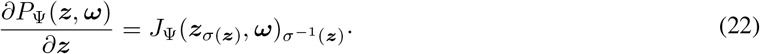

### 4 Algorithm and Pseudocode for SPIRED, SPIRED-Fitness and SPIRED-Stab

#### Algorithm for SPIRED

Algorithm 1 describes the overall approach of SPIRED (see Figure 1a). The 1D features are denoted as *s*, and the 2D features are denoted as *z*. Each Folding Unit processes and updates *s* and *z*, using *z* to predict C_*α*_ coordinates (*xyz*), inter-residue C_*α*_ distances (*Ca*_*dist*) and pLDDT. The embedding *z* from Folding Unit4 is utilized to predict C_*β*_ distance distribution as well as the dihedral angles (*ω* and *θ*) and scalar angles (φ) between residue pairs. The embedding *s* from Folding Unit4 is used to predict torsion angles (*ϕ* and *ψ*).

Details of the Folding Unit1 is illustrated in Algorithm 2 (see Figure 1b). The algorithms for Folding Unit2 and Folding Unit3 are similar, except that the input *z* for Folding Unit1 is a tensor of zeros, while the input *z* for Folding Unit2/3 comes from the previous Folding Unit. Folding Unit4 is more complex than the other ones and its implementation details are shown in Algorithm 3. The TriangularSelfAttentionBlock in these algorithms refers to the corresponding method, *i*.*e*. Triangular Self-Attention, in the ESMFold^9^.

##### Algorithm 1 SPIRED

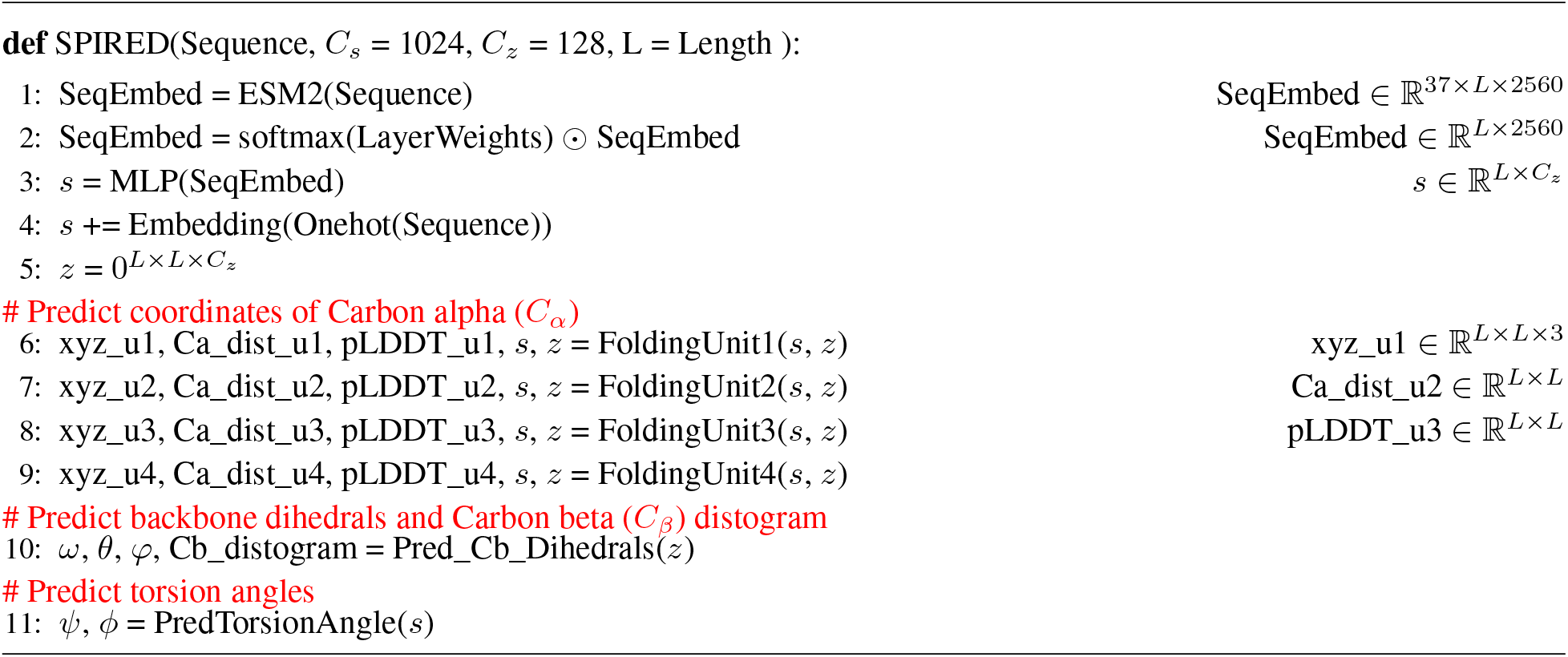

##### Algorithm 2 FoldingUnit1

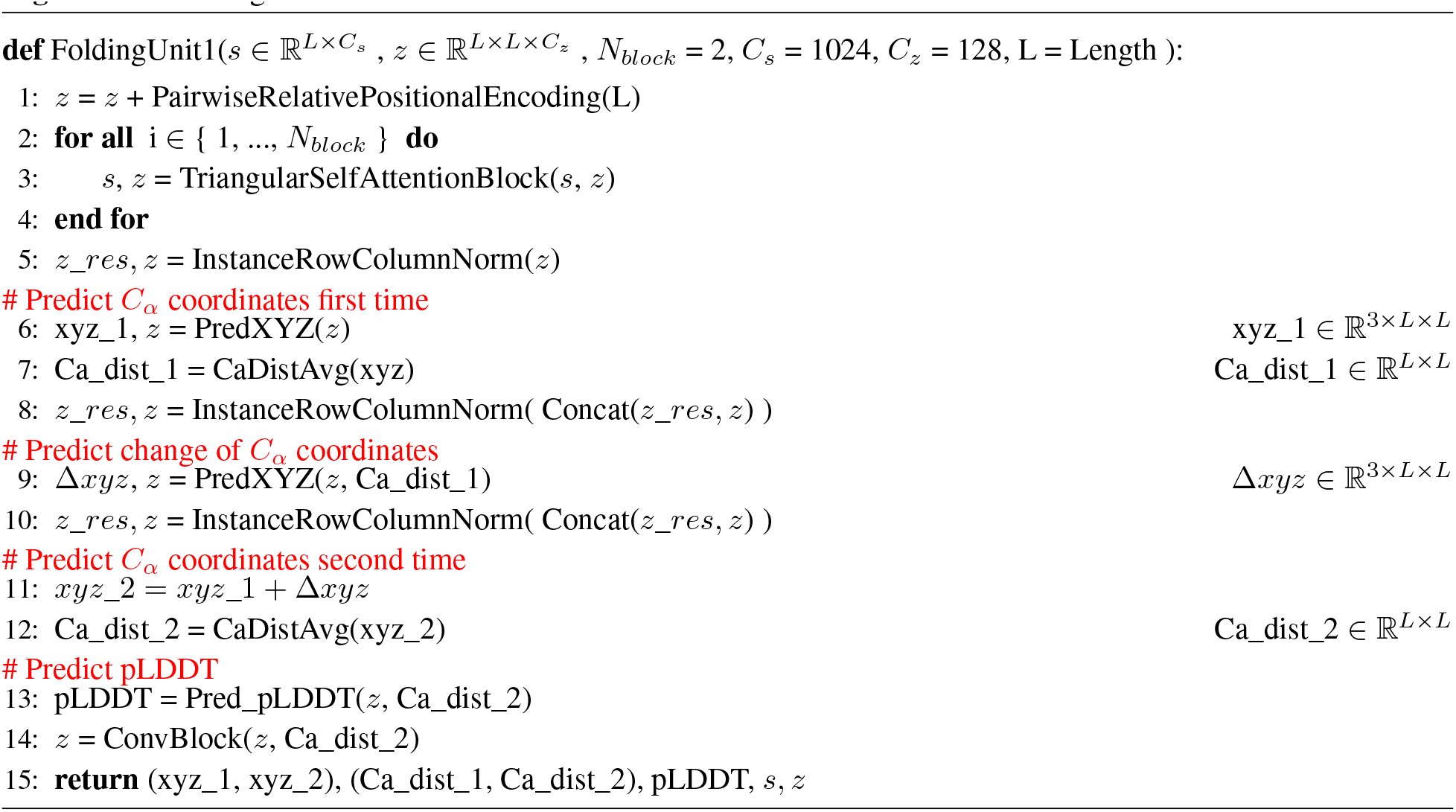

##### Algorithm 3 FoldingUnit4

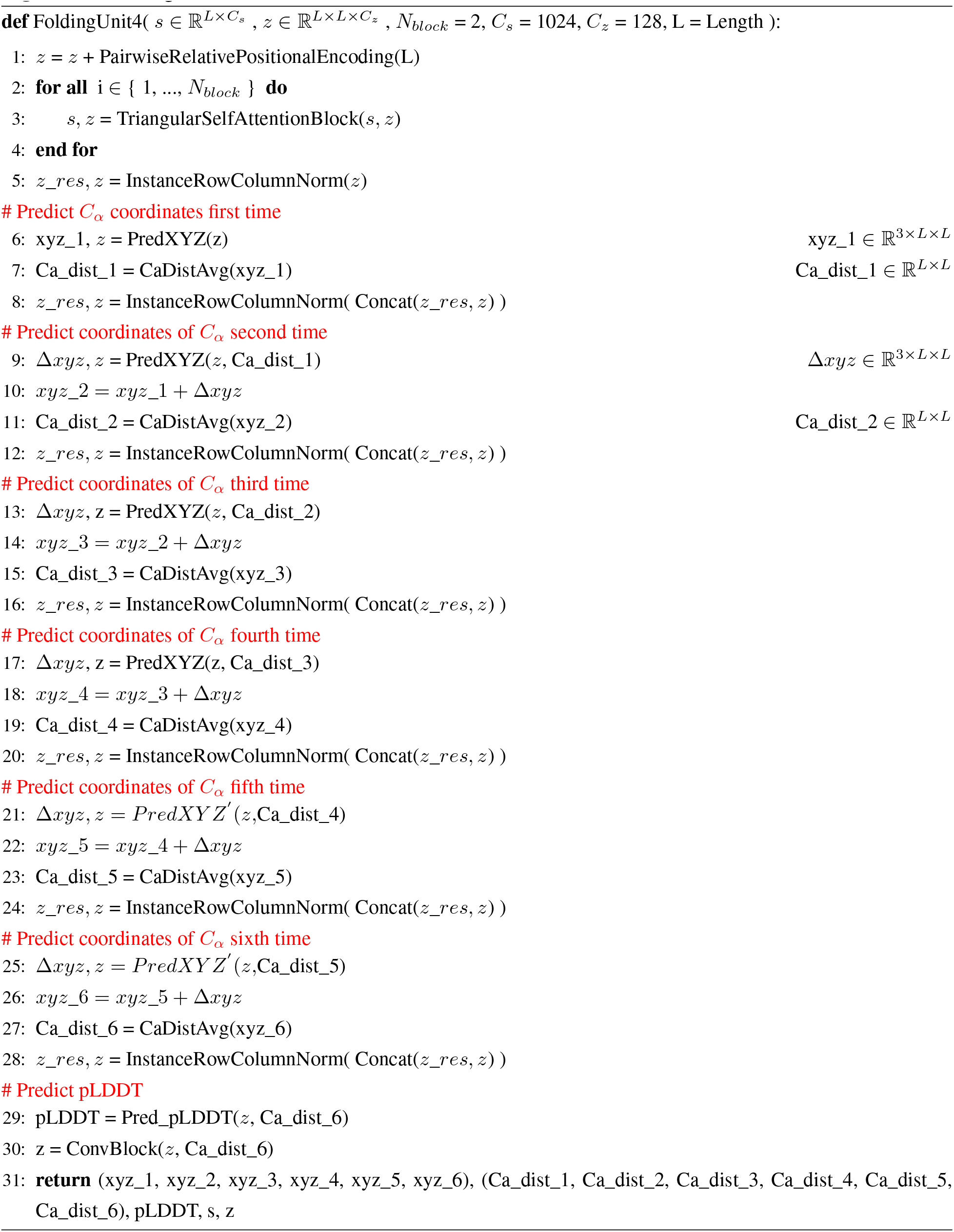

##### Algorithm 4 InstanceRowColumnNorm

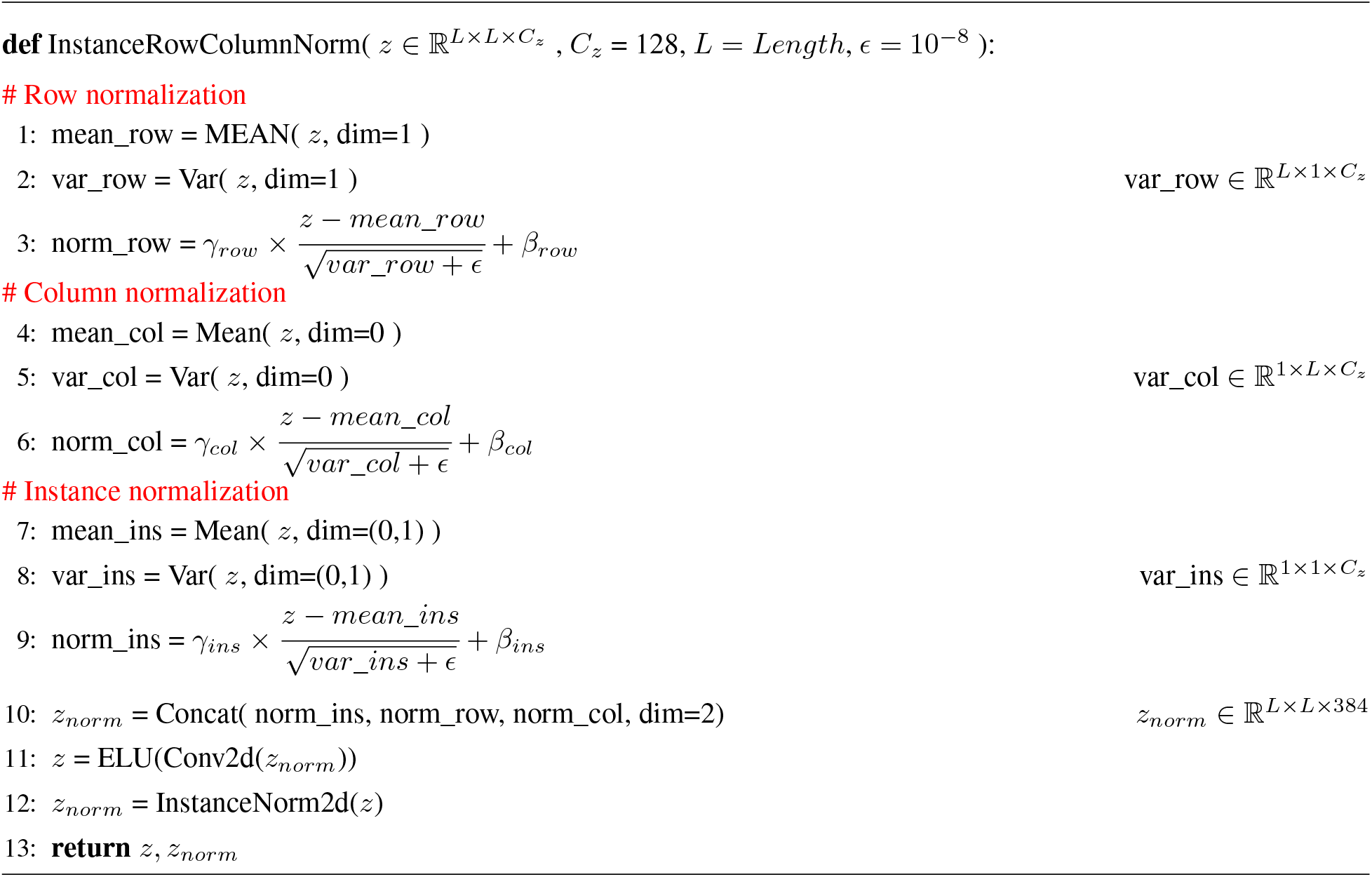

##### Algorithm 5 PredXYZ

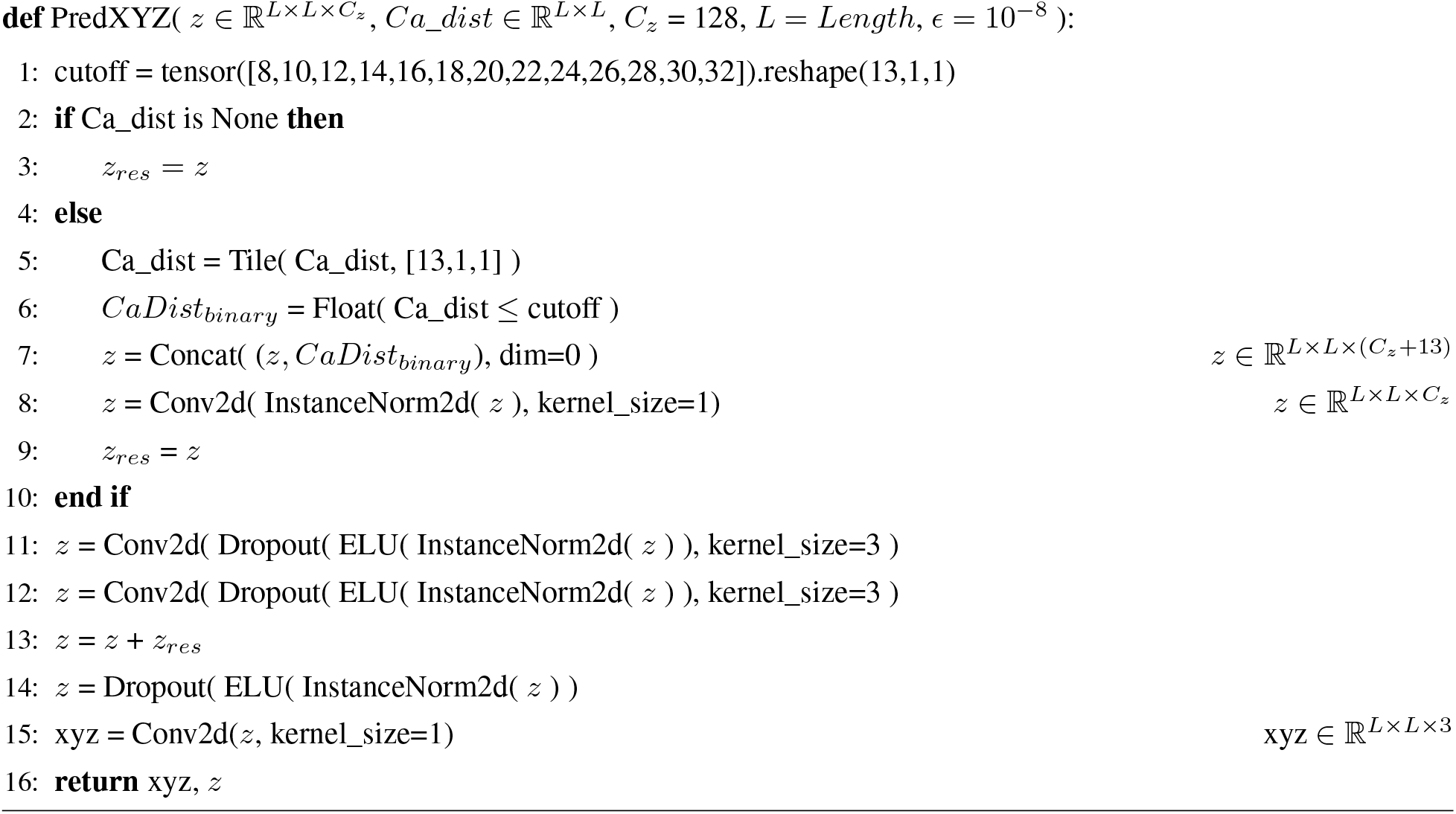

##### Algorithm 6 CaDistAvg

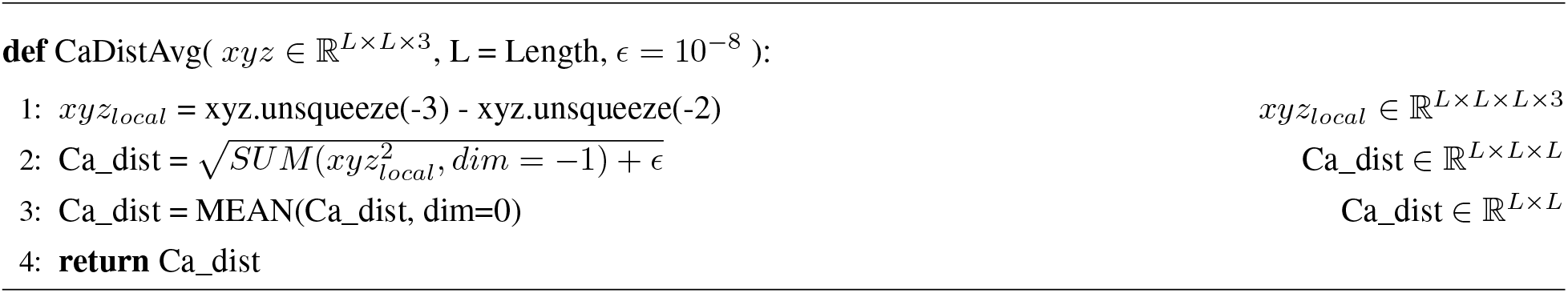

##### Algorithm 7 ConvBlock

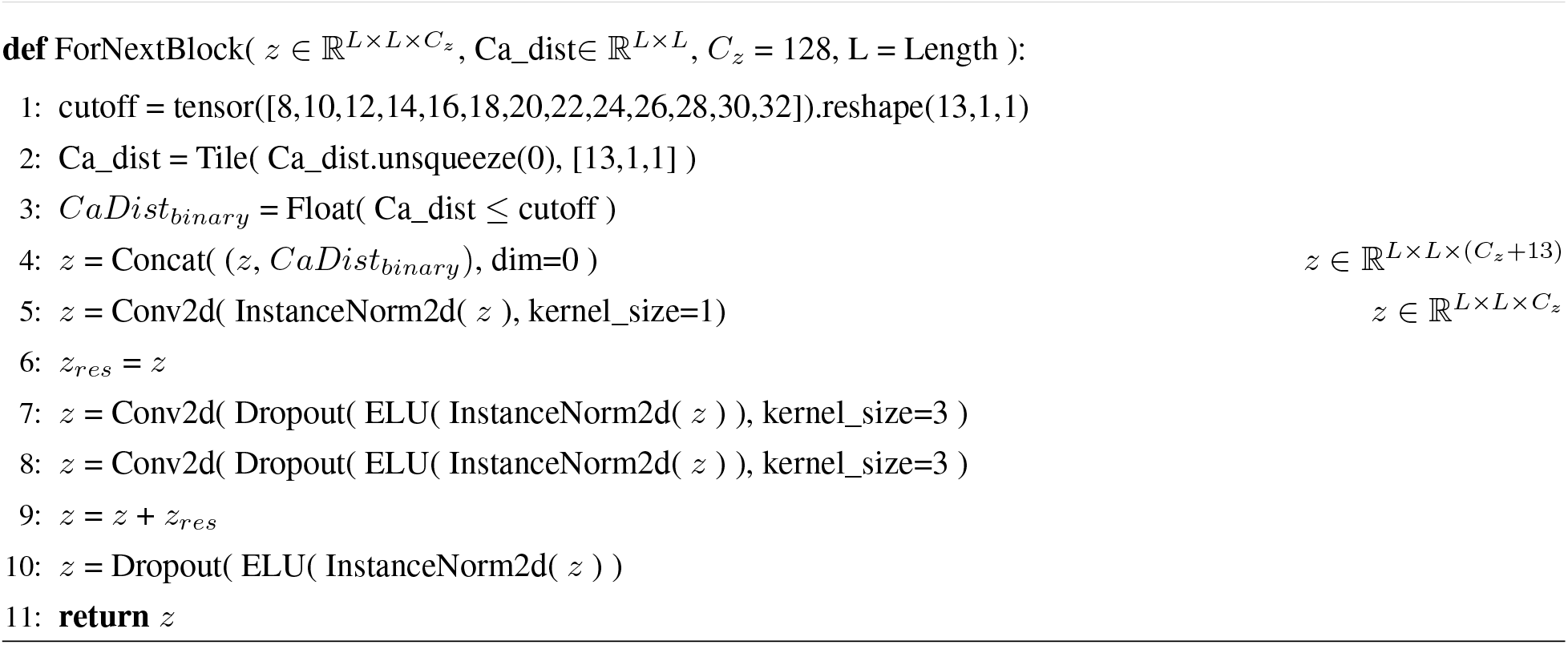

##### Algorithm 8 Pred_Cb_Dihedrals

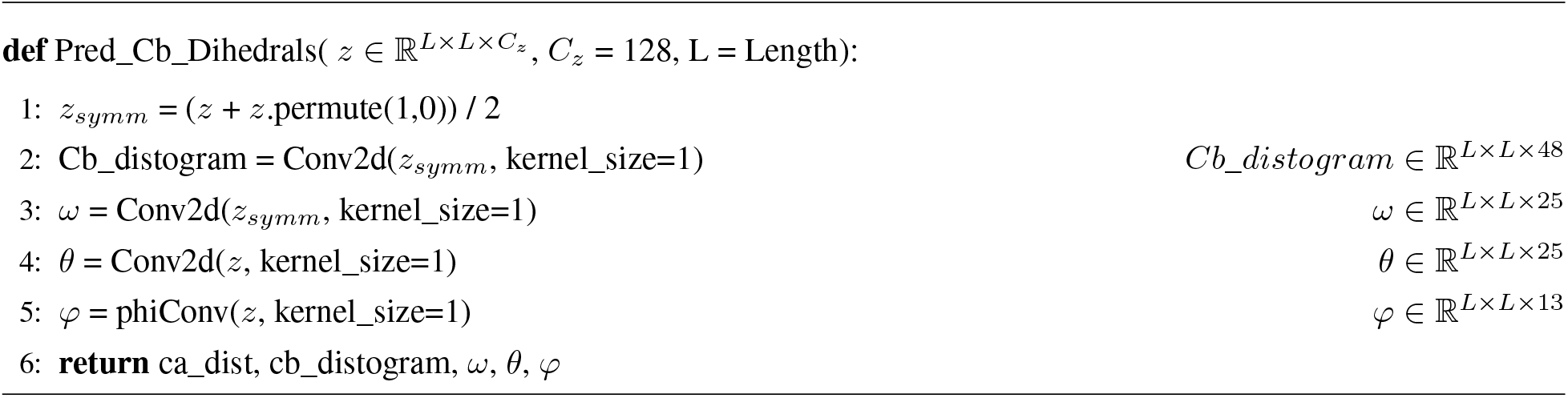

##### Algorithm 9 Pred_TorsionAngle

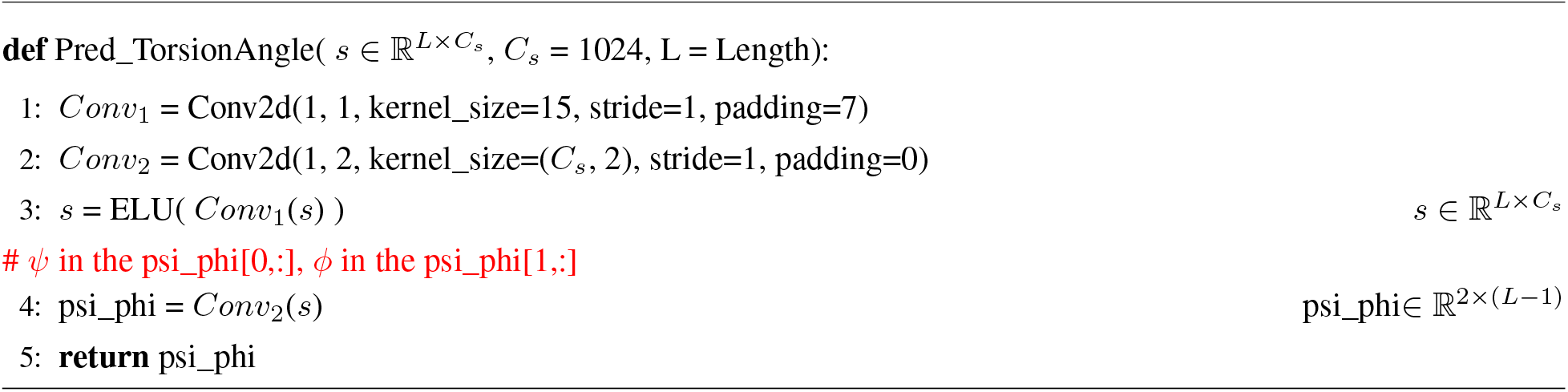

##### Algorithm 10 Pred_pLDDT

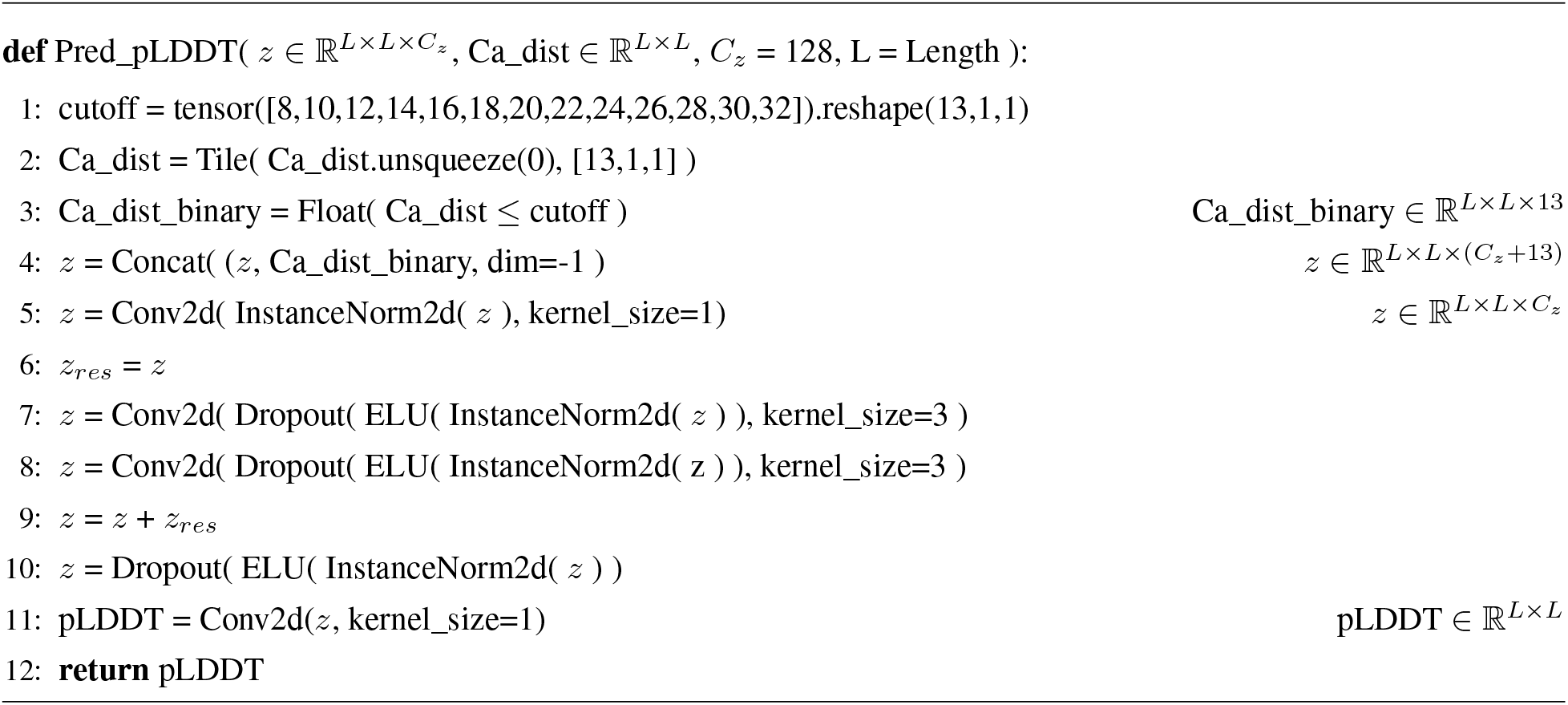

#### 4.2 Algorithm for the loss functions of SPIRED training

##### Algorithm 11 compute_RDloss_1

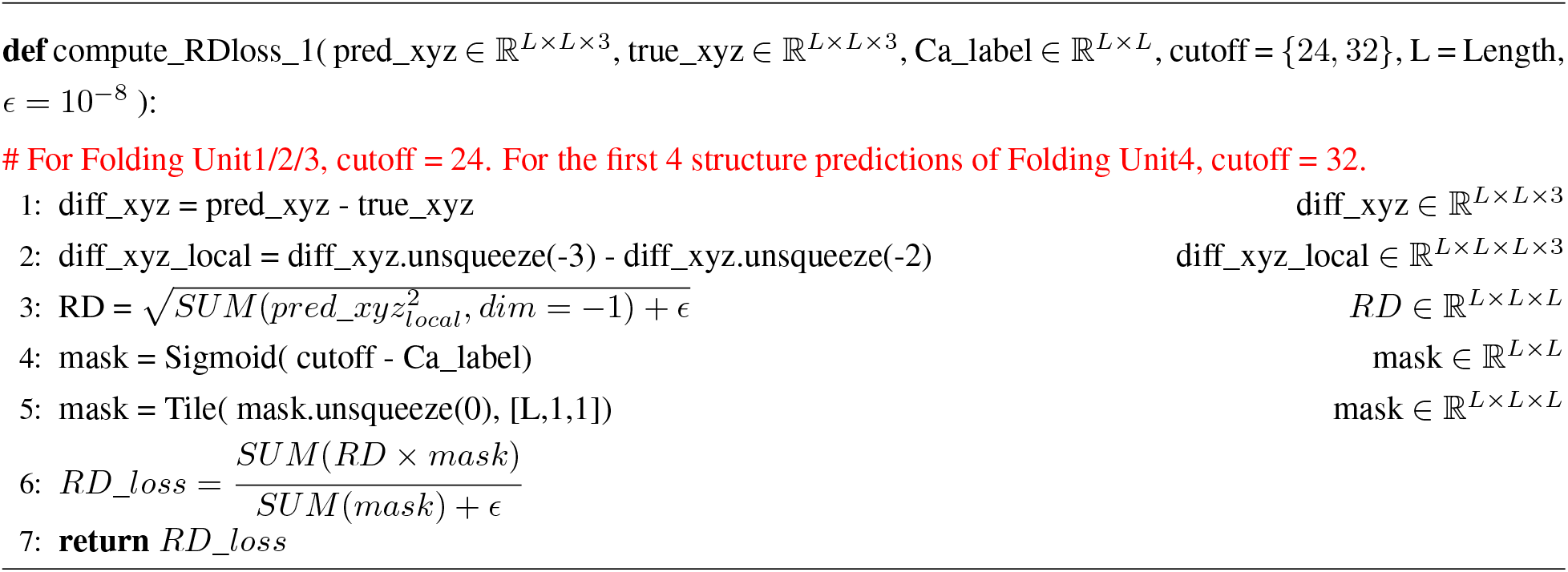

##### Algorithm 12 compute_RDloss_2

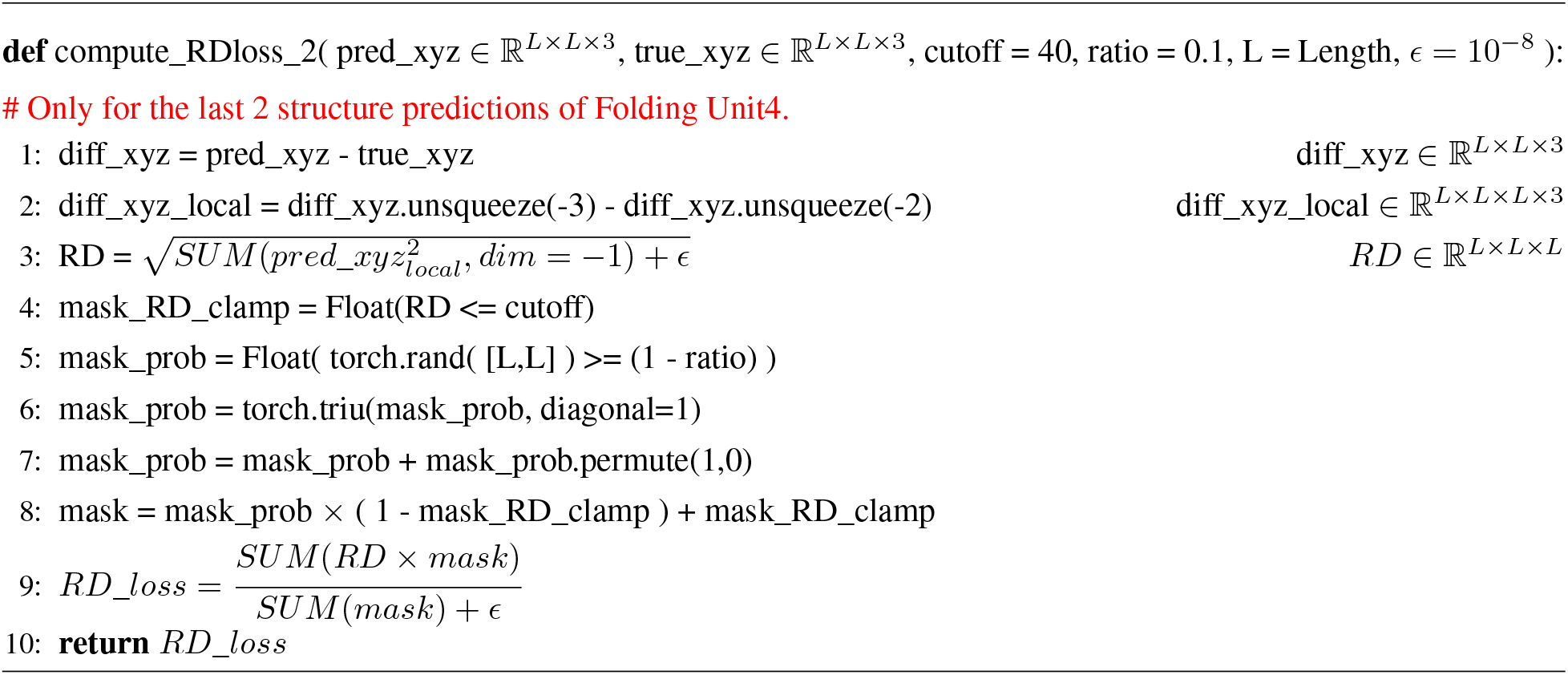

##### Algorithm 13 compute_CaDist_Loss

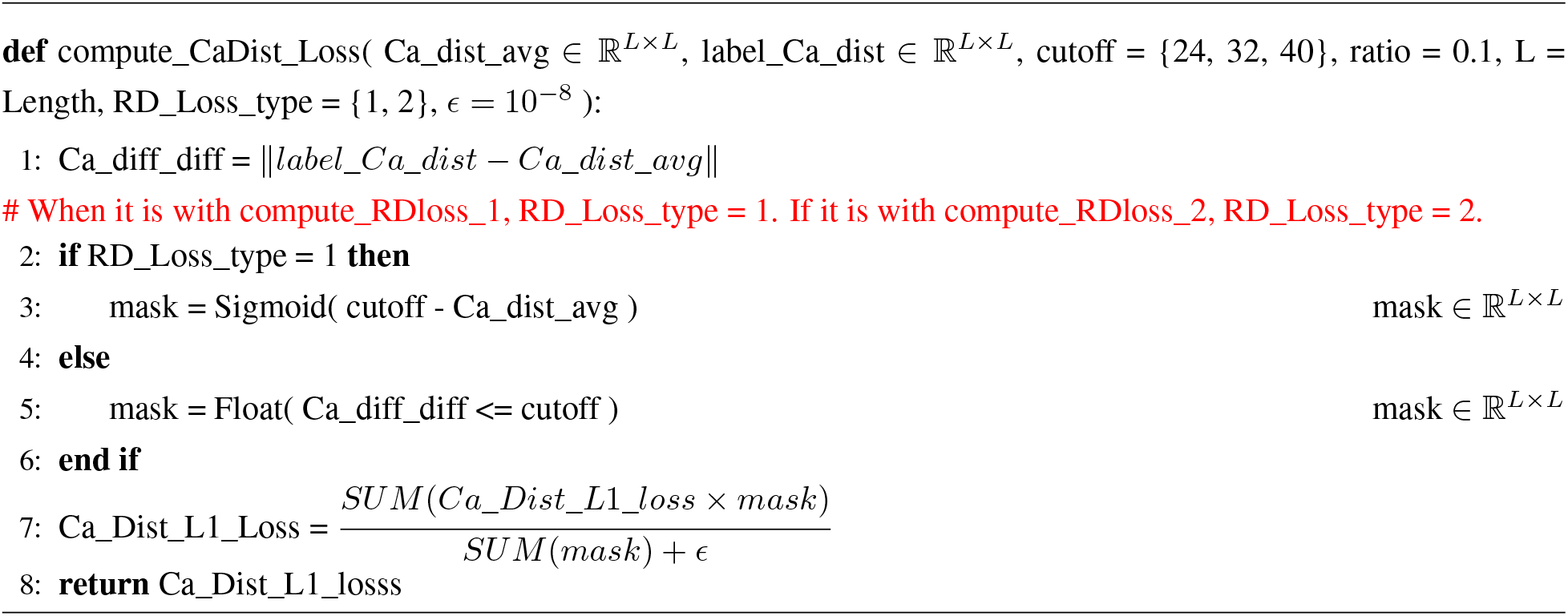

##### Algorithm 14 compute_pLDDT_Loss

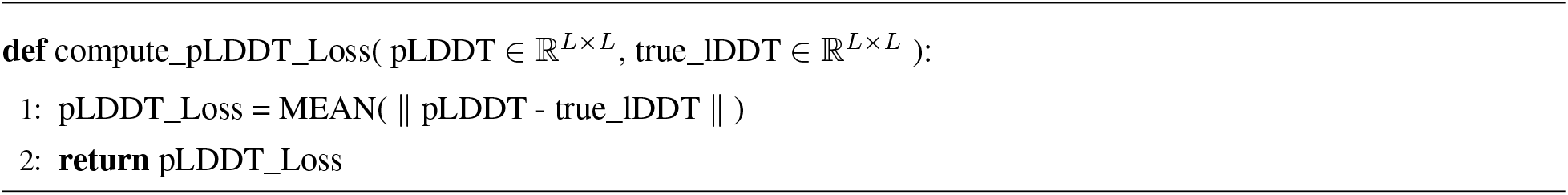

##### Algorithm 15 compute_Violation_Loss

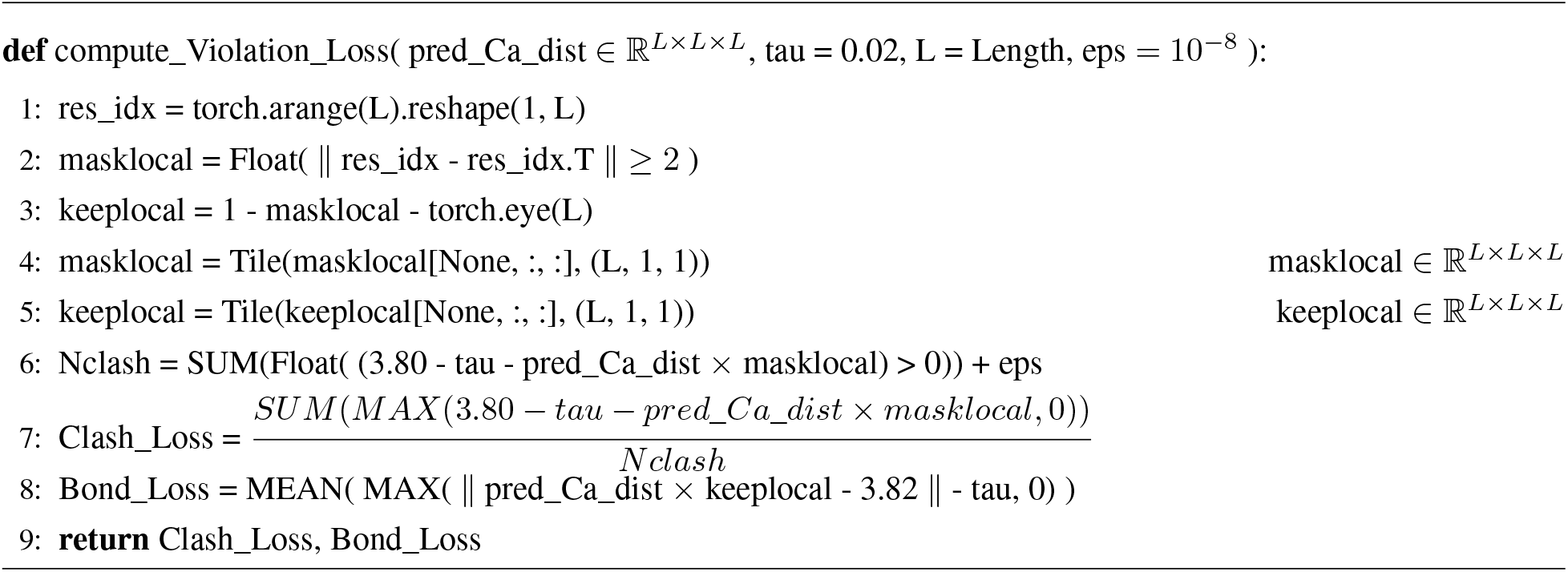

#### 4.3 Algorithm for the fitness and ΔΔG/ΔTm prediction modules in SPIRED-Fitness and SPIRED-Stab

##### Algorithm 16 Fitness_Module

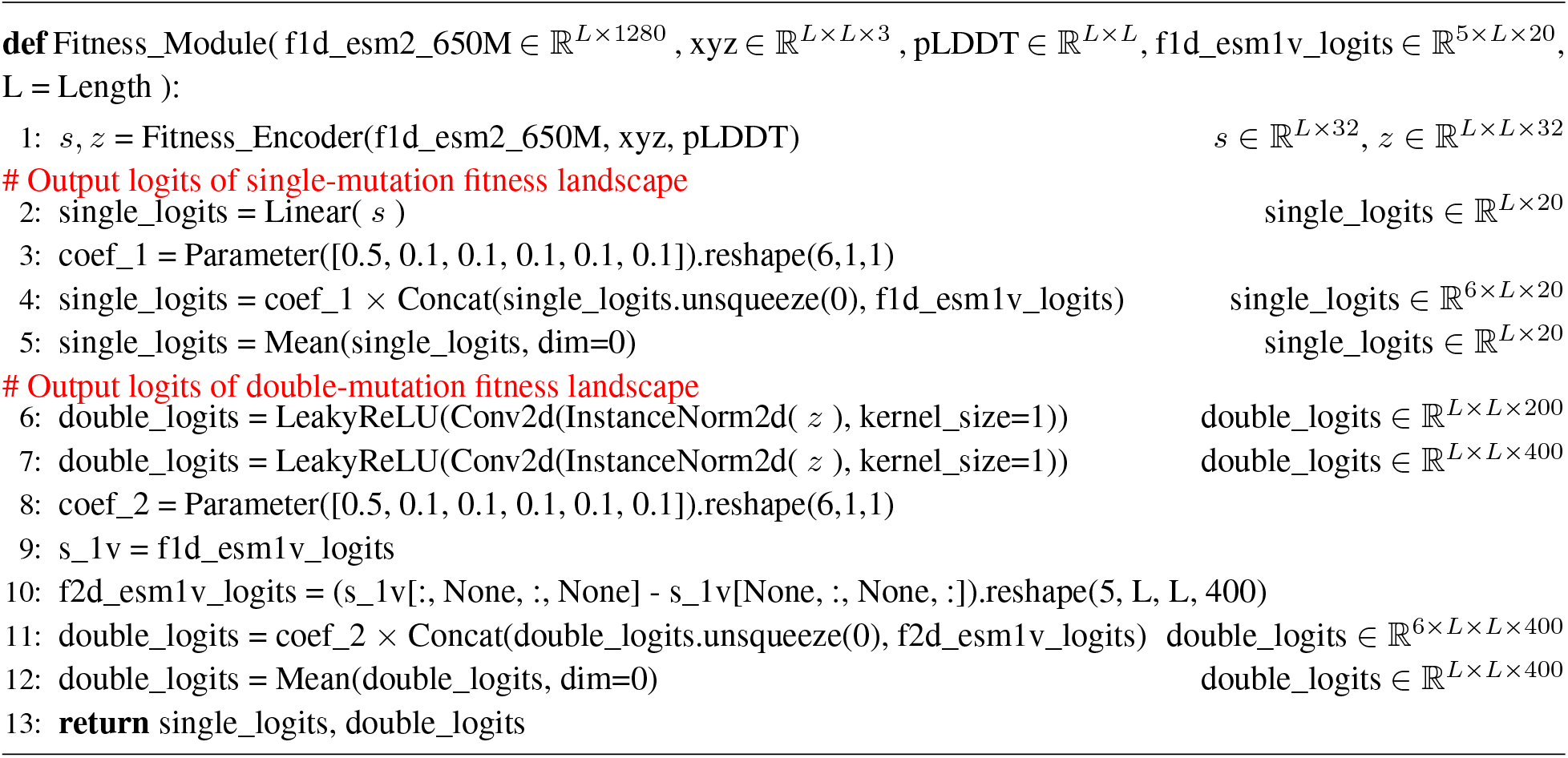

##### Algorithm 17 Fitness_Encoder

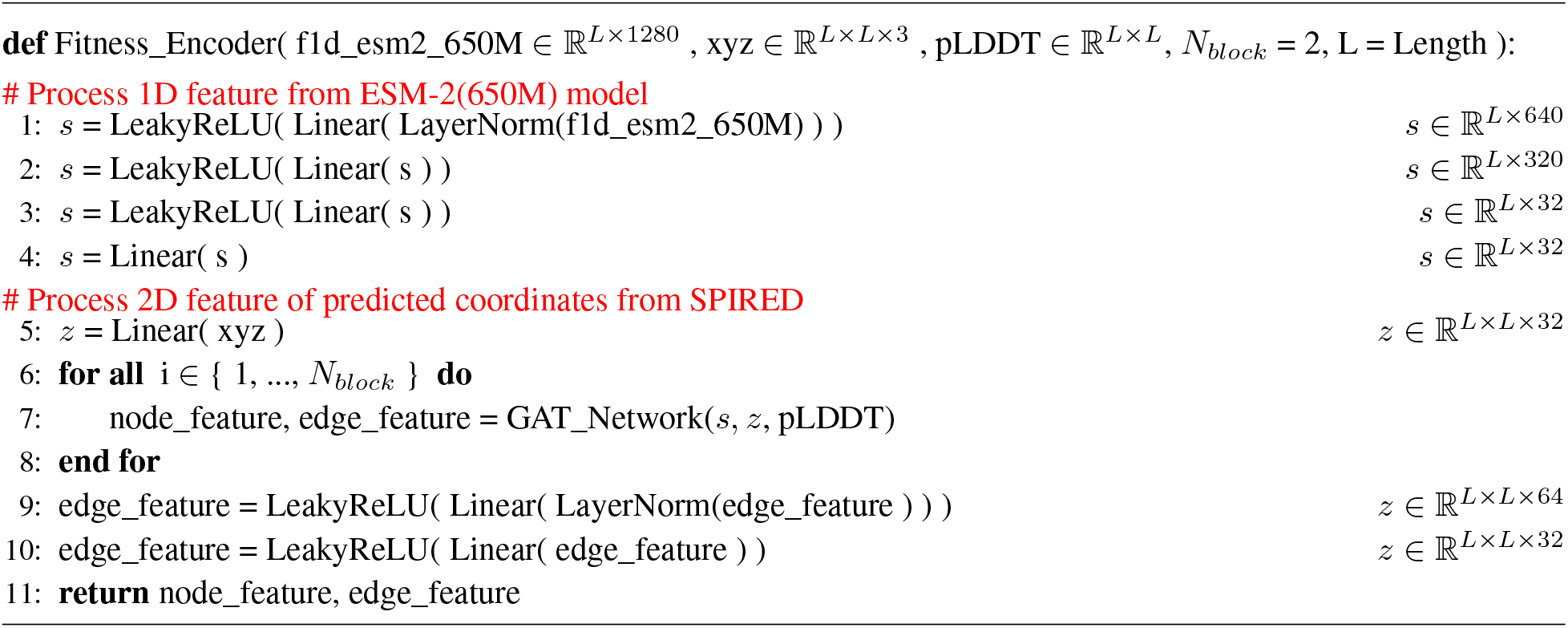

##### Algorithm 18 GAT_Network

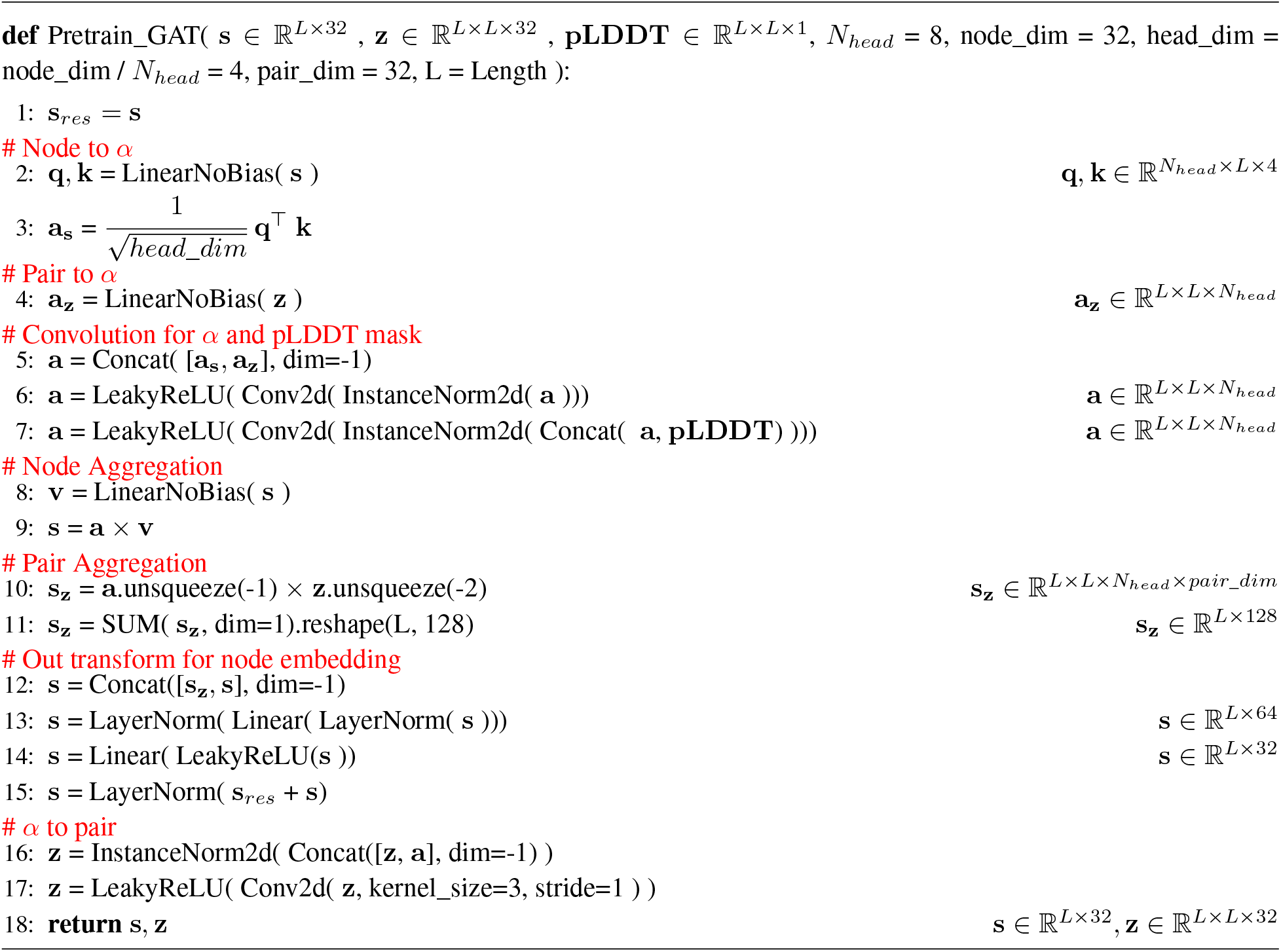

##### Algorithm 19 Stability_Pred_Module

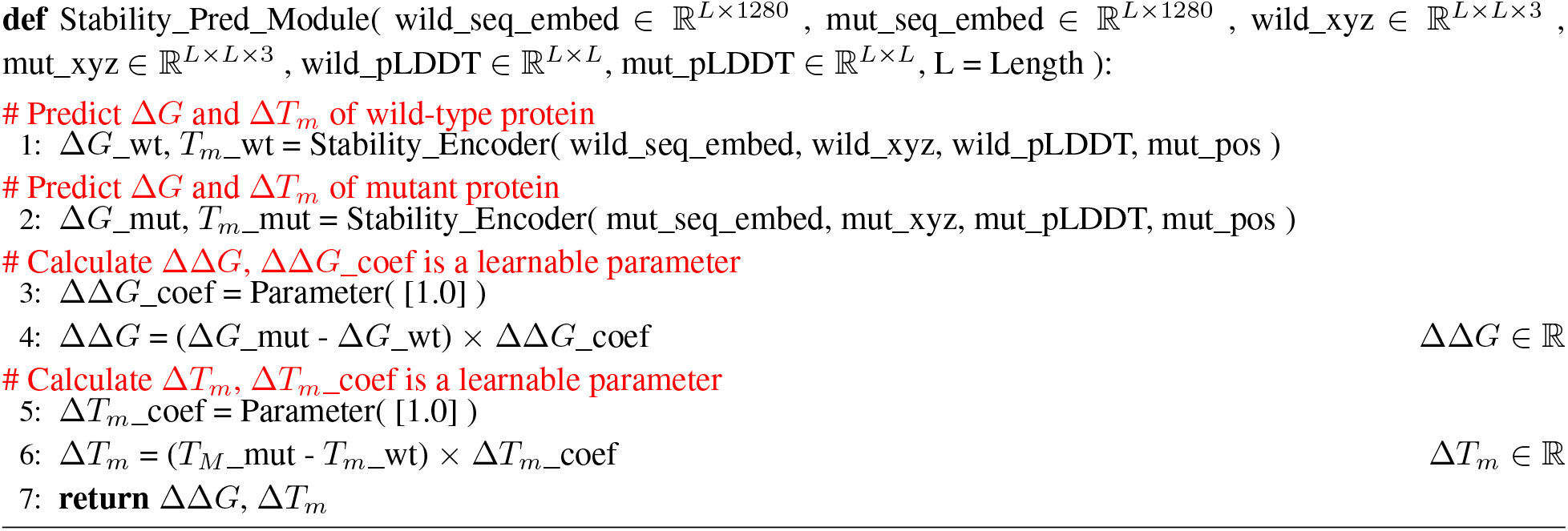

The mut_pos is a tensor of length *L*, used to represent the position of the mutated residue. It has a value of 1 at the index corresponding to the mutation position, and 0 at all other positions.

##### Algorithm 20 Stability_Encoder

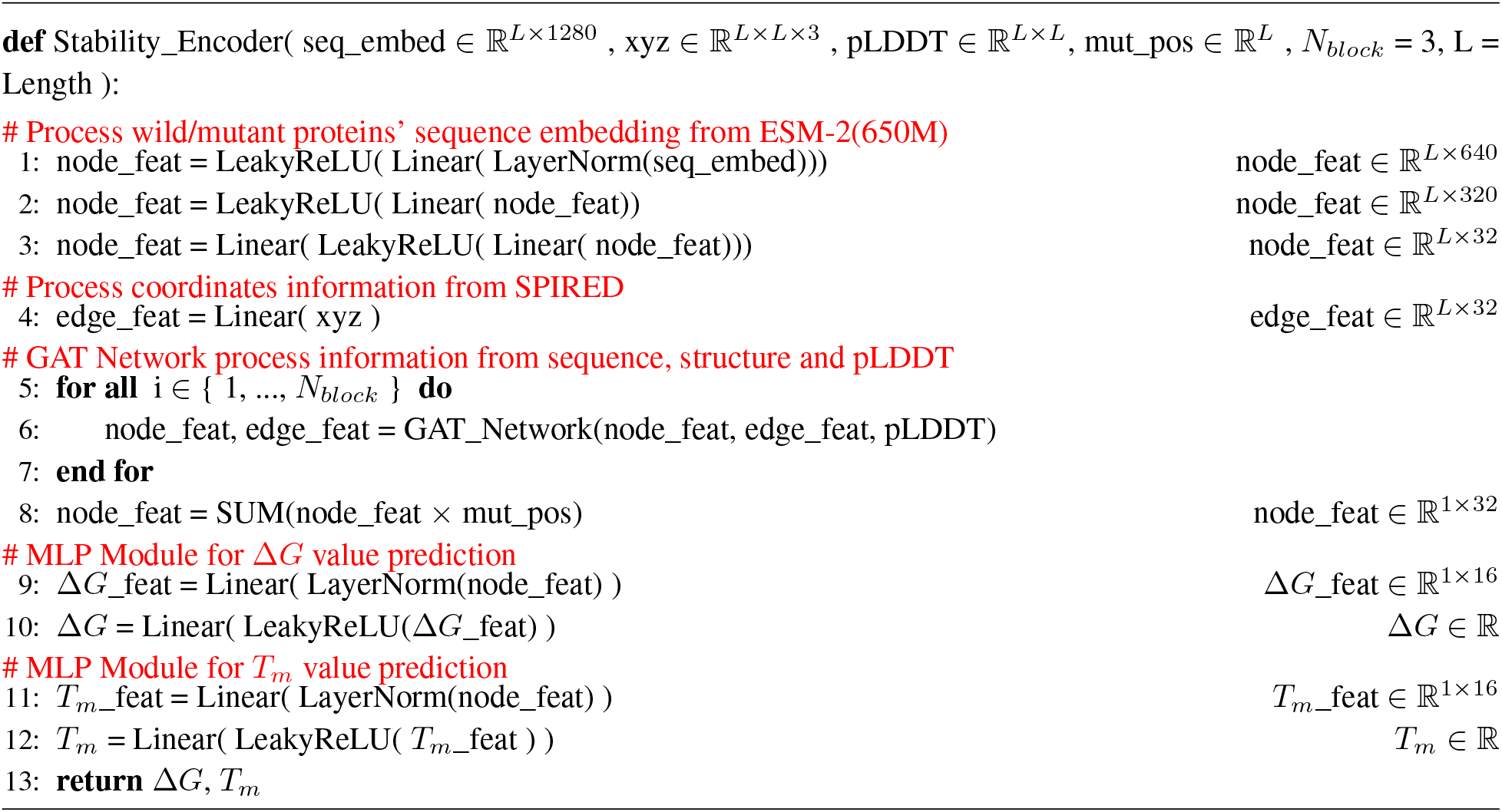

## References

1. Anfinsen, C. B. Principles that Govern the Folding of Protein Chains. Science 181, 223–230. ISSN: 1095-9203. 10.1126/science.181.4096.223 (1973).

2. Morcos, F., Pagnani, A., Lunt, B., et al. Direct-coupling analysis of residue coevolution captures native contacts across many protein families. Proceedings of the National Academy of Sciences 108, E1293–E1301. eprint: https://www.pnas.org/doi/pdf/10.1073/pnas.1111471108. https://www.pnas.org/doi/abs/10.1073/pnas.1111471108 (2011).

3. Marks, D. S., Hopf, T. A. & Sander, C. Protein structure prediction from sequence variation. Nature Biotechnology 30, 1072–1080. ISSN: 1546-1696. 10.1038/nbt.2419 (2012).

4. De Juan, D., Pazos, F. & Valencia, A. Emerging methods in protein co-evolution. Nature Reviews Genetics 14, 249–261. ISSN: 1471-0064. 10.1038/nrg3414 (2013).

5. Rives, A., Meier, J., Sercu, T., et al. Biological structure and function emerge from scaling unsupervised learning to 250 million protein sequences. Proceedings of the National Academy of Sciences 118. ISSN: 1091-6490. 10.1073/pnas.2016239118 (2021).

6. Rao, R., Liu, J., Verkuil, R., et al. MSA Transformer. bioRxiv. https://www.biorxiv.org/content/10.1101/2021.02.12.430858v1 (2021).

7. Jumper, J., Evans, R., Pritzel, A., et al. Highly accurate protein structure prediction with AlphaFold. Nature 596, 583–589. ISSN: 1476-4687. 10.1038/s41586-021-03819-2 (2021).

8. Baek, M., DiMaio, F., Anishchenko, I., et al. Accurate prediction of protein structures and interactions using a three-track neural network. Science 373, 871–876. eprint: https://www.science.org/doi/pdf/10.1126/science.abj8754. https://www.science.org/doi/abs/10.1126/science.abj8754 (2021).

9. Lin, Z., Akin, H., Rao, R., et al. Evolutionary-scale prediction of atomic-level protein structure with a language model. Science 379, 1123–1130. eprint: https://www.science.org/doi/pdf/10.1126/science.ade2574. https://www.science.org/doi/abs/10.1126/science.ade2574 (2023).

10. Wu, R., Ding, F., Wang, R., et al. High-resolution de novo structure prediction from primary sequence. bioRxiv. eprint: https://www.biorxiv.org/content/early/2022/07/22/2022.07.21.500999.full.pdf. https://www.biorxiv.org/content/early/2022/07/22/2022.07.21.500999 (2022).

11. Fang, X., Wang, F., Liu, L., et al. A method for multiple-sequence-alignment-free protein structure prediction using a protein language model. Nature Machine Intelligence 5, 1087–1096. ISSN: 2522-5839. 10.1038/s42256-023-00721-6 (2023).

12. Wang, W., Peng, Z. & Yang, J. Single-sequence protein structure prediction using supervised transformer protein language models. Nature Computational Science 2, 804–814. ISSN: 2662-8457. 10.1038/s43588-022-00373-3 (2022).

13. Chowdhury, R., Bouatta, N., Biswas, S., et al. Single-sequence protein structure prediction using a language model and deep learning. Nature Biotechnology 40, 1617–1623. ISSN: 1546-1696. 10.1038/s41587-022-01432-w (2022).

14. Akdel, M., Pires, D. E. V., Pardo, E. P., et al. A structural biology community assessment of AlphaFold2 applications. Nature Structural & Molecular Biology 29, 1056–1067. ISSN: 1545-9985. 10.1038/s41594-022-00849-w (2022).

15. Mansoor, S., Baek, M., Juergens, D., et al. Zero-shot mutation effect prediction on protein stability and function using RoseTTAFold. Protein Science 32 (2023).

16. Romero, P. A. & Arnold, F. H. Exploring protein fitness landscapes by directed evolution. Nature Reviews Molecular Cell Biology 10, 866–876. ISSN: 1471-0080. 10.1038/nrm2805 (2009).

17. Li, M., Kang, L., Xiong, Y., et al. SESNet: sequence-structure feature-integrated deep learning method for data-efficient protein engineering. Journal of Cheminformatics 15. 10.1186/s13321-023-00688-x (2023).

18. Chen, Y., Lu, H., Zhang, N., et al. PremPS: Predicting the impact of missense mutations on protein stability. PLOS Computational Biology 16 (ed Keskin, O.) e1008543. 10.1371/journal.pcbi.1008543 (2020).

19. Zhou, Y., Pan, Q., Pires, D. E. V., et al. DDMut: predicting effects of mutations on protein stability using deep learning. Nucleic Acids Research 51, W122–W128. ISSN: 1362-4962. 10.1093/nar/gkad472 (2023).

20. Tunyasuvunakool, K., Adler, J., Wu, Z., et al. Highly accurate protein structure prediction for the human proteome. Nature 596, 590–596. ISSN: 1476-4687. 10.1038/s41586-021-03828-1 (2021).

21. Varadi, M., Anyango, S., Deshpande, M., et al. AlphaFold Protein Structure Database: massively expanding the structural coverage of protein-sequence space with high-accuracy models. Nucleic Acids Research 50, D439–D444. ISSN: 0305-1048. eprint: https://academic.oup.com/nar/article-pdf/50/D1/D439/43502749/gkab1061.pdf. 10.1093/nar/gkab1061 (2021).

22. Yang, Z., Zeng, X., Zhao, Y., et al. AlphaFold2 and its applications in the fields of biology and medicine. Signal Transduction and Targeted Therapy 8. ISSN: 2059-3635. 10.1038/s41392-023-01381-z (2023).

23. Xu, Y., Liu, D. & Gong, H. Improving the prediction of protein stability changes upon mutations by geometric learning and a pre-training strategy. bioRxiv. eprint: https://www.biorxiv.org/content/early/2023/05/30/2023.05.28.542668.full.pdf. https://www.biorxiv.org/content/early/2023/05/30/2023.05.28.542668 (2023).

24. Haas, J., Barbato, A., Behringer, D., et al. Continuous Automated Model EvaluatiOn (CAMEO) complementing the critical assessment of structure prediction in CASP12. Proteins: Structure, Function, and Bioinformatics 86, 387–398. ISSN: 1097-0134. 10.1002/prot.25431 (2017).

25. Robin, X., Haas, J., Gumienny, R., et al. Continuous Automated Model EvaluatiOn (CAMEO)—Perspectives on the future of fully automated evaluation of structure prediction methods. Proteins: Structure, Function, and Bioinformatics 89, 1977–1986. ISSN: 1097-0134. 10.1002/prot.26213 (2021).

26. Alexander, L. T., Durairaj, J., Kryshtafovych, A., et al. Protein target highlights in CASP15: Analysis of models by structure providers. Proteins: Structure, Function, and Bioinformatics 91, 1571–1599. ISSN: 1097-0134. 10.1002/prot.26545 (2023).

27. Berman, H. M. The Protein Data Bank. Nucleic Acids Research 28, 235–242. ISSN: 1362-4962. 10.1093/nar/28.1.235 (2000).

28. Fowler, D. M. & Fields, S. Deep mutational scanning: a new style of protein science. Nature Methods 11, 801–807. ISSN: 1548-7105. 10.1038/nmeth.3027 (2014).

29. Ahdritz, G., Bouatta, N., Floristean, C., et al. OpenFold: Retraining AlphaFold2 yields new insights into its learning mechanisms and capacity for generalization. bioRxiv. eprint: https://www.biorxiv.org/content/early/2022/11/22/2022.11.20.517210.full.pdf. https://www.biorxiv.org/content/10.1101/2022.11.20.517210 (2022).

30. Mao, W., Ding, W., Xing, Y., et al. AmoebaContact and GDFold as a pipeline for rapid de novo protein structure prediction. Nature Machine Intelligence 2, 25–33. ISSN: 2522-5839. 10.1038/s42256-019-0130-4 (2019).

31. Yang, J., Anishchenko, I., Park, H., et al. Improved protein structure prediction using predicted interresidue orientations. Proceedings of the National Academy of Sciences 117, 1496–1503. eprint: https://www.pnas.org/doi/pdf/10.1073/pnas.1914677117. https://www.pnas.org/doi/abs/10.1073/pnas.1914677117 (2020).

32. Meier, J., Rao, R., Verkuil, R., et al. Language models enable zero-shot prediction of the effects of mutations on protein function. bioRxiv. eprint: https://www.biorxiv.org/content/early/2021/07/10/2021.07.09.450648.full.pdf. https://www.biorxiv.org/content/early/2021/07/10/2021.07.09.450648 (2021).

33. Steinegger, M. & Söding, J. MMseqs2 enables sensitive protein sequence searching for the analysis of massive data sets. Nature Biotechnology 35, 1026–1028. ISSN: 1546-1696. 10.1038/nbt.3988 (2017).

34. Sillitoe, I., Bordin, N., Dawson, N., et al. CATH: increased structural coverage of functional space. Nucleic Acids Research 49, D266–D273. eprint: https://academic.oup.com/nar/article-pdf/49/D1/D266/35364652/gkaa1079.pdf. 10.1093/nar/gkaa1079 (2020).

35. Kingma, D. P. & Ba, J. Adam: A Method for Stochastic Optimization 2017. arXiv: 1412.6980 [cs.LG].

36. Chandonia, J.-M., Guan, L., Lin, S., et al. SCOPe: improvements to the structural classification of proteins – extended database to facilitate variant interpretation and machine learning. Nucleic Acids Research 50, D553–D559. ISSN: 1362-4962. 10.1093/nar/gkab1054 (2021).

37. Tsuboyama, K., Dauparas, J., Chen, J., et al. Mega-scale experimental analysis of protein folding stability in biology and design. Nature 620, 434–444 (2023).

38. Esposito, D., Weile, J., Shendure, J., et al. MaveDB: an open-source platform to distribute and interpret data from multiplexed assays of variant effect. Genome Biology 20. ISSN: 1474-760X. 10.1186/s13059-019-1845-6 (2019).

39. Rubin, A. F., Min, J. K., Rollins, N. J., et al. MaveDB v2: a curated community database with over three million variant effects from multiplexed functional assays. bioRxiv. 10.1101/2021.11.29.470445 (2021).

40. Riesselman, A. J., Ingraham, J. B. & Marks, D. S. Deep generative models of genetic variation capture the effects of mutations. Nature Methods 15, 816–822. 10.1038/s41592-018-0138-4 (2018).

41. Blondel, M., Teboul, O., Berthet, Q., et al. Fast Differentiable Sorting and Ranking in INTERNATIONAL CONFERENCE ON MACHINE LEARNING, VOL 119 (eds Daume, H. & Singh, A.) 119. International Conference on Machine Learning (ICML), ELECTR NETWORK, JUL 13-18, 2020 (2020).

42. Nikam, R., Kulandaisamy, A., Harini, K., et al. ProThermDB: thermodynamic database for proteins and mutants revisited after 15 years. Nucleic Acids Research 49, D420–D424. 10.1093/nar/gkaa1035 (2020).

43. Xavier, J. S., Nguyen, T.-B., Karmarkar, M., et al. ThermoMutDB: a thermodynamic database for missense mutations. Nucleic Acids Research 49, D475–D479. 10.1093/nar/gkaa925 (2020).

44. Pancotti, C., Benevenuta, S., Birolo, G., et al. Predicting protein stability changes upon single-point mutation: a thorough comparison of the available tools on a new dataset. Briefings in Bioinformatics 23. 10.1093/bib/bbab555 (2022).

45. Hernández, I. M., Dehouck, Y., Bastolla, U., et al. Predicting protein stability changes upon mutation using a simple orientational potential. Bioinformatics 39, btad011. ISSN: 1367-4811. eprint: https://academic.oup.com/bioinformatics/article-pdf/39/1/btad011/48769997/btad011.pdf. 10.1093/bioinformatics/btad011 (2023).

46. Zhang, Y. & Skolnick, J. Scoring function for automated assessment of protein structure template quality. Proteins: Structure, Function, and Bioinformatics 57, 702–710. ISSN: 1097-0134. 10.1002/prot.20264 (2004).

47. Mariani, V., Biasini, M., Barbato, A., et al. lDDT: a local superposition-free score for comparing protein structures and models using distance difference tests. Bioinformatics 29, 2722–2728. ISSN: 1367-4803. eprint: https://academic.oup.com/bioinformatics/article-pdf/29/21/2722/50746741/bioinformatics\_29\_21\_2722.pdf. 10.1093/bioinformatics/btt473 (2013).

48. Luo, Y., Jiang, G., Yu, T., et al. ECNet is an evolutionary context-integrated deep learning framework for protein engineering. Nature Communications 12 (2021).

49. Wang, J., Lisanza, S., Juergens, D., et al. Scaffolding protein functional sites using deep learning. Science 377, 387–394. 10.1126/science.abn2100 (2022).

50. Mansoor, S., Baek, M., Juergens, D., et al. Accurate Mutation Effect Prediction using RoseTTAFold. bioRxiv. 10.1101/2022.11.04.515218 (2022).

51. Rives, A., Meier, J., Sercu, T., et al. Biological structure and function emerge from scaling unsupervised learning to 250 million protein sequences. Proceedings of the National Academy of Sciences 118. 10.1073/pnas.2016239118 (2021).

52. Ouyang-Zhang, J., Diaz, D. J., Klivans, A., et al. Predicting a Protein’s Stability under a Million Mutations. NeurIPS (2023).

53. Worth, C. L., Preissner, R. & Blundell, T. L. SDM–a server for predicting effects of mutations on protein stability and malfunction. Nucleic Acids Research 39, W215–W222. 10.1093/nar/gkr363 (2011).

54. Li, B., Yang, Y. T., Capra, J. A., et al. Predicting changes in protein thermodynamic stability upon point mutation with deep 3D convolutional neural networks. PLOS Computational Biology 16 (ed Fariselli, P.) e1008291. 10.1371/journal.pcbi.1008291 (2020).

55. Pires, D. E. V., Ascher, D. B. & Blundell, T. L. mCSM: predicting the effects of mutations in proteins using graph-based signatures. Bioinformatics 30, 335–342. 10.1093/bioinformatics/btt691 (2013).

56. Rodrigues, C. H., Pires, D. E. & Ascher, D. B. DynaMut: predicting the impact of mutations on protein conformation, flexibility and stability. Nucleic Acids Research 46, W350–W355. 10.1093/nar/gky300 (2018).

57. Rodrigues, C. H., Pires, D. E. & Ascher, D. B. DynaMut2: Assessing changes in stability and flexibility upon single and multiple point missense mutations. Protein Science 30, 60–69. ISSN: 1469-896X. 10.1002/pro.3942 (2020).

58. Capriotti, E., Fariselli, P. & Casadio, R. I-Mutant2.0: predicting stability changes upon mutation from the protein sequence or structure. Nucleic Acids Research 33, W306–W310. 10.1093/nar/gki375 (2005).

59. Laimer, J., Hofer, H., Fritz, M., et al. MAESTRO multi agent stability prediction upon point mutations. BMC Bioinformatics 16. ISSN: 1471-2105. 10.1186/s12859-015-0548-6 (2015).

60. Pires, D. E. V., Ascher, D. B. & Blundell, T. L. DUET: a server for predicting effects of mutations on protein stability using an integrated computational approach. Nucleic Acids Research 42, W314–W319. 10.1093/nar/gku411 (2014).

61. Benevenuta, S., Pancotti, C., Fariselli, P., et al. An antisymmetric neural network to predict free energy changes in protein variants. Journal of Physics D: Applied Physics 54, 245403. 10.1088/1361-463/abedfb (2021).

62. Schymkowitz, J., Borg, J., Stricher, F., et al. The FoldX web server: an online force field. Nucleic Acids Research 33, W382–W388. 10.1093/nar/gki387 (2005).

63. Dehouck, Y., Kwasigroch, J. M., Gilis, D., et al. PoPMuSiC 2.1: a web server for the estimation of protein stability changes upon mutation and sequence optimality. BMC bioinformatics 12, 1–12 (2011).

64. Montanucci, L., Capriotti, E., Frank, Y., et al. DDGun: an untrained method for the prediction of protein stability changes upon single and multiple point variations. BMC Bioinformatics 20. 10.1186/s12859-019-2923-1 (2019).

65. Fariselli, P., Martelli, P. L., Savojardo, C., et al. INPS: predicting the impact of non-synonymous variations on protein stability from sequence. Bioinformatics 31, 2816–2821. 10.1093/bioinformatics/btv291 (2015).

66. Cheng, J., Randall, A. & Baldi, P. Prediction of protein stability changes for single-site mutations using support vector machines. Proteins: Structure, Function, and Bioinformatics 62, 1125–1132. ISSN: 1097-0134. 10.1002/prot.20810 (2005).

67. Li, G., Panday, S. K. & Alexov, E. SAAFEC-SEQ: A Sequence-Based Method for Predicting the Effect of Single Point Mutations on Protein Thermodynamic Stability. International Journal of Molecular Sciences 22, 606. 10.3390/ijms22020606 (2021).

68. Masso, M. & Vaisman, I. I. Accurate prediction of stability changes in protein mutants by combining machine learning with structure based computational mutagenesis. Bioinformatics 24, 2002–2009. 10.1093/bioinformatics/btn353 (2008).

69. Masso, M. & Vaisman, I. I. AUTO-MUTE 2.0: A Portable Framework with Enhanced Capabilities for Predicting Protein Functional Consequences upon Mutation. Advances in Bioinformatics 2014, 1–7. 10.1155/2014/278385 (2014).

70. Pucci, F., Bourgeas, R. & Rooman, M. Predicting protein thermal stability changes upon point mutations using statistical potentials: Introducing HoTMuSiC. Scientific Reports 6. 10.1038/srep23257 (2016).

71. Paszke, A., Gross, S., Massa, F., et al. in Proceedings of the 33rd International Conference on Neural Information Processing Systems (Curran Associates Inc., Red Hook, NY, USA, 2019).

